# UDP-glucose Activation of a G-protein/Sucrose Synthase Signaling Supercomplex

**DOI:** 10.1101/2025.09.10.672744

**Authors:** Haiyan Jia, Fei Lou, Justin Watkins, Celio Cabral Oliveira, Christopher G. Tate, Alan M. Jones

**Author notes:** Correspondence: Haiyan Jia.

## Abstract

*Arabidopsis thaliana* REGULATOR OF G PROTEIN SIGNALING 1 (AtRGS1, hereafter, RGS1) within the heterotrimeric G protein complex plays a role in sensing extracellular signals. High millimolar concentrations of some sugars induce RGS1 endocytosis and therefore G protein activation. However, this is not likely a direct, rather an indirect effect through a sugar metabolite. One metabolite at the plasma membrane where RGS1 resides is UDP-glucose (UDPG) produced by SUCROSE SYNTHASE (SUS) enzymes, previously shown to interact with RGS1. Our direct measurements indicate that the K_d_ for UDPG binding to RGS1 (2.4 µM) is consistent with UDPG activation of RGS1 with an EC_50_ of 17 µM to rapidly trigger G protein activation within minutes. RGS1 associates directly with SUS1 and SUS4, exhibiting affinities (K_d_ = 161 nM and 106 nM, respectively) that are markedly tighter than the corresponding K_m_ values for UDPG formation (40 µM and 70 µM, respectively). Therefore, SUS may be one source for this RGS1 ligand in a manner analogous to “metabolite funneling” in enzymology. UDPG induces rapid phosphorylation of RGS1 at the C-terminal phosphorylation cluster, a site previously implicated in RGS1 endocytosis and G protein activation. UDPG activation of RGS1 changes the global transcript profiles in various processes, largely involving plant defense. These results provide the first molecular evidence for UDPG as a newly recognized low-molecular weight signal controlling key aspects of plant physiology.

## INTRODUCTION

Pathogens, such as bacteria, cause a host plant to release sugars to the apoplast in order to favor their growth (Doidy et al., 2012). Microbe-Associated Molecular Patterns (MAMPs), such as flg22 released from bacterial flagella, are sensed by the host receptor kinase FLS2/BAK1 and defense responses are initiated. These responses include an increase in activity of cell wall invertases (INV) to convert apoplastic sucrose to monosaccharides, and in phosphorylation of the monosaccharide transporter STP13 leading to increased activity to salvage those monosaccharides away from the pathogens (Yamada et al., 2016). As the sugar economy changes (Bolouri Moghaddam and Van den Ende, 2012; Breia et al., 2021; Chen et al., 2023; Yamada and Mine, 2024; Zhang et al., 2025b), the host cell balances the use of this carbon between growth and defense against the pathogen (Belkhadir et al., 2014). As such, sensing apoplastic sugars, i.e. essentially monitoring the host’s nutritional state, is a critical component of host defense strategy.

As for coupling an apoplastic signal to growth and defense, the heterotrimeric G protein complex, composed of Gα, Gβ, and Gγ subunits, works with plasma membrane receptors to affect concomitant nuclear changes (Delgado-Cerezo et al., 2012; Karimian et al., 2024; Lee et al., 2013; Liang et al., 2016; Liang et al., 2018; Liu et al., 2013; Trusov et al., 2006; Trusov et al., 2009; Urano et al., 2013; Zhang et al., 2025a; Zhang et al., 2008; Zhang et al., 2025c; Zhong et al., 2019). For example, phosphorylation-activated heterotrimeric G-protein signaling stabilizes nuclear proteins TCP14 and JAZ3, thereby repressing jasmonate signaling and enhancing plant immunity against biotrophic pathogens (Jia et al., 2025). Phosphorylation-mediated regulation of heterotrimeric G proteins plays diverse roles in plant cellular signaling (Kai Chua and Urano, 2025; Ma et al., 2024). Arabidopsis REGULATOR OF G PROTEIN SIGNALING 1 (RGS1), the prototype of a receptor-like family of seven-transmembrane (7TM) proteins with a cytoplasmic RGS domain (Chen et al., 2003), modulates self-activating heterotrimeric G proteins at the plasma membrane in plants, protists, and fungi (Urano et al., 2012). In plants, RGS1 uniquely integrates receptor-like sensing with GTPase-activating protein activity to couple external stimuli to G protein signaling and accelerate Gα inactivation. Apoplastic sugars (Chen and Jones, 2004; Tunc-Ozdemir et al., 2018; Urano et al., 2012) as well as flg22, induces rapid internalization of RGS1 in plant cells (Watkins et al., 2021), analogous to ligand-induced endocytosis of G protein coupled receptors (GPCRs) in animals (Rajagopal and Shenoy, 2018) except resulting in de-repression of G protein signaling rather than activation as in animals. However, direct physical evidence of sugar binding to RGS1 is limited. Wang et al. concluded that D-glucose at a high concentration (> 41.6 mM) specifically binds to tomato RGS1 in cell-based biolayer interferometry assays using leaf protoplasts (Wang et al., 2022). However, this low affinity of glucose binding to tomato RGS1 is at or near thermal noise and thus is physiologically meaningless. The lack of evidence for sugar binding to RGS1 and the high concentration of sugars, e.g. >100 mM (Chen and Jones, 2004; Fu et al., 2014), required for RGS1 to internalize is inconsistent with a specific orthologous binding site on RGS1. As an alternative, it is plausible that a low-abundant, common sugar metabolite from D-glucose and other photosynthetically-fixed sugars (Tunc-Ozdemir et al., 2018) is the endogenous RGS1 ligand (Urano et al., 2012).

Considerable sugar metabolism occurs on or near the plasma membrane (Amor et al., 1995; Carlson and Chourey, 1996; Stein and Granot, 2019) even in the absence of MAMPs. One common metabolite for various sugar substrates at the plasma membrane is UDP-glucose (UDPG). As shown in Figure 1A, UDPG is produced by sucrose synthases (in Arabidopsis, SUS1 to SUS6, phylogeny shown in supplemental Figure 1 (Baud et al., 2004)) that are localized in the cytosol, cell wall, and plasma membrane (Stein and Granot, 2019). UDPG is formed directly from UDP and sucrose and formed indirectly from several other sugars via enzymatic conversion. As illustrated in Figure 1A, the enzymatic network is complex. Sucrose is cleaved to D-glucose and fructose via INVs localized in the cytosol, vacuole, and cell wall (Stein and Granot, 2019). Monosaccharides such as D-glucose and fructose are rapidly converted to glucose-6-phosphate (G6P) (Moore et al., 2003) by hexokinase (HXK) and fructokinase (FK), then converted to glucose-1-phosphate (G1P) by phosphoglucomutase (PMG) (Doello and Forchhammer, 2023), which in turn enters the UDPG pool via UDPG pyrophosphorylase (UGPase) and UDP-sugar pyrophosphorylase (USPase) (Janse van Rensburg and Van den Ende, 2017). UDPG is metabolized by UDP-*N*-acetylglucosamine pyrophosphorylase 1 (UAP1) (Janse van Rensburg and Van den Ende, 2017), sucrose phosphate synthase (SPS) (Janse van Rensburg and Van den Ende, 2017), and trehalose-6-P synthase (TPS) (Ponnu et al., 2011) to trehalose-6-P (T6P). T6P has been reported as a signaling molecule regulating plant growth and defense responses (Gobel and Fichtner, 2023; Paul et al., 2018).

**Figure 1.**
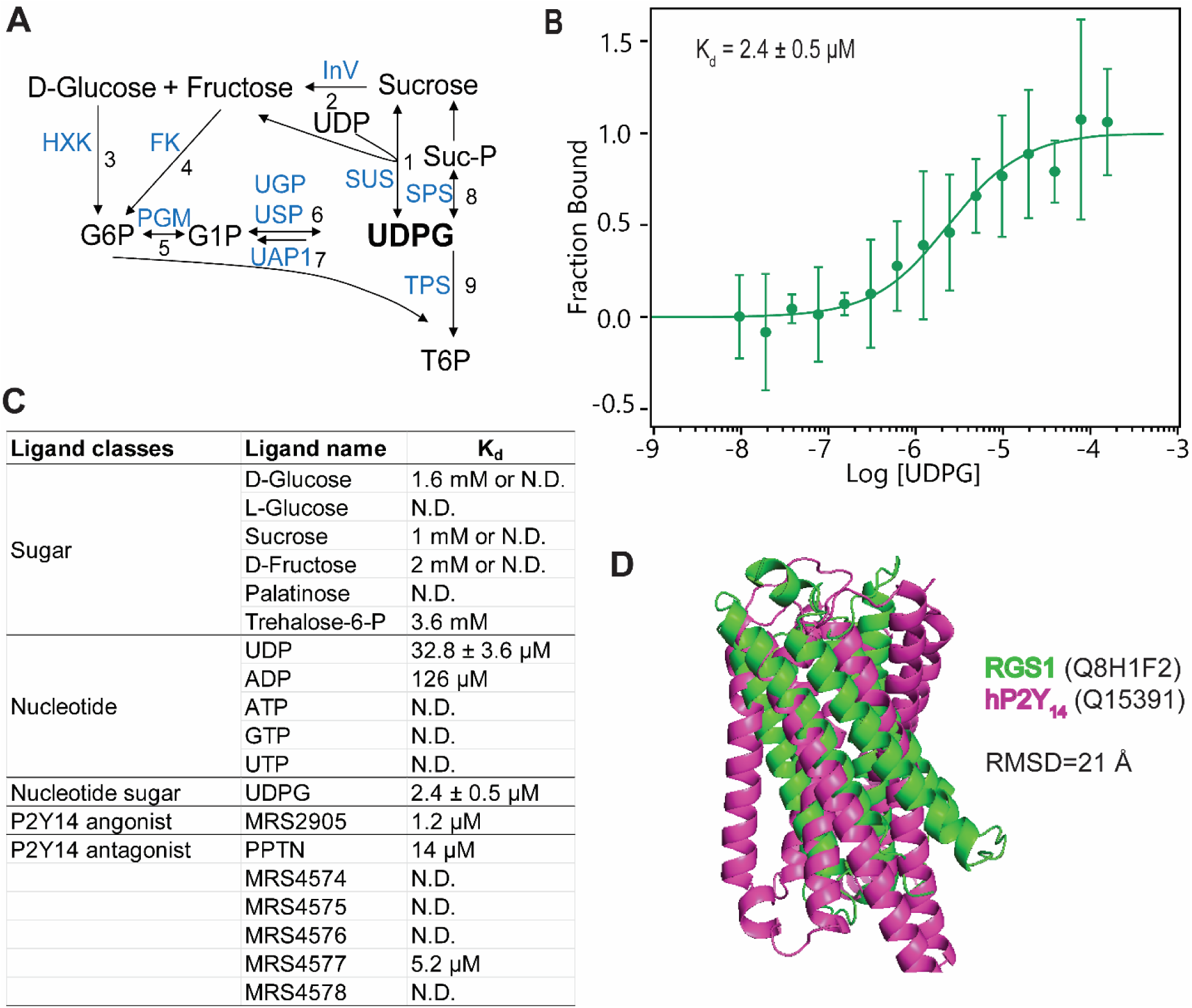
UDPG is a candidate ligand to RGS1 receptor. **(A)** Summary of sugar metabolism at the plasma membrane. One common metabolite for various sugars at the plasma membrane is UDPG. UDPG is produced from UDP and sucrose by sucrose synthases (SUS1 to SUS6) that are localized in the cytosol, cell wall, and at the plasma membrane (see **1** in panel). Sucrose is metabolized to D-glucose and fructose by invertases (INV) localized in the cytosol, vacuole, and cell wall (**2**). Monosaccharides such as D-glucose and fructose are rapidly converted to glucose-6-phosphate (G6P) (**3** and **4**) by hexokinase (HXK) and fructokinase (FK), and then into glucose-1-phosphate (G1P) by phosphoglucomutase (PMG) (**5**), which is in turn can enter the UDP-glucose pool via UDP-glucose pyrophosphorylase (UGPase) and UDP-sugar pyrophosphorylase (USPase) (**6**). UDPG is metabolized by UDP-*N*-acetylglucosamine pyrophosphorylase 1 (UAP1) (**7**), sucrose phosphate synthase (SPS) (8), and trehalose-6-P synthase (TPS) (**9**) to trehalose-6-P (T6P). **(B)** Quantification by MST of UDPG binding to full-length recombinant GFP-RGS1 produced by cell-free *in vitro* translation as described in Methods. Binding affinities were obtained by fitting MST data (n = 5) to a one-site binding model using MO Affinity Analysis software (NanoTemper Technologies). K_d_ values are reported as best-fit ± SE from nonlinear regression. Goodness-of-fit metrics are provided in supplemental Figure 2C. **(C)** Quantification of ligand binding to recombinant GFP–RGS1 by MST. K_d_ values in the millimolar range approach the thermal noise limit and are near the detection threshold. For UDP, data are averages of three independent replicates ± SD. For UDPG, K_d_ values are reported as best-fit ± SE from nonlinear regression. For low-affinity interactions, some measurements were below the detection limit and are indicated as “N.D.” (not detected). **(D)** Structural overlay of the modeled 7TM domains of RGS1 and human P2Y_14_. The modeled structure of the 7TM domain of RGS1 was determined as described in Methods. The 7TM structure models of RGS1 (green) and hP2Y_14_ (pink) were predicted using AlphaFold3 and superimposed using PyMOL. The overall RMSD (Å) between the two structures was calculated using PyMOL. Corresponding pLDDT confidence scores are shown in supplemental Figure 5.

SUS plays a profound role in plant stress adaptation by regulating carbon partitioning to support osmotic adjustment, energy metabolism, and cell wall biosynthesis (Stein and Granot, 2019). Under abiotic stresses such as drought, salinity, and cold, elevated SUS activity enhances sink strength and soluble sugar accumulation, contributing to cellular protection and stress resilience (Huang et al., 2025; Li et al., 2023a; Li et al., 2023b; Tian et al., 2025; Wang et al., 2025). These findings position SUS as a nexus of metabolic flux and stress-responsive signaling in plants.

Based on a yeast two-hybrid screen, RGS1 interacts with SUS1 and SUS4 and other sugar-metabolizing enzymes and inhibitors (Klopffleisch et al., 2011). This raises the possibility that there is a metabolite shunt to RGS1 to appraise the sugar economy for growth and defense strategies. While we do not rule out the possibility that RGS1 binds mono and disaccharides, it is difficult to comprehend and then reconcile how a single receptor has such a structurally-broad pharmaco-profile. It is also difficult to understand the high sugar dose required to elicit RGS1-mediated responses (Fu et al., 2014; Grigston et al., 2008).

Plants perceive extracellular signals to optimize growth vs. defense in a dynamically changing environment (Giolai and Laine, 2024; Monson et al., 2022; Wu et al., 2020). UDPG, a nucleotide sugar historically known for its role in glycosylation and cell wall biosynthesis, fairly recently emerged as a potent extracellular signaling molecule (Janse van Rensburg and Van den Ende, 2017). However, the molecular mechanisms underlying its perception and signaling remain unclear. Here, we show that extracellular UDPG activates heterotrimeric G protein signaling through direct interaction with the RGS1/SUS co-receptor complex, triggering broad transcriptional reprogramming. We posit that this response prioritizes local defense—via activation of pattern-triggered immunity (PTI) and jasmonate/ethylene signaling—while concurrently suppressing salicylic acid–mediated systemic acquired resistance and growth-related pathways. Our findings suggests that extracellular UDPG activates a G-protein/sucrose synthase signaling super-complex, modulating the defense–growth trade-off in plants.

## RESULTS AND DISCUSSION

### UDPG binds to RGS1 with physiologically-meaningful affinity

Many sugars activate G protein signaling by initializing the internalization of RGS1 (Chen and Jones, 2004), suggesting that a putative metabolic or transport step precedes RGS1 activation. For the reasons discussed above, one possible step by which different sugars induce RGS1 endocytosis is indirectly through formation of a metabolite in common. Our hypothesis is that UDPG is that metabolite.

As UDPG is our proposed direct ligand to RGS1, microscale thermophoresis (MST) (Jerabek-Willemsen et al., 2011) was used to quantitate the direct UDPG binding to recombinant GFP-RGS1 produced from cell free expression (Figure 1B; Supplemental Figures 2A and 2B). MST detects label-free, nanomolar to millimolar affinity interactions using a miniscule amount of protein that, otherwise, would be insufficient for all other approach-to-equilibrium binding assays (Jerabek-Willemsen et al., 2011). A dissociation constant of ∼2.4 µM K_d_ was determined (Figure 1B; Supplemental Figure 2C). The negative control GFP showed no interaction with UDPG (Supplemental Figure 4). We further expanded this approach to test possible binding of other sugars, nucleotides, nucleotide sugars and animal P2Y_14_ agonists and antagonists (Chambers et al., 2000; Freeman et al., 2001) to recombinant GFP-RGS1 (Figure 1C; Supplemental Figures 3 and 4). RGS1 bound nucleotides (e.g., UDP and ADP) and nucleotide sugar UDPG better than the tested monosaccharides and disaccharides (Figure 1C). The low affinity (∼mM K_d_ or too low to calculate, “N.D.”; Figure 1C) detected for other sugars is consistent with a lack of physiologically meaningful binding to RGS1 as reasoned above. In animals, UDP is a partial agonist and UDPG is a full agonist that binds with moderate affinity to the UDPG receptor, P2Y_14_, (Chambers et al., 2000; Freeman et al., 2001), therefore we tested the P2Y_14_ agonist MRS2905 and 6 P2Y_14_ antagonists (PPTN, MRS4574 to MRS4578, Supplemental Figure 3), and found MRS2905, PPTN and MRS4577 also bound recombinant GFP-RGS1 with a K_d_ ∼ 1-14 µM (Figure 1C; Supplemental Figure 4) while no binding was detected from the antagonists MRS4574, MRS4575, MRS4576, or MRS4578 (Figure 1C; Supplemental Figure 4). Despite this overlap in pharmacological profiles of RGS1 and P2Y_14_, we found no conspicuous structural homology between the modeled 7TM domain of RGS1 and the P2Y_14_ receptor as indicated by a high root-mean-square deviation (RMSD =21 Å) (Figure 1D; corresponding pLDDT confidence scores are shown in supplemental Figure 5). As a reference for comparison, the modeled 7TM domain of RGS1 exhibits relatively high structural similarity to animal class C GPCRs, with an RMSD of 5.9 Å upon superposition with PDB entry 4OR2 (Lou et al., 2024). This degree of similarity is greater than the similarity between the animal GPCR classes.

### UDPG, a novel activator of G-protein-coupled signaling

Multiple sugars activate G signaling in plants by inducing RGS1 endocytosis which, in turn, modulate signaling by de-repressing the self-activating canonical Gα subunit (Fu et al., 2014; Urano et al., 2012). This is a quantifiable response that is directly and linearly proportional to the degree of activated G signaling (Fu et al., 2014). Given that UDPG is a metabolite in common to many apoplastic sugars and produced from multiple reactions (Figure 1A), UDPG may be the physiological element in detecting photosynthetically-fixed sugars for G protein activation *in planta* (Tunc-Ozdemir et al., 2018). This rationale motivated our choice to test RGS1-YFP internalization as a function of dose and time. Five-d-old, etiolated seedlings expressing RGS1-YFP were treated with UDPG at a series of doses from nM to mM, and the subcellular localization of RGS1 at 20 min was observed (Figure 2A). UDPG induced RGS1 internalization with an EC_50_ of ∼17 µM (R^2^ = 0.54 in a nonlinear least square fit to dose response) and reached saturation at ∼123 µM UDPG within 20 mins (Figure 2B). This EC_50_ is ∼10,000-fold lower than that of D-glucose (Fu et al., 2014; Urano et al., 2012). The maximum steady-state internalization levels (48% ± 8% and 54% ± 8%) were reached within 30 mins at saturating UDPG concentrations of 2 mM and 100mM, respectively (Figure 2C).

**Figure 2.**
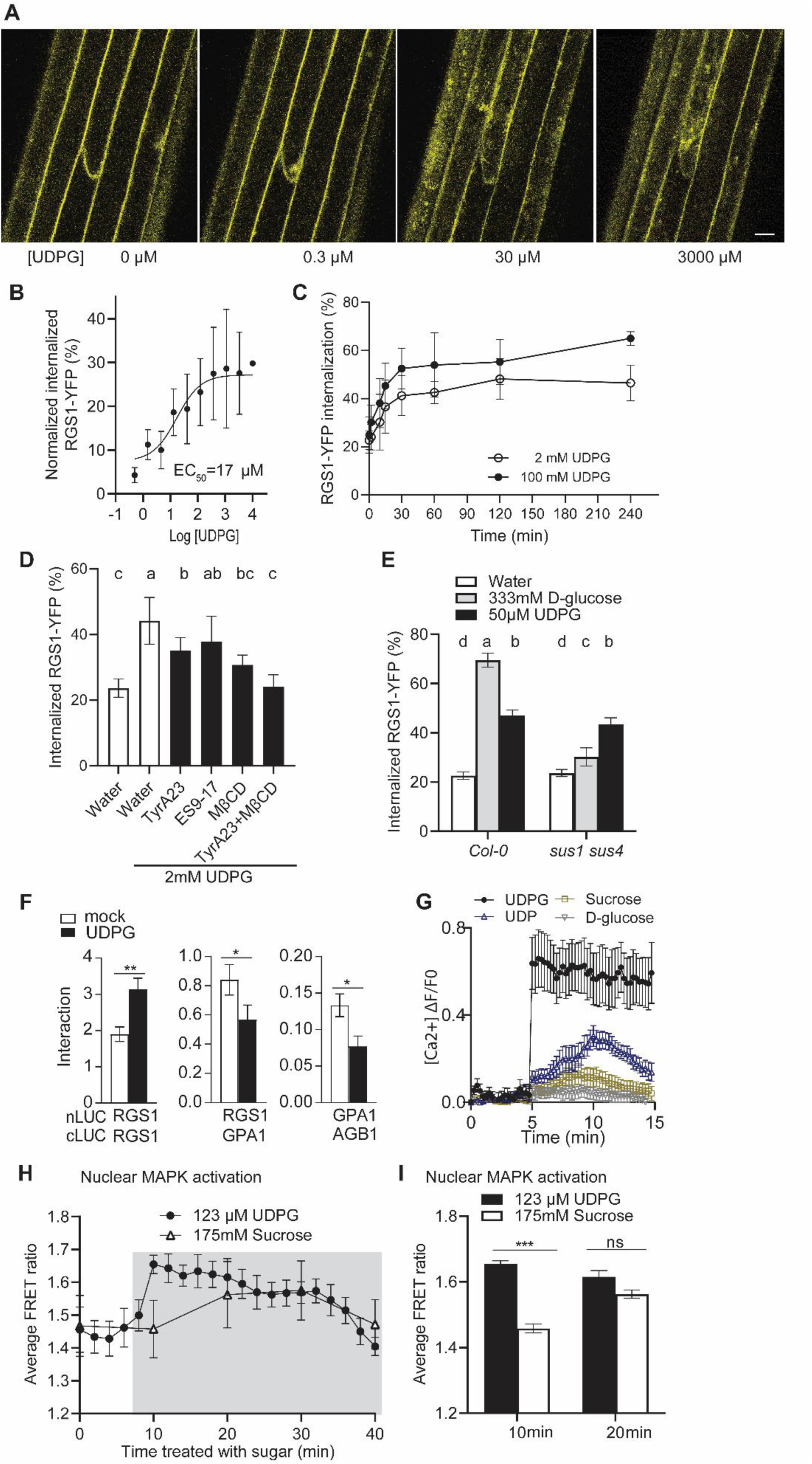
UDPG is a candidate ligand to the RGS1 receptor complex to activate RGS1/G signaling. **(A)** UDPG induces RGS1-YFP internalization. Five-d-old, etiolated seedlings expressing RGS1-YFP were treated with UDPG at 0, 0.3, 30, and 3000 µM and the subcellular localization of RGS1 was imaged at 20 min by confocal microscopy as detailed in methods. Scale bar = 10 µm. (**B**) The dose response of UDPG-induced RGS1-YFP internalization. Five-d-old, etiolated seedlings expressing RGS1-YFP were treated with UDPG at the indicated doses (Log [UDPG]), and the subcellular localization of RGS1-YFP was imaged at 20 mins. Application of UDPG induced RGS1 internalization with an EC_50_ of ∼17µM. Note that this EC_50_ is ∼10,000 times lower than for D-glucose (Fu et al., 2014). Error bars represent standard deviation. (**C**) Time course quantification of RGS1-YFP internalization. Five-d-old, etiolated seedlings expressing RGS1-YFP were treated with 2 mM and 100 mM UDPG, and the subcellular localization of RGS1-YFP was imaged over a time course of 0, 2, 10, 15, 30, 60, 120, and 240 mins using confocal microscopy as detailed in methods. A concentration of UDPG of ∼123 µM is predicted to be just below saturation of the RGS1 response while 2 mM is above saturation. Each data point represents the mean of at least 3 replicates. Error bars represent standard deviation. **(D)** UDPG-induced RGS1-YFP internalization was partially inhibited by MβCD, TyrA23, and ES9-17. Etiolated seedlings expressing RGS1-YFP were treated with water or endocytosis inhibitors (50 µM TyrA23, 100 µM ES9-17, 5 mM MβCD, or 50 µM TyrA23 + 5 mM MβCD) for 60 min, followed by incubation in the same solution supplemented with 2 mM UDPG for 30 min prior to imaging epidermal cells. Internalized RGS1-YFP was quantified to determine total endocytosis of RGS1-YFP. Average values are calculated from n = 10–23 replicates. Error bars = 95% confidence interval. Different letters a, b, c, and d indicate statistically different classes using one-way ANOVA, Tukey’s multiple comparisons test (*P* < 0.05). **(E)** D-glucose-induced RGS1-YFP internalization is SUS1/4 dependent while UDPG is not. Percent internalized RGS1-YFP internalization in response to 30-minute treatment with water, 333 mM D-glucose, or 50 µM UDPG in Col-0 or *sus1/sus4* double mutant. Average values are calculated from n = 38–71 replicates. Error bars represent 95% confidence interval. Different letters a, b, c, and d indicate statistically different classes between treatments using two-way ANOVA, Tukey’s multiple comparisons test (*P* < 0.05). **(F)** UDPG modulates interactions among G protein signaling components RGS1, GPA1, and AGB1(AGG1). Firefly luciferase complementation assay (FLCA) was conducted to assess the impact of 500 µM UDPG on the association of RGS1 dimer and RGS1 with G protein partners. Proteins were transiently expressed in *N. benthamiana* leaves as described in methods. Asterisks denote statistically different classes between treatments calculated by Student’s *t*-test (two tailed): \**P* < 0.05, \*\**P* < 0.01. **(G)** UDPG activates calcium signaling while D-glucose and sucrose do not. Calcium increase was measured *in vivo* in 5-day-old etiolated hypocotyl cells of R-GECO1-expressing Col-0 lines treated with 10 mM UDPG, UDP, D-glucose, or sucrose. Fluorescence intensity changes were quantified in 12-18 regions of interest (ROIs) using ImageJ. Fractional fluorescence changes (ΔF/F) were calculated as (F − F0)/F0, where F0 is the baseline average fluorescence intensity. N = 3 biological replicates per group. Error bars represent SEM. **(H)** UDPG activates nuclear MAPK6. FRET ratios representing nuclear MAPK6 activation (SOMA-NLS) in Col-0 cotyledons treated with 175 mM sucrose and 123 µM UDPG. The graph is a representative of at least 9 cells for each. Sucrose increased FRET ratios in 5 out of 8 cotyledons thus the reason for the higher standard error after sucrose treatment. The difference in kinetics between the two treatments was consistently observed. **(I)** FRET ratios (at indicated 10 and 20 mins) representing nuclear MAPK6 activation (SOMA-NLS) in cotyledons treated with sucrose and UDPG as shown in **(H)**. Asterisks denote statistically different classes between treatments calculated by Student’s *t*-test (two tailed): \*\*\**P* < 0.001; ns = no statistically significant difference.

Previous studies distinguished D-glucose-biased from flg22-biased RGS1-dependent signaling (Watkins *et al*., 2021). The mechanism for this signaling bias occurs through differential compartmentalization of RGS1 with a unique set of interacting partners at the plasma membrane. The MAMP flg22-activated RGS1 pool resides within a membrane domain that is entirely inhibited by tyrphostin A23 (TyrA23) and ES9 (Dejonghe et al., 2019), inhibitors of clathrin-mediated endocytosis. In contrast, the D-glucose-activated RGS1 pool resides equally in two membrane domains; one is inhibited by TyrA23 and the other inhibited by methyl-β-cyclodextrin (MβCD), an inhibitor of sterol-dependent endocytosis, indicating that this RGS1 pool is within lipid rafts. As shown in Figure 2D, a saturating level of UDPG induced RGS1 internalization that was sensitive to both TyrA23 and MβCD, suggesting that the UDPG pool is similar to the previously shown D-glucose bias (P = 0.04 for water *vs.* TyrA23, and *P* = 0.0003 for water *vs.* MβCD using one-way ANOVA). Application of both TyrA23 and MβCD inhibitors completely blocked UDPG-induced RGS1 internalization (P < 0.0001) as previously shown for D-glucose and sucrose. The similar inhibitor profile for UDPG compared with D-glucose is consistent with the hypothesis that D-glucose, sucrose, and likely other sugars, act via their conversion to UDPG, thus tying UDPG action shown here to the large body of literature describing D-glucose and sucrose action on RGS1 trafficking.

### D-glucose-induced RGS1-YFP internalization is SUS1/4 dependent

RGS1 was identified as a potential interactor with SUS1 and SUS4 by a Y2H screen (Klopffleisch et al., 2011). UDPG is metabolized from other sugars, including D-glucose and sucrose (Figure 1A). To determine whether UDPG- or D-glucose-induced RGS1 internalization requires SUS, we quantified RGS1-YFP internalization following 333 mM D-glucose and 50 µM UDPG-treatment in Col-0 and the *sus1/4* double mutant. We found that D-glucose-induced RGS1-YFP internalization was nearly ablated in the *sus1/4* double mutant, while UDPG induced RGS1-YFP internalization was statistically unchanged (Figure 2E). This demonstrates that D-glucose-induced RGS1-YFP internalization is SUS1/4 dependent.

### UDPG activates heterotrimeric G proteins

To further test whether UDPG modulates the association of G protein signaling components RGS1, GPA1, and AGB1(AGG1), we used firefly luciferase complementation assay (FLCA) to quantify the RGS1-RGS1 and RGS1-G protein interactions (Jia et al., 2025; Zhou et al., 2018). UDPG (500 µM) enhanced RGS1 dimerization (Students *t*-test, *P* < 0.01, mock *vs.* 500 µM UDPG, Figure 2F). The reduction of interaction between RGS1-GPA1 and GPA1-AGB1 in response to UDPG suggests that G protein signaling is activated by UDPG (Students *t*-test, *P* < 0.05, mock *vs.* 500 µM UDPG, Figure 2F) by de-sequestration from the inactive RGS1/G protein heterotrimeric complex.

### UDPG activates calcium and MAPK signaling

Five-day-old, etiolated hypocotyls of R-GECO1-expressing Col-0 plants were treated with 10mM UDPG, UDP, sucrose, or D-glucose, and *in vivo* calcium signals were imaged immediately following sugar application (Figure 2G). Treatment with UDPG elicited a rapid increase in calcium levels, reaching its peak in ∼ 5 minutes, and the elevated signal persisted for at least 10 minutes. The equivalent concentration of UDP induced a lower calcium peak that appeared later around 10 minutes before decaying quickly. UDP may be a partial agonist or, based on its lower affinity for RGS1 and the delay in the calcium response, UDP acts indirectly through conversion to UDPG. No significant upregulation of the calcium signal was observed in response to sucrose or D-glucose treatment, as previously reported (Watkins *et al*., 2021).

Given that UDPG activates G proteins and that G proteins are known to couple to MAPK signaling cascades, we tested whether UDPG induces nuclear MAPK activation in Col-0 cotyledon cells *in vivo* with FRET (Zaman et al., 2019). We observed that 123 µM UDPG triggered nuclear MAPK activation, reaching a maximum within 2 minutes, and maintaining this level for approximately 20 min before beginning to decay. Furthermore, we compared MAPK activation induced by a low dose of UDPG and a high dose of sucrose. MAPK activation occurred in Col-0 cotyledons with both UDPG (123 µM) and sucrose (175 mM) but with different kinetics. Sucrose required ∼15 mins to reach the maximum activation vs. 2 mins for UDPG. The delayed MAPK activation observed with a high dose of sucrose (Figure 2H, I) may be due to the time required for prior metabolism to UDPG by SUS. However, further studies are needed to test this possibility.

### UDPG- induced RGS1 internalization is phosphorylation dependent

Urano et al., 2012 (Urano et al., 2012) showed that sugar-induced RGS1 internalization depends on phosphorylation of Ser427, Ser428 and/or Ser435 residues located at the RGS1 C terminal tail, designated the phospho-cluster. Mutation of these serine residues (RGS1^3SA^-YFP) confers loss of internalization response to sugar (Urano et al., 2012). To test if UDPG induces RGS1 phosphorylation, seedlings overexpressing RGS1-TAP were treated with a near-saturating dose of 100 µM UDPG for 0, 3, 15, 30, and 120 min. RGS1-TAP was immunoprecipitated and immunoblotted with antisera against full-length RGS1 and a phospho-specific antibody recognizing Ser427, Ser428, and Ser435 in the C-terminal tail. UDPG (100 µM) induced RGS1 phosphorylation within 3 min (P < 0.05) (Figure 3A) and began decay before 120 min. This result indicates that RGS1 is rapidly phosphorylated in response to UDPG, consistent with the rapid RGS1 internalization observed with a low EC_50_ for UDPG (Figure 2B). Proteasome degradation or other feedback loops that switch off the G protein signaling by dephosphorylation of RGS1 (Watkins et al., 2021) may contribute to the decay of phosphorylated RGS1 pool, which will be addressed in future studies.

**Figure 3.**
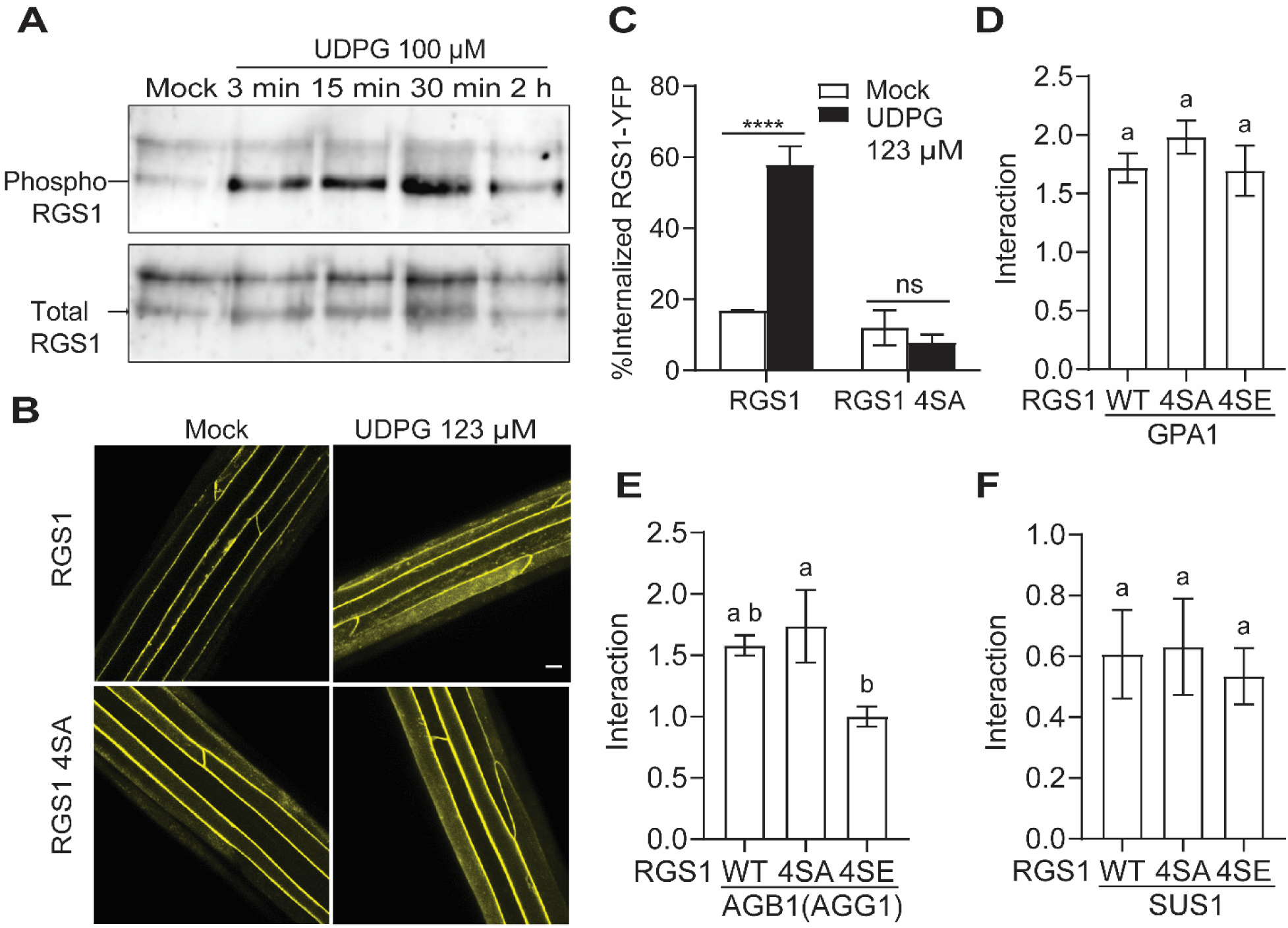
UDPG induced RGS1 internalization is phosphorylation dependent. **(A)** RGS1 is phosphorylated in response to 100 µM UDPG within 3 mins. Seedlings expressing RGS1-TAP were treated with 100 µM UDPG for 3, 15, 30 and 120 minutes, respectively. Phosphorylated RGS1 was detected with an anti-phospho-RGS1 antibody recognizing a cluster of phosphorylated serines in the C-terminal tail. (Urano et al., 2012). Total RGS1 was detected using antisera raised against full-length RGS1. (Urano et al., 2012). **(B)** Phospho-null mutant RGS1 does not respond to UDPG-induced internalization. RGS1-YFP and phospho-null mutant RGS1 4SA (S428/431/435/436A)-YFP were used to test for UDPG-induced internalization. Etiolated hypocotyls stably expressing RGS1-YFP were exposed to 123 µM UDPG for 15 minutes prior imaging. Scale bar = 10 µm. **(C)** Quantification of internalized RGS1-YFP levels shown in panel B. Error bars = SEM, n = 5. Asterisks denote statistically different classes between treatments calculated by Student’s *t*-test (two tailed): \*\*\*\**P* < 0.0001; ns = no statistically significant difference. **(D-F)** RGS1 phosphorylation mutants affect interaction with AGB1 but not GPA1 or SUS1. RGS1-4SA-HiBiT-nLUC (phospho-null) and RGS1-4SE-HiBiT-nLUC (phosphomimic) were transiently co-expressed with cLUC tagged GPA1, AGB1 (AGG1), or SUS1 in *N. benthamiana* leaves. Interactions were measured using FLCA. Relative luminescence unit (RLU) values were normalized to the RGS1-HiBiT signal (Boursier et al., 2020), so the Y-axis has no units. The RGS1-4SE phosphomimic significantly reduces its interaction with cLUC-AGB1, while interactions with cLUC-GPA1 and cLUC-SUS1 remain unaffected.

To test whether UDPG-induced RGS1 internalization is phosphorylation dependent, we used RGS1 phospho-null mutations at Ser427, Ser428, Ser431, and Ser435 (RGS1^4A^-YFP) residues and quantified the percentage of RGS1-YFP and RGS1^4A^-YFP internalization by 123 µM UDPG and mock treated 5-d-old, etiolated seedlings at 20 mins (Figures 3B, C). We observed that 58% RGS1-YFP internalized in response to UDPG *vs.* a baseline of 17% in mock treatment (Student’s *t*-test, two tailed, P < 0.0001), while no internalization difference was observed for the phospho-null mutant RGS1^4A^-YFP in response to UDPG *vs.* mock (8% vs 12%, Student’s *t*-test, two tailed, P = 0.46), suggesting that UDPG-induced RGS1 internalization is dependent on phosphorylation of RGS1 at Ser427, Ser428, Ser431, and Ser435.

To test how RGS1 phosphorylation modulates RGS1-GPA1, RGS1-AGB1(AGG1), and RGS1-SUS1 interaction, we quantified the interaction by FLCA in *N. bethamiana* leaves by transiently expressing the partner proteins (Figure 3D-F). RGS1 4SE phosphomimic mutant reduced RGS1-AGB1(AGG1) interaction (Figure 3E). RGS1 4SE phosphomimic or 4SA phospho-null mutant did not change RGS1-GPA1 and RGS1-SUS1 interaction (Figure 3D and 3F). The results indicate that phosphorylated RGS1 remains associated with GPA1 and SUS1, as observed for the RGS1 WT and the phospho-null RGS1 mutant, while the RGS1 interaction with AGB1 (AGG1) was significantly reduced.

### A Novel SUS/RGS1/G-protein Complex

SUS1 and SUS4 are closely-related members within the SUS family (Supplemental Figure 1), and they are not only present in the cytosol but also associated with the membrane and cell wall (Stein and Granot, 2019). Where and when do RGS1 and SUS1 interact in the cell? Using Bimolecular Fluorescence Complementation (BiFC), RGS1 and SUS1 interacted in punctae along the cell membrane (Figure 4A, red arrows). The RGS1-GPA1 and RGS1-AGB1(AGG1) control interactions showed an even distribution along the plasma membrane (Figure 4A; Supplemental Figure 6, RGS1 and EV ctrl). Because FLCA is reversible, quantitation of the interactions between RGS1 and SUS proteins was performed (Figure 4B). RGS1 interacted with SUS1, SUS4, and SUS3, but not SUS6. The positive controls, RGS1-GPA1, GPA1-AGB1(AGG1), and RGS1-RGS1 showed strong interactions (Figure 4B). RGS1 interacted directly with the AGB1(AGG1) dimer *in vivo* (Figure 4B) suggesting protein scaffolding occurs within the complex.

**Figure 4.**
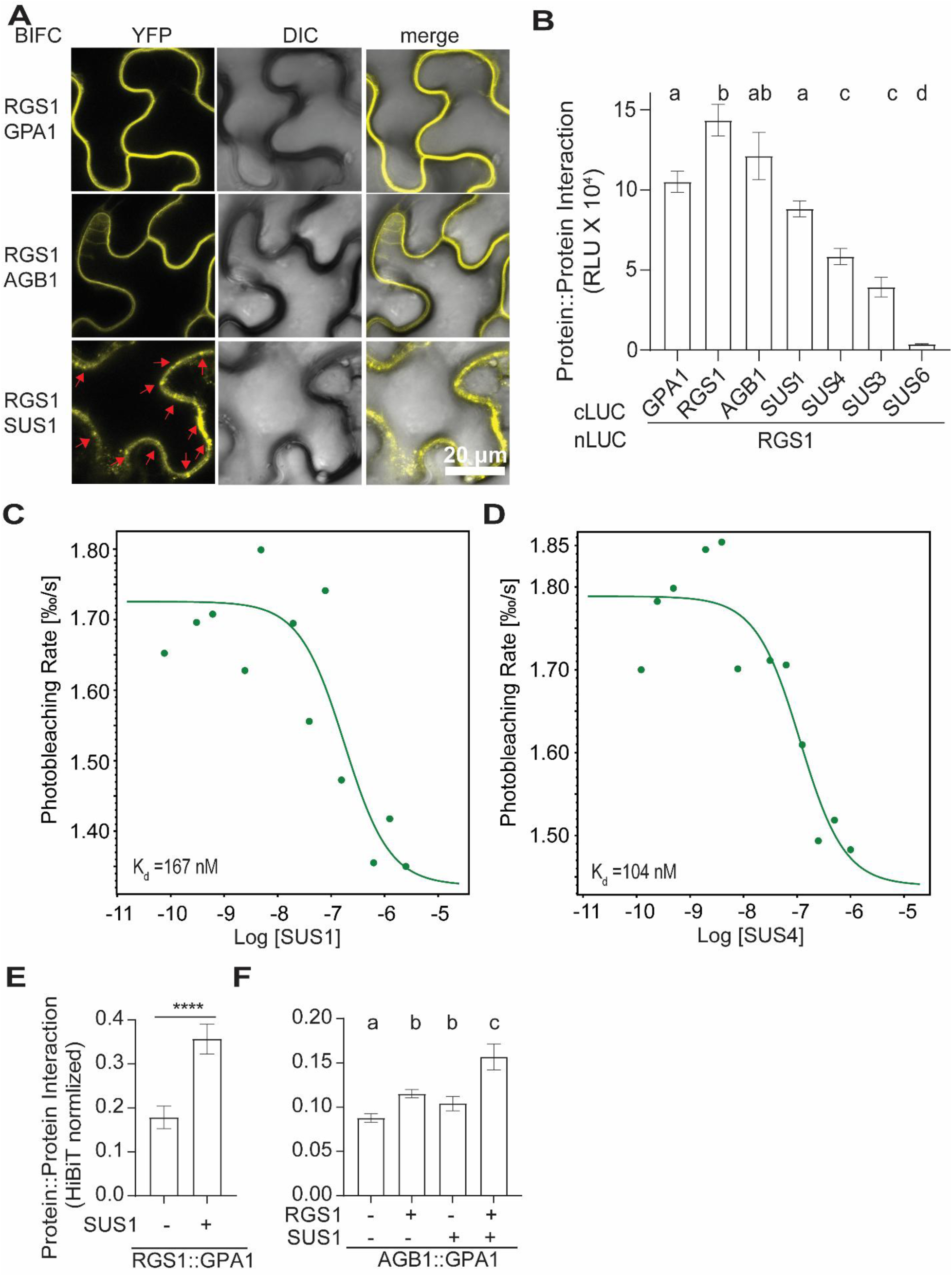
A novel SUS/RGS1/G-protein complex. **(A)** BiFC assays show that RGS1 interacts with SUS1 at the cell periphery. RGS1-nYFP and cYFP-HA-tagged GPA1, AGB1, and SUS1 were transiently expressed in *N. benthamiana* leaves for 48 hours, and YFP signal complementation was imaged to confirm protein interactions and subcellular localization. BiFC between RGS1 and SUS1 shows punctate signals along the cell membrane (red arrows). **(B)** RGS1 interacts with SUS1, SUS3, and SUS4 *in vivo*. FLCA assays were performed by transiently expressing RGS1-nLUC and cLUC-fused GPA1, RGS1, AGB1, SUS1, SUS3, SUS4, and SUS6 in *N. benthamiana* leaves. Luminescence signal was measured to quantify protein interaction strength. **(C, D)** Microscale thermophoresis (MST) assays confirm direct, high-affinity binding of RGS1 to SUS1 **(C)** and SUS4 **(D)**. sfGFP–RGS1 (ΔC-tail) expressed in insect cells was used as the target, and His–SUS1 or His–SUS4 expressed in *E. coli* were used as ligands in binding assays. Binding isotherms show data from one representative replicate. Comparable results were obtained in two independent assays (Supplemental Figures 7 and 8). **(E, F)** SUS1 enhances RGS1-GPA1 and GPA1-AGB1 (AGG1) interactions in *N. benthamiana* leaves. FLCA assays were performed by transiently expressing RGS1-nLUC, AGB1-nLUC, and cLUC-GPA1 in presence and in absence of SUS1 in *N. benthamiana* leaves. Luminescence signal was measured to quantify protein interaction strength.

To confirm direct physical interactions between RGS1 and SUS1 (as well as SUS4), we quantified the *in vitro* binding affinity using MST assays. High-affinity interactions were detected between RGS1 and SUS1 (K_d_ = 161 ± 9 nM; Figure 4C and supplemental Figure 7) and between RGS1 and SUS4 (K_d_ = 106 ± 3 nM; Figure 4D and supplemental Figure 8), consistent with *in vivo* quantitative measurements (Figures 4A, B). The affinities of RGS1 for SUS1 and SUS4 were markedly higher—15-fold and 23-fold, respectively— than that for UDPG (K_d_ = 2.4 ± 0.5 µM; Figure 1B), suggesting that the RGS1–SUS super-complex is constitutively self-stable *in vivo*. This idea is explored next.

SUS1 significantly enhanced the RGS1-GPA1 (Student’s *t*-test, *P* < 0.0001, Figure 4E) and GPA1-AGB1(AGG1) (one-way ANOVA, P < 0.05, Figure 4F) interactions in the FLCA assays suggesting that SUS1 stabilizes the RGS1-GPA1-AGB1 (AGG1) complex. SUS1 and RGS1 acted additively to enhance the GPA1-AGB1(AGG1) interaction (Figure 4F). These results suggests that SUS1 stabilizes the RGS1/G protein complex and acts additively with RGS1 to repress G protein signaling by sequestering the heterotrimeric G protein.

To gain structural insights into the interactions among SUS1, RGS1, and heterotrimeric G proteins, we performed complex structure prediction using AlphaFold3 (https://alphafoldserver.com/) (Supplemental Figures 9A-B, only 1 SUS1 protein shown for clarity). The resulting model predicts that the C-terminal tail of RGS1 (residues S428– G459) forms multiple contact interfaces with SUS1. In addition, the α4 helix of GPA1 appears to contribute to an additional interaction surface with SUS1. These predicted interfaces suggest a coordinated structural arrangement that may stabilize the SUS1– RGS1–G protein complex. Consistent with these modeling results, our previous *in vivo* and *in vitro* assays demonstrated high-affinity interactions between RGS1 and SUS1 in a large oligomeric complex consistent with the punctae formation shown in Figure 4A. Together, these findings provide a testable structural framework for understanding how SUS1 associates with RGS1 and components of the G protein signaling complex. Future work using mutational analyses of the predicted interfaces, targeted crosslinking, and cryo-EM or crystallography will be required to validate or reject this model in solution.

### RGS1 and SUS1 function in the same genetic pathway to repress growth and flg22-induced ROS production

To investigate the genetic interactions between RGS1 and SUS1 in growth and defense we chose three informative phenotypes. These included: rosette leaf size at short day conditions (Figures 5A, B), hypocotyl elongation in 48 h dark-grown seedlings (Figure 5C), and the response to the bacterial elicitor flg22 in reactive oxygen species (ROS) production (Figure 5D). We observed that *rgs1/sus1* mutants qualitatively resemble *rgs1* and *sus1* in these tested growth and defense phenotypes (Figures 5A, 5B, 5C and 5D, relative to Col-0, one-way ANOVA, Tukey’s multiple comparisons test, *P* < 0.05). These findings suggest that RGS1 and SUS1 function in the same genetic pathway. To confirm that the phenotypes observed in the *sus1* mutant is primarily due to de-repression of AGB1, we further generated *sus1 agb1* double mutants. The *sus1 agb1* plants exhibited smaller rosette leaves and reduced ROS, similar to the *agb1* single mutant (Figure 5B and Figure 5D, one-way ANOVA, Tukey’s multiple comparisons test, *P* < 0.05). In sum, these results are consistent with a mechanistic scheme in which the RGS1-SUS1 complex functions upstream to inhibit G protein activation, thereby repressing both growth and defense responses. Binding of UDPG to the RGS1–SUS1 coreceptor may relieve this repression by promoting RGS1 internalization, which in turn releases G proteins from the complex (Figure 2F) to modulate growth and defense programs. Our profile comprises only three phenotypes among many in growth and defense. Therefore, in the next section, we extend validation using transcriptional profiling.

**Figure 5.**
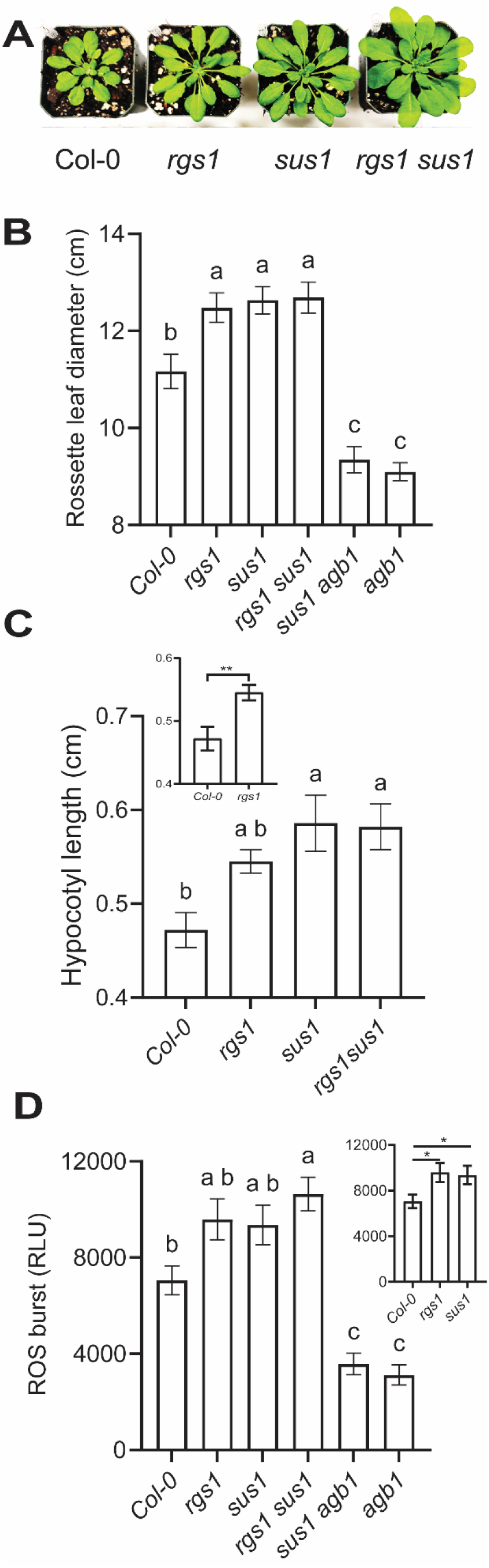
RGS1 and SUS1 function in the same genetic pathway to regulate rosette leaf size, hypocotyl elongation, and flg22-Induced ROS burst. **(A,B)** *rgs1/sus1* mutants resemble single mutants with enlarged rosette Leaves. Plants are 4-week-old grown under short day conditions: 8h light/16h dark, 22 °C. Rosette leaf size of Col-0, *rgs1*, *sus1*, *rgs1 sus1, sus1 agb1,* and *agb1* was measured. n = 12-15 plants are quantified for each genotype. **(C)** *rgs1/sus1* mutants resemble single mutants with longer hypocotyl under dark. Seeds were germinated under dark for 48 hours on ½ MS plates supplemented with no sugar. The hypocotyl length was imaged at 48h and measured by image J. n = 17-28. The graph inset in the upper left shows that the hypocotyl lengths of Col-0 and the *rgs1* mutant differ significantly, consistent with previous findings. Asterisks denote statistically different between Col-0 and the *rgs1* mutant calculated by Student’s *t*-test (two tailed): \*\**P* < 0.01. **(D)** *rgs1/sus1* mutants resemble single mutants with elevated flg22 triggered ROS burst. Plants were 4 weeks old and grown under short-day conditions (8 h light / 16 h dark, 22 °C). For each genotype, 10–15 plants were analyzed. ROS burst was elicited using 100 nM flg22, and luminescence was measured with a plate reader. The inset in the upper right shows that the ROS burst in Col-0, *rgs1*, and *sus1* mutants differs significantly. Asterisks indicate statistical significance between genotypes, calculated using a two-tailed Student’s *t*-test (\**P* < 0.05). Error bars represent SEM. Different letters a, b, and c indicate statistically significant differences between genotypes using one-way ANOVA, Tukey’s multiple comparisons test (*P* < 0.05) in B, C, and D.

### UDPG and RGS1 co-regulated genes enriched with defense markers

Building on our findings that UDPG is a potent ligand capable of activating RGS1/G protein signaling (Figures 1-3) and supported by the observed physical association of the RGS1/SUS1/G-protein super complex (Figure 4) and the genetic interaction among RGS1, SUS1, and AGB1 (Figure 5), we hypothesize that RGS1 and SUS1 act in concert to balance growth and defense. This regulation may occur under basal conditions as well as in response to UDPG perception, functioning through the G protein signaling network and/or additional pathways. This hypothesis was tested using transcriptional profiling. We conducted RNA-seq analysis on 5-day-old, dark-grown Col-0 and *rgs1-2* seedlings, following treatment with either a mock control (NaCl) or a saturating concentration of UDPG for 2.5 hours in the dark (Supplemental Tables 1 and 2). The profile revealed 848 genes that were up-regulated, and 362 genes that were down-regulated in response to UDPG treatment *vs.* mock treatment in Col-0 (Figure 6A; Supplemental Table 1). Gene ontology terms (biological processes) enriched in the set of 848 upregulated-as well as 362 down-regulated genes are shown in Supplemental Figures 10 and 11, respectively. RNA-seq analysis revealed that UDPG induces PTI genes, including receptors (*EFR*, *LYK5*, *RLP6*) and downstream signaling components (*BSK6*, *MAPKKK15/16/18*, *PUB18/23/26*). It also activates the jasmonate/ethylene (JA/ET) signaling pathway, marked by induction of *ERF1/2/15/109/019*, *JAZ1/5/6/7/8*, and *MYC2*. Hormonal crosstalk was evident from upregulation of gibberellin inactivation genes (*GA2OX2/6*), auxin-responsive genes (*IAA5/6/19*), and stress-responsive transcription factors (*WRKY6/22/29/31/38*, *ATAF1/2*). Detoxification enzymes (*APX1*, *PER1*, *GSTU25*), redox regulators (*CYP78A7*, *UGT73B5*), and sugar/cell wall modulators (SWEET15, *EXLA1/2*, *XTH17/19*, *MLO4/6/12*) were also elevated, reflecting enhanced defense readiness. In contrast, UDPG coordinately suppressed JA antagonistic, salicylic acid (SA)-dependent systemic acquired resistance genes: *PAD4*, *EDS5/16*, *FMO1*, *SARD1*, and *WRKY51/54/70*, tryptophan-derived secondary metabolism, and glucosinolate biosynthesis genes. In addition, these growth-related regulator genes were suppressed: *CYCB1/2s*, *CDKB1;2*, *CDKB2;2*, *MYB3R-4*, *TPX2*, *CSLD5*, *MAP65-4*, and *CDC20.2*. This transcriptional shift suggests that elevated extracellular UDPG functions as a signal that activates localized PTI and JA/ET defenses while dampening SA signaling, developmental programs, and systemic immunity.

**Figure 6.**
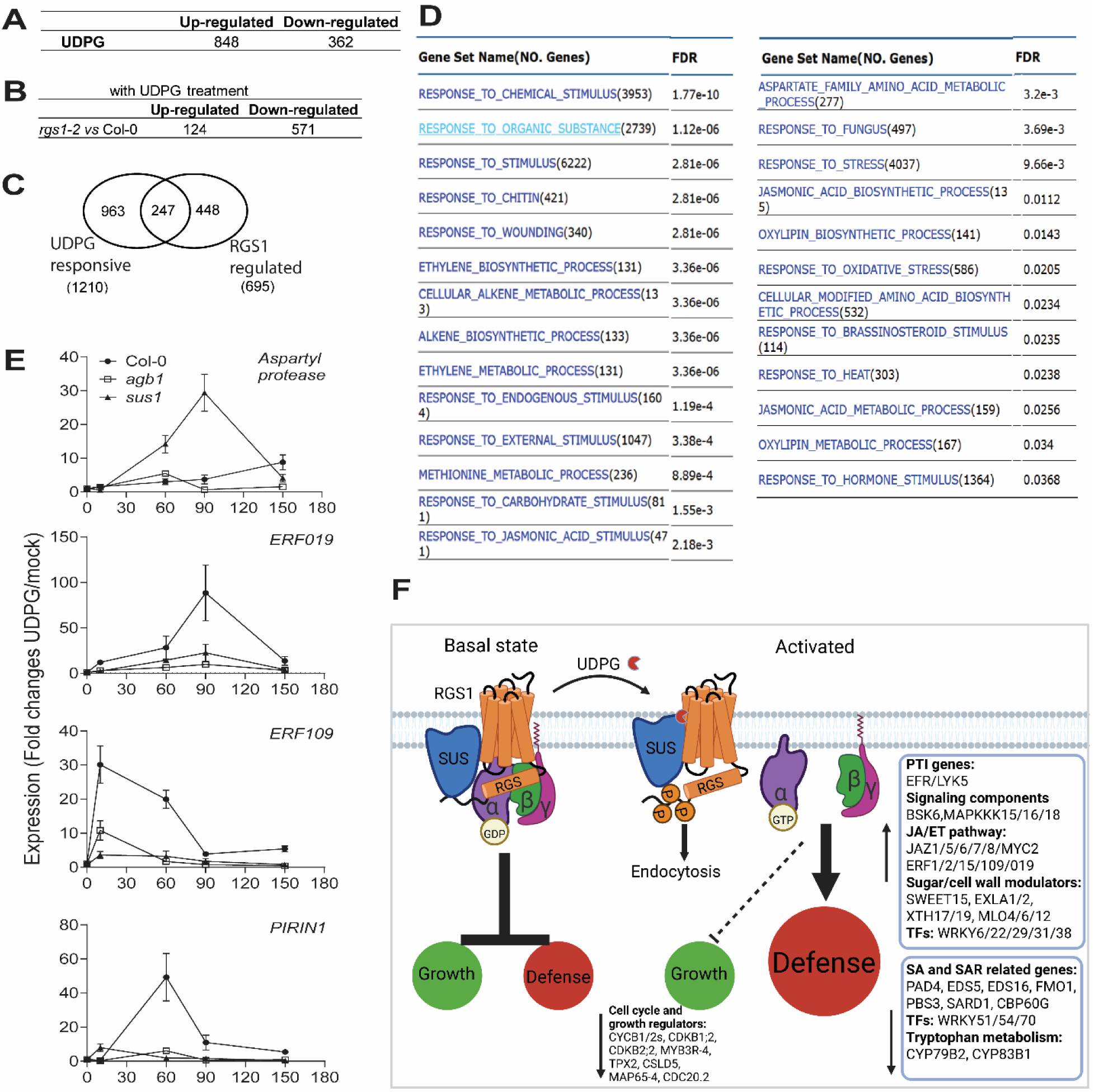
UDPG and RGS1 co-regulated genes enriched with defense markers. **(A)** Number of genes up-and down-regulated by UDPG treatment *vs.* mock in Col-0 were identified by using the DE-seq2 package (P-adj ≤ 0.05; 1.5-fold-change difference). Supplemental Table 1 shows the complete RNA-seq results for all transcriptional response to UDPG in Col-0. RNA seq data is from 4 biological reps of mock treated Col-0 and 3 biological reps of UDPG treated Col-0. **(B)** Transcriptomic analysis of *rgs1-2* mutant response to UDPG treatment. To investigate how RGS1 modulates UDPG induced responses, we performed transcriptomic analysis of *rgs1-2 vs.* Col-0 under the same UDPG treatment conditions (see Supplemental Table 2). **(C)** Identification of UDPG and RGS1 co-regulated genes. By comparing the list of UDPG response genes (Figure 6A, 1210 genes) and RGS1-regulated genes upon UDPG treatment (Figure 6C, 695 genes), we identified 247 co-regulated genes (Supplemental Figure 13 and Supplemental Table 3). **(D)** Functional enrichment of UDPG and RGS1 co-regulated genes. Gene ontology analysis of the 247 UDPG and RGS1 co-regulated genes revealed enrichment in terms related to defense response. **(E)** Transcriptional response of defense marker genes to UDPG and the role of SUS1 and AGB1. Five-day-old Col-0, *sus1*, and *agb1* hypocotyls were treated with mock and 500 µM UDPG for 0, 10, 60, 90, and 150 minutes. Transcript levels of four UDPG- and RGS1-dependent defense marker genes—*At5g19110* (*aspartyl protease*), *ERF019*, *ERF109*, and *PIRIN1*—were quantified. **(F)** Model of UDPG signaling via the RGS1/SUS co-receptor complex and G proteins. For simplicity, the RGS1 dimer and the SUS oligomer are not shown. In the basal state, the RGS1/SUS complex represses heterotrimeric G-protein signaling to maintain growth–defense homeostasis. Upon extracellular UDPG binding, this repression is lifted, activating G protein– and RGS1/SUS–mediated signaling pathways. The resulting transcriptional reprogramming enhances pattern-triggered immunity, jasmonate/ethylene signaling, stress responses, and cell wall remodeling, while concurrently suppressing salicylic acid–mediated systemic immunity, secondary metabolism, and growth. This shift prioritizes local defense over systemic responses and developmental programs.

To investigate how RGS1 modulates UDPG-induced transcriptional responses, we compared *rgs1-2* and Col-0 transcriptomes after UDPG treatment. In *rgs1-2*, 124 genes were upregulated and 571 downregulated relative to Col-0 (Figure 6C; supplemental Table 2), with enriched GO terms related to immunity, metabolism, and hormone signaling (Supplemental Figures 12 A-B). UDPG treatment in the *rgs1-2* mutant leads to upregulation of MAP kinase components (*MAPKKK13*, *MKK7*), ethylene-related transcription factors (*RAP2.6*, *ERS1*), and cytokinin/ABA-responsive genes (*ARR3/4*, *CKX3*, *CYP707A2*), indicating enhanced immune and hormonal signaling in the absence of RGS1. Elevated expression of sugar and amino acid transporters (*STP1/4*, *UMAMIT25/34*), cell wall-growth and remodelling enzymes (*LAC7*, *XTH26*, *BGAL1/4*), and stress-responsive genes (*MIOX4*, *FDH*) suggests that RGS1 restrains metabolic and structural remodeling during defense. In contrast, genes critical for growth, hormone homeostasis (*YUC3*, *IAA19*, *LOG1*, *CKX3*), nutrient transport (*NRT1.5*, *NRT2.1*, *UMAMIT46*, *GDU4*), and cell wall integrity (*XTH20*, *EXLA2*, *LAC17*, *TBL34*) were downregulated. Defense-related genes (*WRKY31*, *PR1*, *CHI*) and the chitin receptor LYK5 were also suppressed, indicating compromised pattern-triggered immunity. These results position RGS1 as a key integrator of UDPG signaling, coordinating immune activation with metabolic balance and growth-defense tradeoffs.

To define genes co-regulated by UDPG and RGS1, we identified 247 overlapping genes between the UDPG-responsive (1,210 genes) and RGS1-regulated (695 genes) transcriptomes (Figure 6C; Supplemental Figure 13; Supplemental Table 3). These genes were significantly enriched in biological processes related to responses to chitin, wounding, fungal pathogens, sugars, and JA (Figure 6D; Supplemental Figure 13; Supplemental Table 3). This co-regulated gene set includes canonical immune markers and hormone-responsive transcription factors (Supplemental Table 3), supporting a role for RGS1 in modulating a core defense program triggered by extracellular UDPG.

We assessed the kinetics and magnitude of Col-0 hypocotyl responses to UDPG at 0, 10, 60, 90, and 150 min by qRT-PCR, measuring transcript levels of four defense marker genes—*At5g19110* (*aspartyl protease*), *ERF019*, *ERF109*, and *PIRIN1*— identified as UDPG-inducible in RNA-seq analysis. We observed rapid transcriptional activation (Figure 6E): *ERF019* peaked as early as 10 minutes, while *ERF109* and *PIRIN1* peaked within 60-90 minutes. At peak levels, *ERF019*, *ERF109*, and *PIRIN1* transcripts increased 30-100-fold from baseline, whereas *At5g19110* (*aspartyl protease*) showed a slower rise, reaching 10-fold by 150 minutes. These results reveal distinct quick and slow response patterns in UDPG-responsive genes.

To elucidate the roles of SUS1 and AGB1 in regulating UDPG-induced defense marker genes (Figure 6E), we assessed the expression of these markers in *sus1* and *agb1* mutants under the same time course and experimental conditions as in Col-0. Among the four UDPG-induced genes analyzed, *At5g19110* (*aspartyl protease*) was suppressed by SUS1 but positively regulated by AGB1, whereas the expression of *ERF019*, *ERF109*, and *PIRIN1* required both SUS1 and AGB1. These results suggest that SUS1 and AGB1 modulate both the kinetics and amplitude of UDPG-induced marker gene expression, highlighting a UDPG signaling pathway perceived by the RGS1-SUS1 complex and transmitted via G protein signaling to downstream targets (Figure 6F).

## CONCLUSIONS

Previous work showed that glucose is a signal that operates through RGS1, however from the onset it was clear that this could not be a direct effect because the concentration of glucose to elicit signaling is too high. These observations and a prediction made by mathematical modeling suggested there is a yet-to-be identified “component X” (Fu et al., 2014) between glucose and RGS1. We show here that “component X” is likely UDPG operating through RGS1 and SUS1 at a physiological level consistent with the measured EC_50_ for induction of RGS1 endocytosis and G protein activation. A new signaling complex of high order is comprised of SUS1, RGS1, GPA1, and AGB1. This complex senses sugar at the plasma membrane through conversion to UDPG which binds within the complex leading to changes in the steady-state levels of mRNA involved in plant defense and growth (Figure 6F).

## METHODS

### Plant materials and growth conditions

For RGS1 internalization assays, wild-type (Col-0) *Arabidopsis thaliana* seeds expressing 35S:RGS1–YFP were surface-sterilized, and 10–20 seeds were sown in 1 mL of liquid ¼-strength Murashige and Skoog (MS) medium without sucrose in 12-well plates. Seeds were stratified at 5 °C for 2 days, exposed to light for 2 h, and then grown in darkness at RT for 3–5 days. For rosette leaf measurements, plants are 4-week-old grown under short day conditions: 8h light/16h dark, 22 °C. For hypocotyl elongation experiments, seeds were surface-sterilized and germinated on ½ MS medium supplemented with no sugar. The plates were incubated in the dark at room temperature (RT) for 48 hours, covered with aluminum foil to prevent light exposure. Hypocotyl length was imaged at 48 hours and measured using ImageJ.

Seedling Growth and Treatment Conditions for RNA-Seq Analysis. *Arabidopsis thaliana* (Col-0 and *rgs1-2*) seedlings were grown in ¼ MS liquid medium without sugar under dark conditions for 5 days, with plates covered with aluminum foil to prevent light exposure. Seedlings were then treated with either mock solution (1mM NaCl) or 500 µM UDPG for 2.5 hours in the dark, with continued light exclusion using aluminum foil. Plant samples were collected immediately after treatment for RNA-seq analysis.

*N. benthamiana* plants were grown in a controlled growth chamber under long-day conditions (16 h light/8 h dark) at 25°C.

### Protein expression and purification

Twinstrep-RGS1-Flag or GFP-twinstrep-RGS1-Flag (hereafter GFP-RGS1) were cloned into pEU-E01-MCS expression vector (Cell Free Science). After plasmid miniprep, plasmid DNA was cleaned and concentrated by DNA clean & concentratorTM-25 (Cat. No. D4334, ZYMO RESEARCH) to ∼ 1µg/µl. In vitro transcription and cell free expression of RGS1 using WEPRO® 7240 Expression Kits (CFS) following the manufacturer’s instruction in the presence of detergents. Translation mixture was spun down 14,000 rpm for 5 min at 4 C, and then the soluble fraction was mixed with 5 volume of binding buffer (1XPBS pH 7.4, 1 mM DTT, 1 X protease inhibitor cocktail, 0.025 % C12E8) and purified by affinity column strep-tactin sepharose (50% suspension, cat no. 2-1201-010, IBA). RGS1 was finally eluted in elution buffer (1XPBS pH 7.4, 1 mM DTT, 1 X protease inhibitor cocktail, 0.025% C12E8, 2.5 mM d-Desthiobiotin).

We prepared microgram amounts of RGS1 in a Cell Free Synthesis (CFS) System charged with wheat germ extract (Li et al., 2016). The yield of soluble RGS1 expressed in CFS was optimized by exploring various types of detergents (Supplemental Figure 2A) for both the lack of inhibition of the translational machinery and for solubilization of RGS1. The nonionic detergents C12E8, MNG-3, Brij35, and Brij58 solubilize better than other tested detergents for expression and purification of RGS1 with minimal inhibition of the CFS reaction.

The construct HA–Flag–10xHis–sfGFP–TEV–Optimized–RGS1(ΔC-tail) was synthesized and cloned into pFASTBAC (GenScript). sfGFP–RGS1(ΔC-tail) was expressed in insect cells and purified following the established protocol for the *Saccharomyces cerevisiae* class D GPCR Ste2 (Velazhahan et al., 2021).

Recombinant expression and purification of SUS1 and SUS4 proteins. Full-length *Arabidopsis SUS1* (At5g20830) and *SUS4* (At3g43190) coding sequences were cloned into the pENTR/D-TOPO vector (Invitrogen) and transferred into the pDEST17 expression vector (Invitrogen) via LR Clonase II Gateway recombination. This vector allows expression of recombinant proteins with an N-terminal 6×His tag under the control of the T7 promoter. Recombinant proteins were expressed in *E. coli* ArcticExpress RP cells (Agilent Technologies). For expression, single colonies were inoculated into 5 mL LB with antibiotics and grown overnight at 37 °C with shaking. This starter culture was used to inoculate 500 mL LB medium, and cells were cultured at 37 °C until OD₆₀₀ reached ∼0.6. Cultures were then cooled on ice for 30 min and shifted to 12 °C. Protein expression was induced with 0.5 mM isopropyl β-D-1-thiogalactopyranoside (IPTG), and cells were incubated for an additional 24 h at 12 °C with shaking (200 rpm). For protein purification, cells were resuspended in lysis buffer [50 mM Tris-HCl (pH 8.0), 300 mM NaCl, 20 mM imidazole, 2 mM MgCl₂, 5 mM 2-mercaptoethanol, 1× protease inhibitor cocktail, 0.1% Thesit (Sigma, 88315), 0.25 mg/mL lysozyme, and 5% glycerol]. Lysates were clarified by centrifugation at 16,000 rpm for 40 min at 4 °C, and the supernatant was incubated with Ni-NTA resin for 30 min at 4 °C with gentle rotation. The resin was washed with wash buffer (same as lysis buffer but without lysozyme and Thesit) and eluted with elution buffer (wash buffer supplemented with 250 mM imidazole). The eluate containing His-SUS1 and His-SUS4 was further purified by size-exclusion chromatography (Superdex 200 10/300 GL, GE Healthcare) in running buffer [30 mM Tris-HCl (pH 8.0), 200 mM NaCl, 1 mM DTT, 150 mM sucrose, 1 mM EDTA, and 10% glycerol]. Purified proteins were aliquoted, snap-frozen in liquid nitrogen, and stored at –80 °C.

### Microscale-thermophoresis assays

RGS1 and ligand binding was estimated by using microscale thermophoresis (Jerabek-Willemsen *et al*., 2011). A range of the indicated concentrations of the required ligand (Supplemental Figure 4) was incubated with 20 nM of purified GFP-RGS1 5 min in assay buffer (1XPBS pH 7.4, 0.025% C12E8). The sample was loaded into the NanoTemper glass capillaries and micro thermophoresis carried out using blue laser with 40% LED power and medium MST. The K_d_ for UDPG was determined using the mass action equation in MO. Affinity Analysis V2.3, based on data from at least three independent experiments. The dissociation constants (K_d_) of the remaining ligands were determined using MO.Control software by fitting the normalized fluorescence response (ΔFnorm) against ligand concentration to a 1:1 binding model with non-linear regression (Jerabek-Willemsen et al., 2011). sfGFP-RGS1 (ΔC-tail) was used to quantify RGS1-SUS1 and RGS1-SUS4 *in vitro* binding affinities in MST assays (Supplemental Figures 7,8). The instrument used was a NanoTemper monolith NT.115.

### Firefly luciferase complementation assays

RGS1-nLUC, AGB1-nLUC, cLUC-GPA1, cLUC-AGB1, cLUC-SUS1, cLUC-SUS3, cLUC-SUS4, and cLUC-SUS6 plasmids were constructed using pCAMBIA/des/nLuc and pCAMBIA/des/cLuc vectors (Lin et al., 2015). The pART27H-mCherry-AtAGG1 plasmid was provided by Dr. Jose R. Botella (University of Queensland, Australia). All constructs were transformed into *Agrobacterium tumefaciens* GV3101 and co-expressed in *Nicotiana benthamiana* leaves by agroinfiltration (Zhou et al., 2018). Luminescence measurements were performed as described previously (Jia et al., 2025).

### BiFC

For BiFC assays, fusion constructs with nYFP or cYFP were transiently introduced into *N. benthamiana* leaves via *Agrobacterium*-mediated infiltration. YFP fluorescence was visualized using a Zeiss LSM 880 confocal microscope.

### Confocal imaging and RGS1 internalization quantification

Images were acquired using a Zeiss LSM 880 confocal laser scanning microscope (Zeiss Microscopy, Oberkochen, Germany) equipped with AiryScan and GaAsP detectors. YFP was excited at 514 nm, and emission was collected between 525 and 565 nm. Image acquisition, processing, and RGS1 internalization measurements were carried out following previously established protocols (Watkins et al., 2021).

### Pharmacological inhibition of RGS1 internalization

Chemicals were purchased or synthesized as described previously (Watkins *et al*., 2021). RGS1 internalization was inhibited using TyrA23, MβCD, or ES9-17 (Dejonghe et al., 2019). Three-day-old seedlings were pre-incubated with each inhibitor at the indicated concentrations for 60 min, then treated with the inhibitor together with 2 mM UDPG for 30 min prior to image acquisition.

### Live cell imaging of MAPK reporter (SOMA-NLS) lines and live cell Ca^2+^ imaging with R-GECO1

Detached 5-day-old etiolated hypocotyls were prepared for confocal microscopy using the HybriWell™ method, following previously described protocols (Watkins et al., 2021). The digital images for MAPK activity and Ca^2+^ signals were analyzed with Fiji (Schindelin et al., 2012).

### RGS1 phosphorylation

*rgs1-2* seedlings complemented with *35S::RGS1-TAP* (Tandem Affinity Purification) were grown for 7 days under low constant light and treated with mock or 100 µM UDPG for 0, 3, 15, 30, and 120 minutes. Total protein was extracted using transmembrane protein extraction buffer (20 % glycerol, 2 % Triton X-100, 1 mM EDTA, 150 mM NaCl, 50 mM Tris-HCl, pH 7.5, 1 mM PMSF). Purification was performed using IgG-Agarose beads (Sigma) and eluted by TEV protease cleavage of TAP tag.

Phosphorylation levels were assessed via western blot analysis using an anti-phospho-RGS1 antibody (Urano et al., 2012), which specifically recognizes phosphorylated Ser428, Ser435, and Ser436, three of the five residues within the serine cluster. Total RGS1 levels were detected using an anti-RGS1 antibody, which binds to the entire cytoplasmic region of RGS1 (Urano et al., 2012).

### Structural modeling of the RGS1-SUS1-G protein supercomplex

The structural model of the RGS1-SUS1-G protein supercomplex was generated using AlphaFold3 (https://alphafoldserver.com/), an advanced protein structure prediction algorithm developed by Google DeepMind. AlphaFold3 utilizes deep learning-based methods to predict protein-protein interactions and complex assemblies with high accuracy. The input sequences for RGS1, SUS1, and heterotrimeric G protein components were submitted to the AlphaFold3 modeling pipeline, and the resulting structures were analyzed for interface interactions and structural stability.

### ROS burst assay

Four-week-old *Arabidopsis thaliana* plants grown under short-day conditions (8 h light/16 h dark, 22 °C) were used for ROS assays. Leaf discs (6 mm diameter) were collected with a cork borer and incubated overnight in sterile water in 96-well plates (Greiner Bio-One, Cat. No. 655075). On the following day, the water was exchanged for assay solution containing 17 mg/mL luminol (Sigma-Aldrich, Cat. No. A8511), 10 mg/mL horseradish peroxidase (Kementec, Cat. No. 4120), and 100 nM flg22 (GenScript, Cat. No. RP19986). Luminescence was recorded immediately at 470 nm using a SpectraMax L Microplate Reader (Molecular Devices).

### RNA-seq library preparation and sequencing

Total RNA was extracted using Monarch® Total RNA Miniprep Kit (T2010S) following manufacturer recommendations. Extracted RNA was quantified on Qubit (Invitrogen), and RNA Integrity value (RIN) was estimated based on TapeStation (Agilent) result. RNA with a RIN value above seven was used for library preparation. We used the KAPA Stranded mRNAseq system (Roche) for messenger RNAseq library preparation. After completion of library preparation, all samples were pooled based on the qubit and TapeStation results. The prepared pool was sequenced on one lane of the NovaSeq6000 SP PE 2×50 system producing around 30 million clusters per sample. Raw data were uploaded to UNC servers for further analysis.

### Transcriptional profiling

RNA-seq data analysis was performed using the Galaxy platform (https://usegalaxy.org/) for sequencing data generated by the Illumina NovaSeq6000 SP platform. The raw sequencing files in FASTQ format were uploaded to Galaxy, and quality control was conducted using FastQC to assess read quality, GC content, and adapter contamination. Low-quality reads and adapters were removed using Trimmomatic. The cleaned reads were aligned to the reference genome of Arabidopsis thaliana using HISAT2, and transcript quantification was performed with StringTie to obtain gene expression levels. Differential expression analysis was carried out using DESeq2 to identify significantly differentially expressed genes.

Gene ontology and pathway enrichment analyses were performed using PlantGSEA toolkit (Ma et al., 2022; Yi et al., 2013) (https://systemsbiology.cau.edu.cn/PlantGSEAv2), and ShinyGO 0.82 (https://bioinformatics.sdstate.edu/go/).

### Gene expression analysis by quantitative real-time PCR

To assess the transcriptional response of defense marker genes to UDPG, 5-day-old dark-grown Col-0, *sus1*, and *agb1* seedlings were treated with mock or 500 µM UDPG for 0, 10-, 60-, 90-, and 150-min. Total RNA was extracted using the Monarch® Total RNA Miniprep Kit (Cat. #T2010S, NEB) and quantified with a NanoDrop 2000 UV/VIS Spectrophotometer (Thermo Scientific). First-strand cDNA was synthesized with the LunaScript RT SuperMix Kit (Cat. #E3010, NEB). qRT-PCR was performed with the Luna® Universal qPCR Master Mix (Cat. #M3003, NEB) on a QuantStudio 7 Flex Real-Time PCR System (Applied Biosystems). UBQ10 was used as the internal reference gene. Relative expression was calculated by the 2^(-ΔΔCt) method (Schmittgen and Livak, 2008), with results presented as fold change (UDPG/mock). Primer sequences are listed in Supplemental Table 4.

### Statistical analysis

The data in Figures 2-5 were analyzed by One way ANOVA or two-way ANOVA test (*P* < 0.05) in GraphPad prism. The data in Figure 2F, 2I, 3C, 4E, 5C and 5D, were analyzed by two-tailed Student t-test: *, *P* < 0.05; **, *P* < 0.01; ***, *P* < 0.001.

## RESOURCE AVAILABILITY

### Lead contact

Further information and requests for resources and reagents should be directed to and will be fulfilled by the lead contact, Haiyan Jia.

### Materials availability

The materials generated in this study are available from the lead contact possibly with a completed Materials Transfer Agreement if not already deposited to the Arabidopsis Biological Research Stock Center for public distribution.

## FUNDING

This work was supported by grants to A.M.J. from the NIGMS, NSF (MCB-1713880). The Division of Chemical Sciences, Geosciences, and Biosciences, Office of Basic Energy Sciences of the US Department of Energy through the grant DE-FG02-05er15671 to A.M.J. funded work on optimizing GFP-RGS1 expression. C.G.T was supported by core funding from the UK Medical Research Council [MRC U105197215]. For the purpose of open access, the MRC Laboratory of Molecular Biology has applied a CC BY public copyright license to any author accepted manuscript version arising.

## ACKNOWLEDGEMENTS

We thank Jing Yang for technical assistance. We thank Dr. Ken Jacobson for providing P2Y ligands, guidance in testing for functional homology to the P2Y receptors and editing the manuscript.

## AUTHOR CONTRIBUTIONS

H.J., F.L., J.W., and C.C.O. performed experiments and analyzed data. C.G.T. assisted with RGS1 expression in insect cells. H.J. and A.M.J. wrote the manuscript and designed the figures. All authors, together with Dr. Ken Jacobson, contributed to editing. A.M.J. managed the entire project.

## Notes

### Competing Interest Statement

The authors have declared no competing interest.

