## SUPPLEMENTAL INFORMATION for "UDP-glucose Activation of a G-protein/Sucrose Synthase Signaling Supercomplex"

This PDF file includes:

Figures S1 to S13

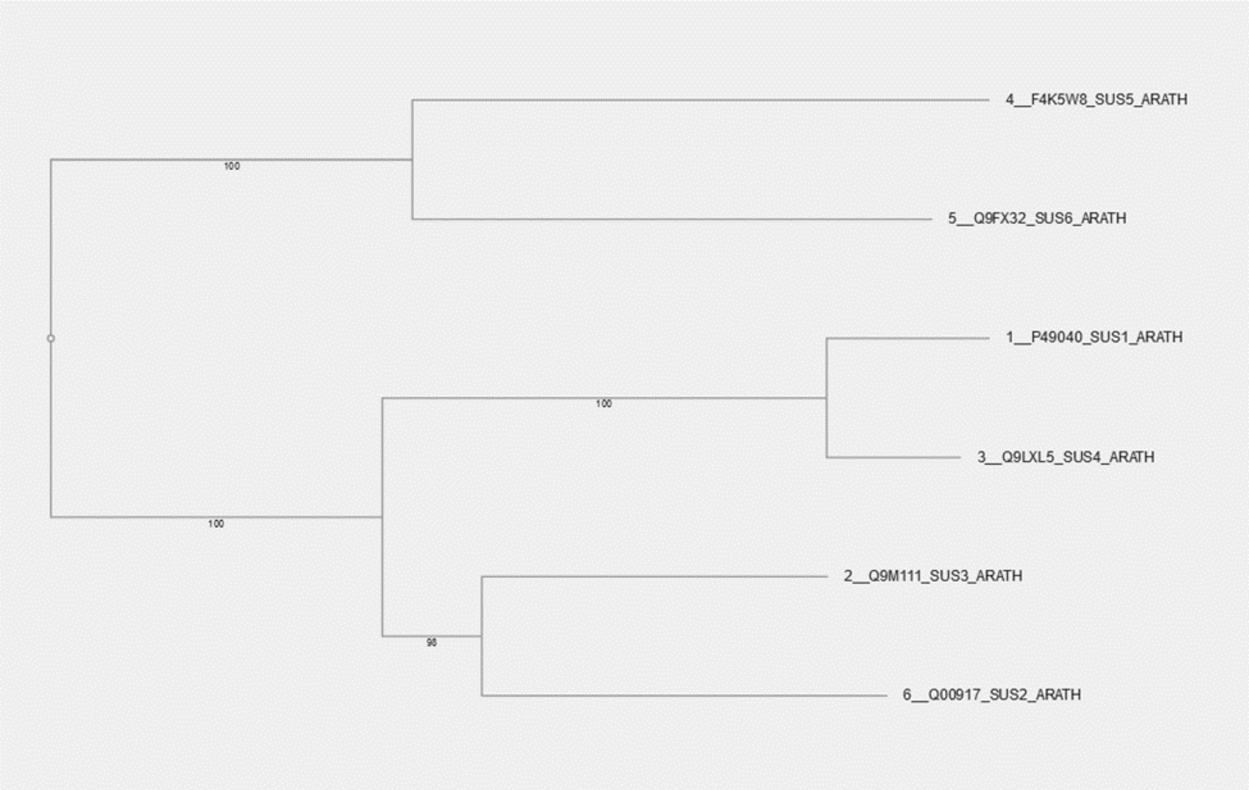


**Figure S1.**

**Phylogenetic relationships and subcellular localization of Arabidopsis sucrose synthase (SUS) isoforms**

A phylogenetic tree was constructed using full-length sequences of six Arabidopsis sucrose synthase (SUS) proteins (SUS1 to SUS6). Sequence alignment and tree generation were performed using the MAFFT online server (https://mafft.cbrc.jp/alignment/server/phylogeny.html). The alignment included 6 sequences across 788 amino acid sites. Phylogeny was inferred using the Neighbor-Joining method with the JTT substitution model, assuming a uniform rate of evolution (Alpha = ∞). Bootstrap analysis was conducted with 1000 replicates to assess branch support. Predicted subcellular localization of sucrose synthase isoforms: SUS1, cytosol and plasmodesmata (Aryal et al., 2014; Fernandez-Calvino et al., 2011); SUS2, cytoplasm, plastid membrane, and peripheral membrane(Nunez et al., 2008); SUS4, vacuole(Szponarski et al., 2004); SUS5 and SUS6, cell wall and secreted(Barratt et al., 2009).

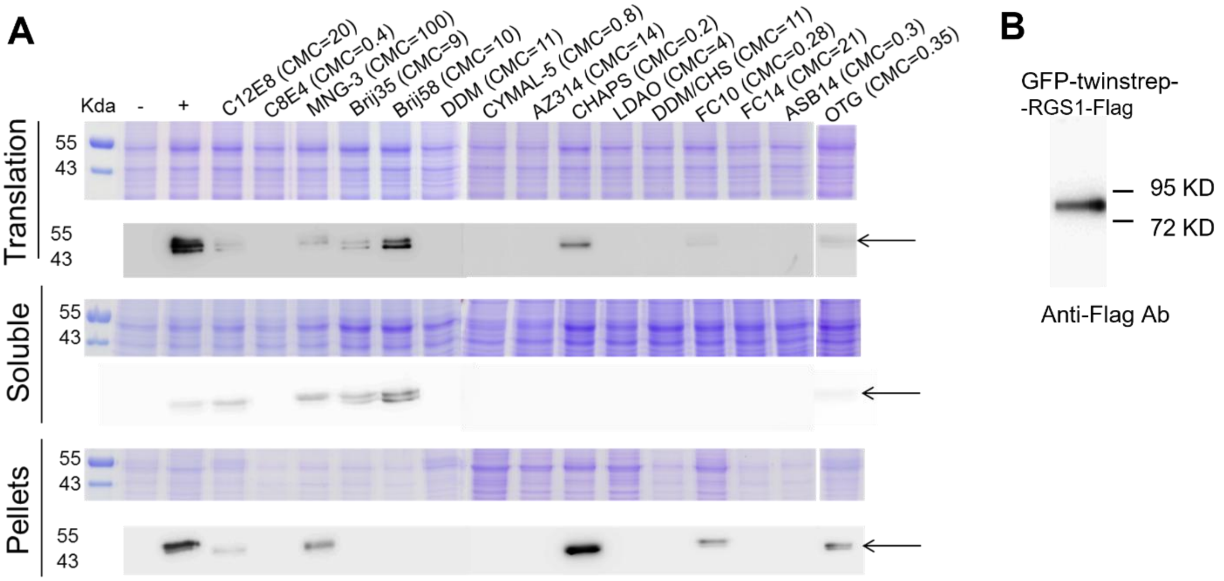


**C**

| | |
|---|---|
| Name: | UDP-glucose Tray |
| Graph Color: | ● |
| Target Name: | R72-GFP-RGS1 |
| Target Concentration: | 20 nM |
| Ligand Name: | UDP-glucose |
| Ligand Concentration: | 156 μM to 0.00954 μM |
| n: | 5 |
| Comments: | |
| Excitation Power: | 40% |
| MST Power: | 40% |
| Temperature: | 22.3°C |
| Kd: | 2.3759E-06 |
| Kd Confidence: | ± 5.3189E-07 |
| Response Amplitude: | 17.881259 |
| TargetConc: | 2E-08[Fixed] |
| Unbound: | 259.08 |
| Bound: | 276.96 |
| Std. Error of Regression: | 1.275515 |
| Reduced $\chi^2$: | 0.10977203 |
| Signal to Noise: | 15.142093 |

**Figure S2.**
**Expression, purification, and MST analysis of GFP–RGS1 produced by wheat germ cell-free system**

**A. Expression of RGS1 using the wheat germ cell-free expression system (CFS) in the presence of 15 different detergents**

Twinstrep-RGS1-Flag was expressed from CFS. Detergents used in wheat germ cell-free expression of RGS1 are: C12E8, dodecyl octaethylene glycol ether; C8E4, tetraethylene Glycol Monooctyl Ether; MNG-3, lauryl maltose neopentyl glycol; Brij35, polyethylene glycol (23) monododecyl ether; Brij58, Polyethylene glycol hexadecyl ether; DDM, n-dodecyl- β - D -maltoside; CYMAL-5, 5-Cyclohexyl-1-Pentyl-β D-Maltoside; AZ314, n-Tetradecyl-N,N-Dimethyl-3-Ammonio-1-Propanesulfonate; CHAPS, 3-((3 cholamidopropyl)dimethylammonio)-1-propansulfonate; LDAO, lauryl dimethyl amide oxide; DDM/CHS, Dodecyl-β-D-Maltoside (100 mg/ml):CHS-TRIS (10 mg/ml) Solution; FC10, Fos-Choline-10; FC14, Fos-Choline-14; ASB14, 3-[N,N-Dimethyl(3-myristoylaminopropyl)ammonio]propanesulfonate; OTG, n-Octyl beta-D-thioglucopyranoside. We universally used 0.1% detergent concentration and the corresponding Critical micelle concentration (CMC) is indicated.
Protein samples were analyzed by SDS-PAGE followed by coomassie blue staining (upper panels) and western blotting by anti-Flag antibody (lower panels). Translation, cell free reaction mixture; Soluble and Pellet obtained after centrifugation of cell free reaction mixture; -, negative control (no RGS1 mRNA); + , positive control (RGS1 expressed in the absence of detergent). The black arrowheads indicate RGS1. Translation-2% of total sample (220 µl reaction mix); Soluble and Pellet-26% of total sample (220 µl reaction mix) loaded on the gel.

**B. Detection of purified CFS-expressed GFP-RGS1 by western blot.**

GFP-Twinstrep-RGS1-Flag was expressed using a wheat germ cell-free system (CFS) in the presence of 0.1% C12E8 detergent. The protein was purified and analyzed by western blotting using an anti-Flag antibody.

**C. Goodness-of-fit analysis of MST binding curve for UDPG and GFP–RGS1 interaction**

Data (n = 5, see Figure S4) were fitted to a one-site binding model using MO. Affinity Analysis software (NanoTemper Technologies). Key fitting statistics are presented.

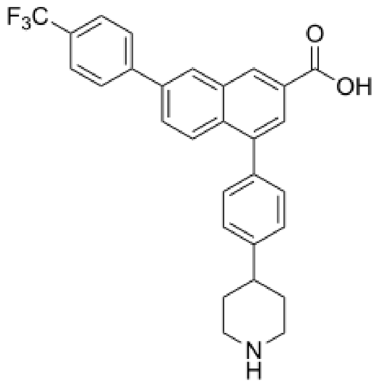

**PPTN**

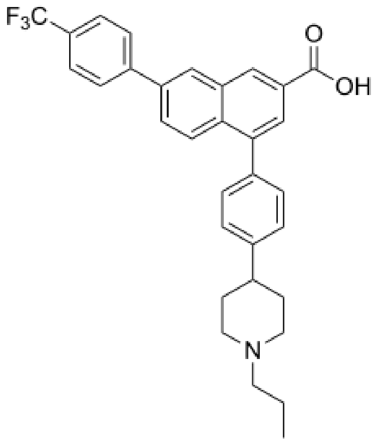

**MRS4574**

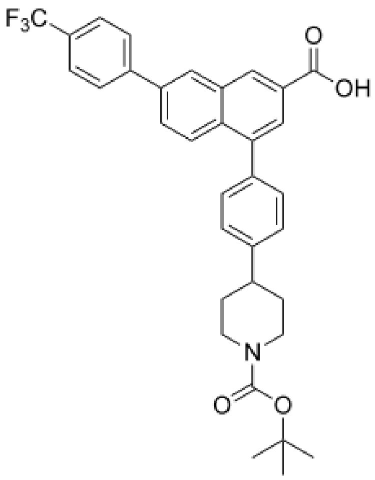

**MRS4575**

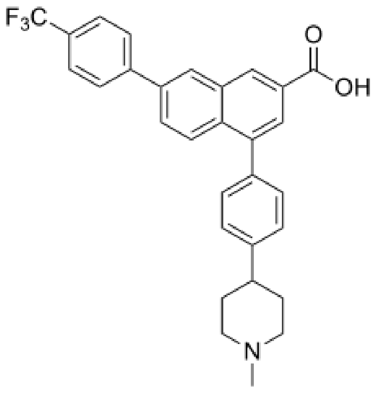

**MRS4576**

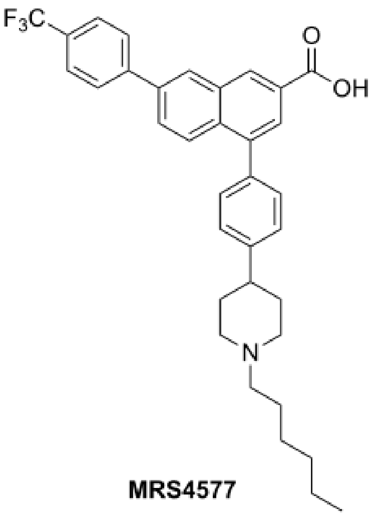

**MRS4577**

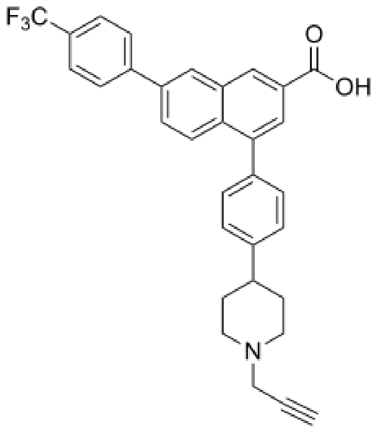

**MRS4578**

**Figure S3.**

**Chemical structures of $P2Y_{14}$ antagonists tested in Figure 1C.**

Structures of the $P2Y_{14}$ receptor antagonists evaluated for activity are shown.

**Negative control**

Target: GFP

Ligand: UDPG

Response evaluation: Initial Fluorescence

Signal to Noise: 0.5 (should be ≥ 5 to conclude target-ligand binding)

Conclusion: No binding

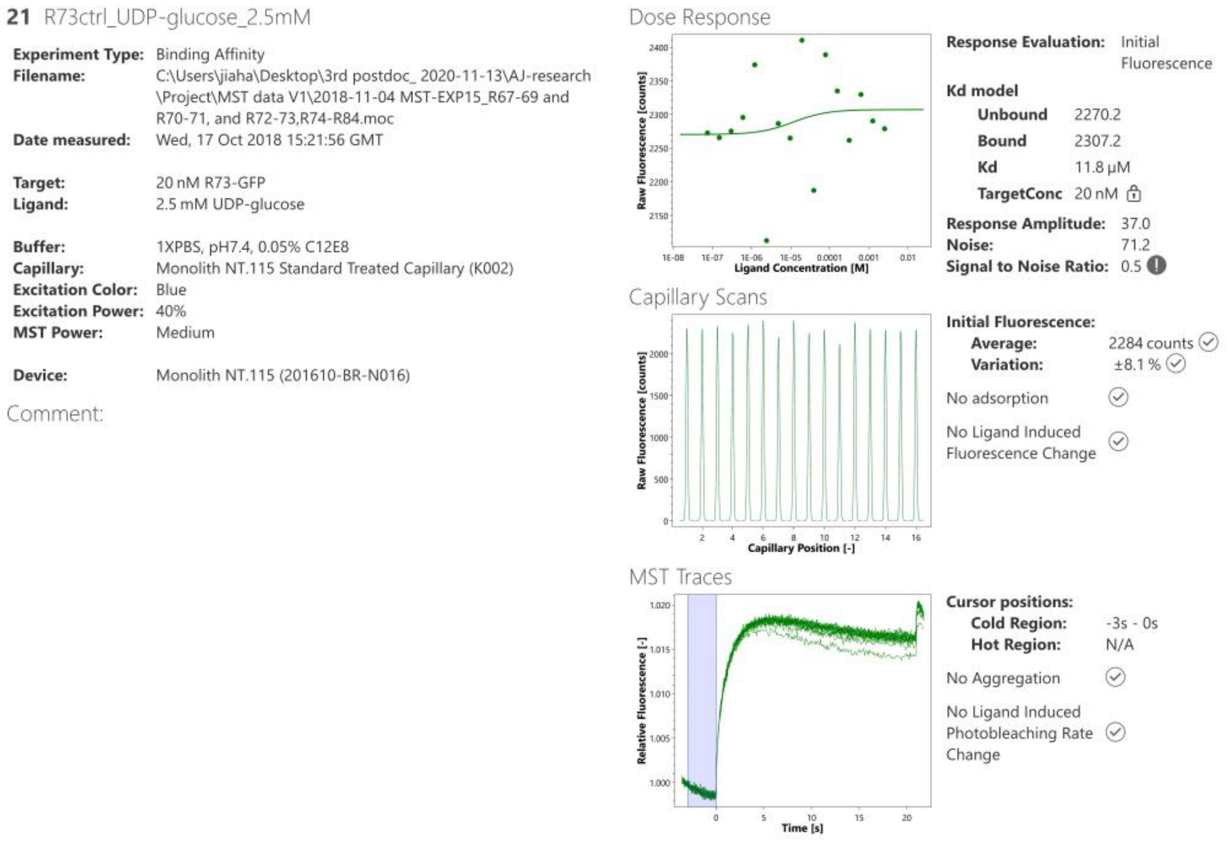


**Figure S4.**

**Binding profiles of tested ligands to GFP–RGS1 determined by microscale thermophoresis (MST).**

Assay conditions are described in Methods.

**Ligand class: Sugar**

Target: GFP-RGS1

Ligand: D-Glucose

Rep1

Response evaluation: Initial Fluorescence

Signal to Noise: N/A (should be ≥ 5 to conclude target-ligand binding)

Conclusion: No binding

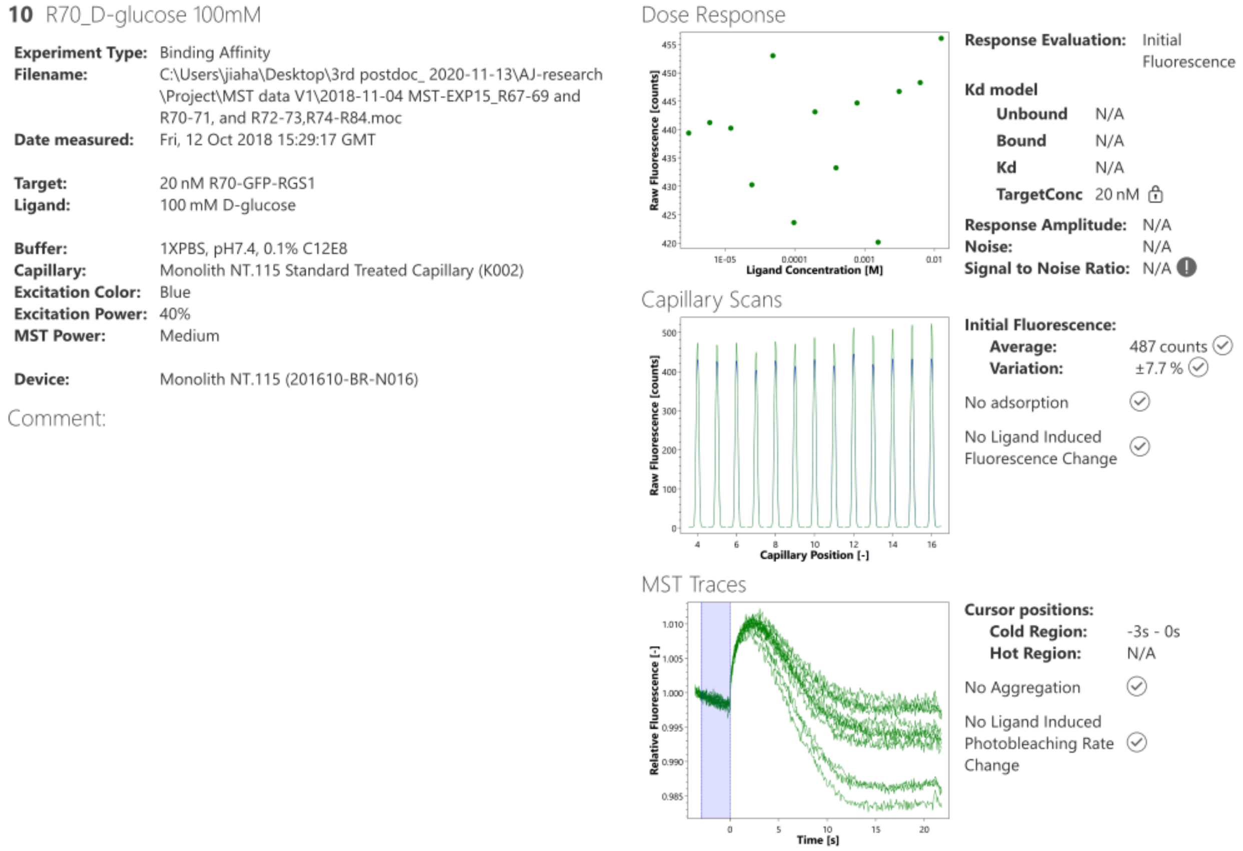


**Figure S4 (continued)**

**Binding profiles of tested ligands to GFP–RGS1 determined by microscale thermophoresis (MST).**

Assay conditions are described in Methods.

**Ligand class: Sugar**

Target: GFP-RGS1

Ligand: D-Glucose

Rep2

Response evaluation: Initial Fluorescence

Signal to Noise: 5.4 (should be ≥ 5 to conclude target-ligand binding)

Conclusion: binding

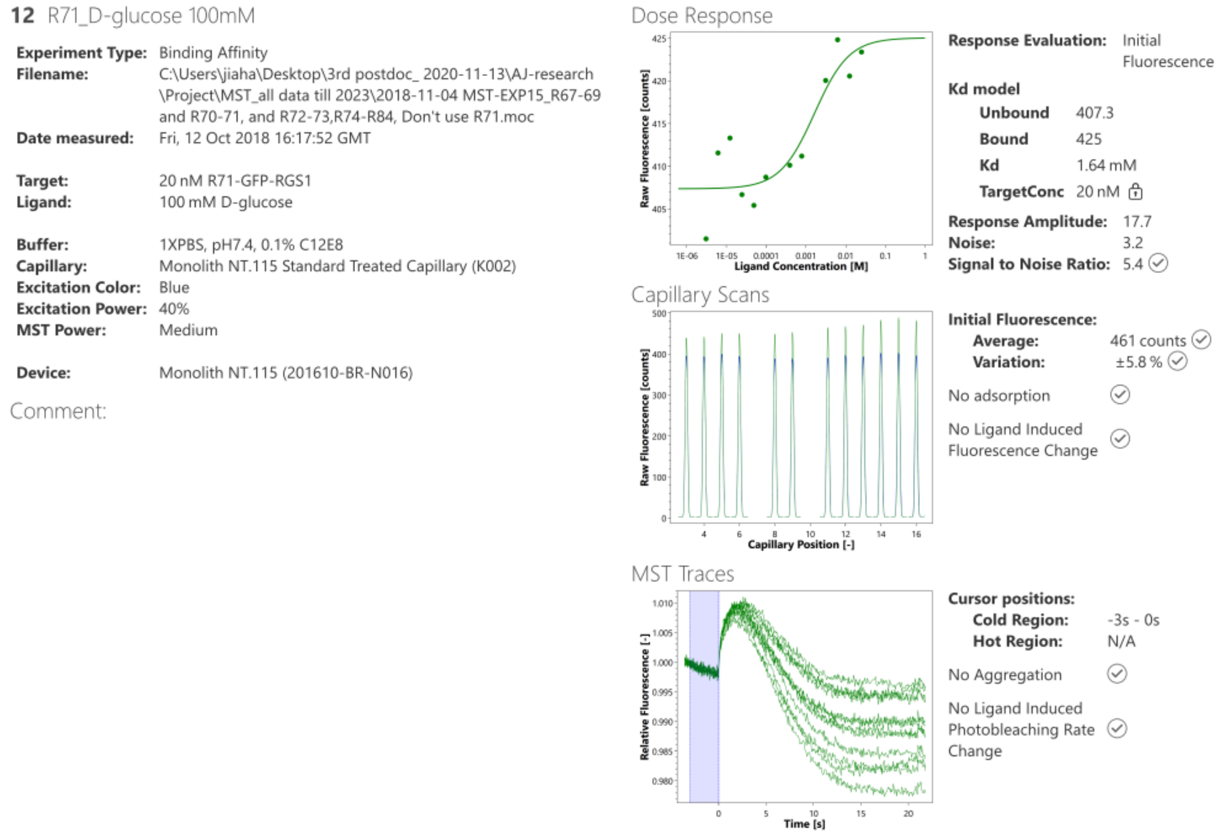


**Figure S4 (continued)**

**Binding profiles of tested ligands to GFP–RGS1 determined by microscale thermophoresis (MST).**

Assay conditions are described in Methods.

**Ligand class: Sugar**

Target: GFP-RGS1

Ligand: L-Glucose

Response evaluation: Initial Fluorescence

Signal to Noise: N/A (should be ≥ 5 to conclude target-ligand binding)

Conclusion: No binding

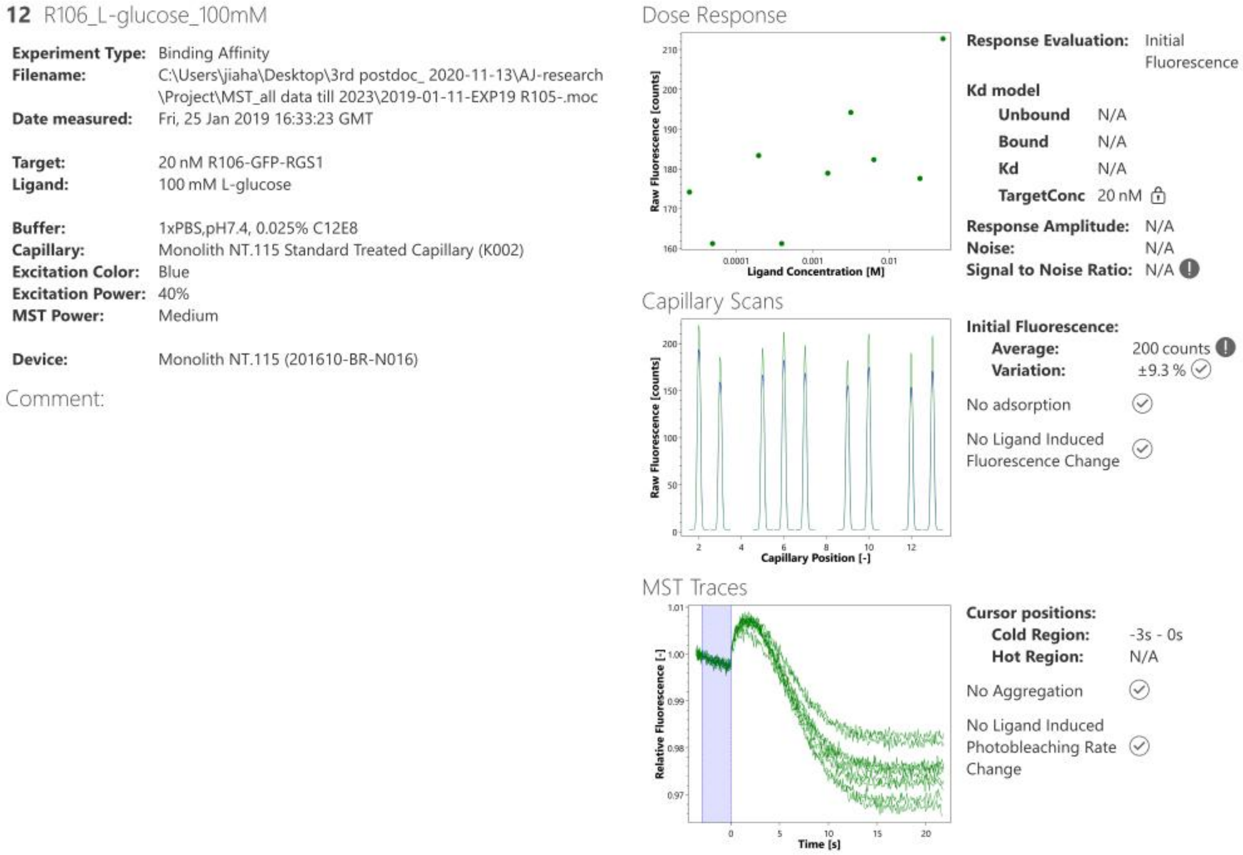


**Figure S4 (continued)**

**Binding profiles of tested ligands to GFP–RGS1 determined by microscale thermophoresis (MST).**

Assay conditions are described in Methods.

**Ligand class: Sugar**

Target: GFP-RGS1

Ligand: Sucrose

Rep1

Response evaluation: Initial Fluorescence

Signal to Noise: 1.5 (should be ≥ 5 to conclude target-ligand binding)

Conclusion: No binding

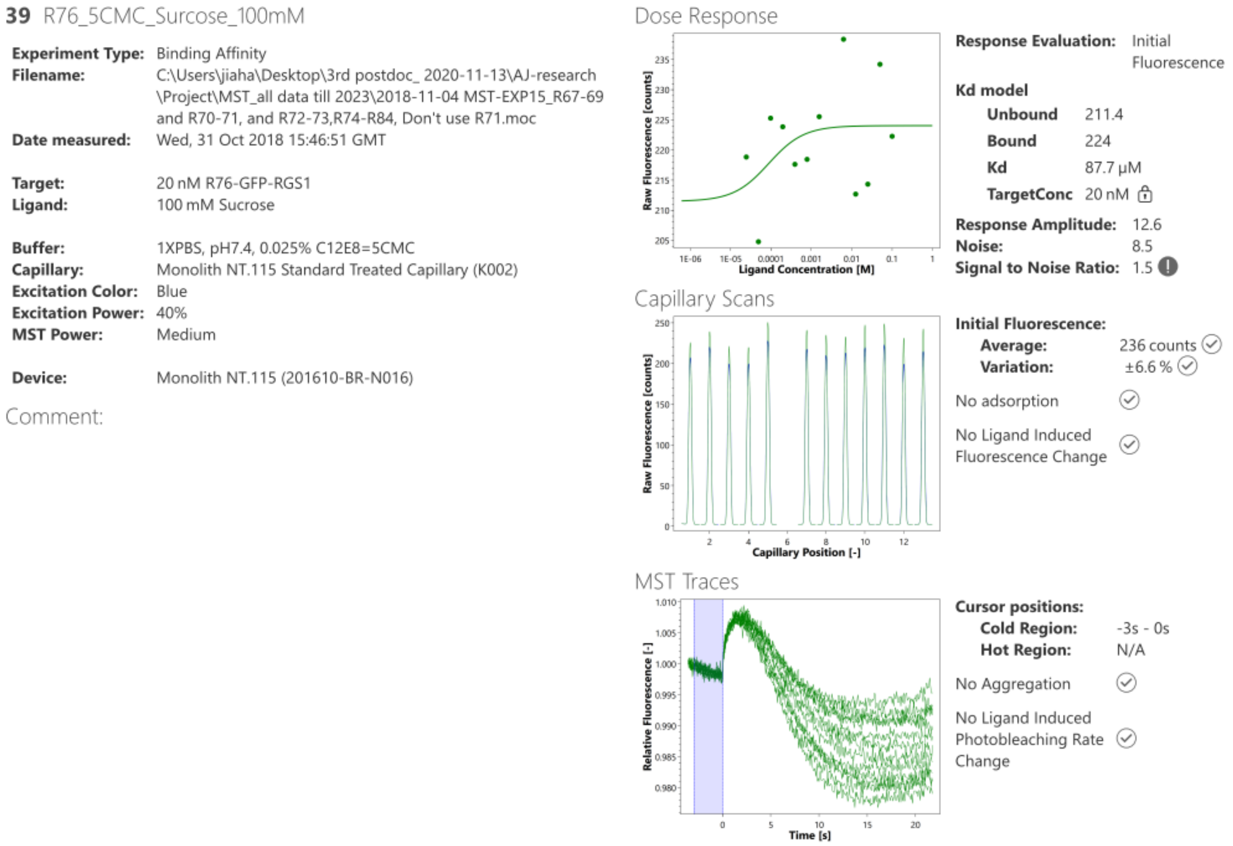


**Figure S4 (continued)**

**Binding profiles of tested ligands to GFP–RGS1 determined by microscale thermophoresis (MST).**

Assay conditions are described in Methods.

**Ligand class: Sugar**

Target: GFP-RGS1

Ligand: Sucrose

Rep2

Response evaluation: Initial Fluorescence

Signal to Noise: 6.8 (should be ≥ 5 to conclude target-ligand binding)

Conclusion: binding

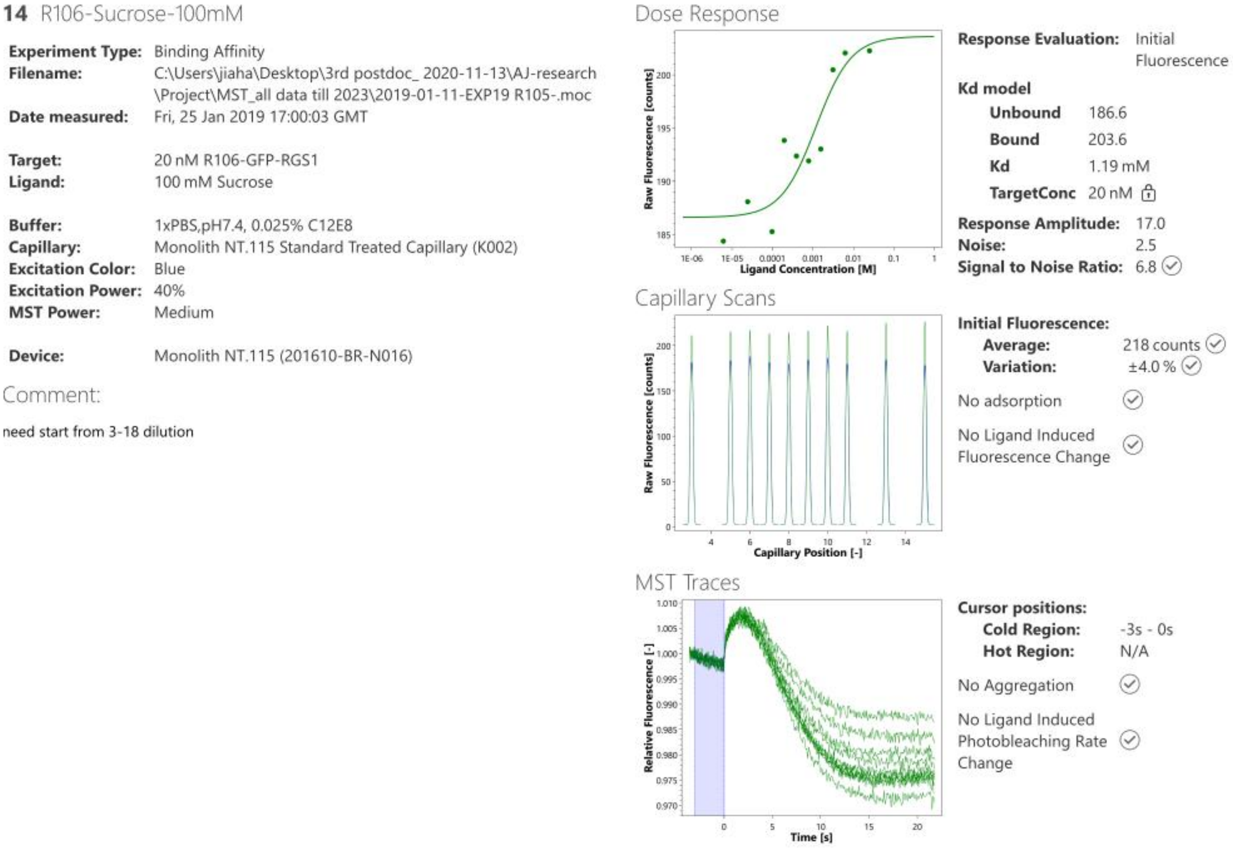


**Figure S4 (continued)**

**Binding profiles of tested ligands to GFP–RGS1 determined by microscale thermophoresis (MST).**

Assay conditions are described in Methods.

**Ligand class: Sugar**

Target: GFP-RGS1

Ligand: D-Fructose

Rep1

Response evaluation: Initial Fluorescence

Signal to Noise: 1.5 (should be ≥ 5 to conclude target-ligand binding)

Conclusion: No binding

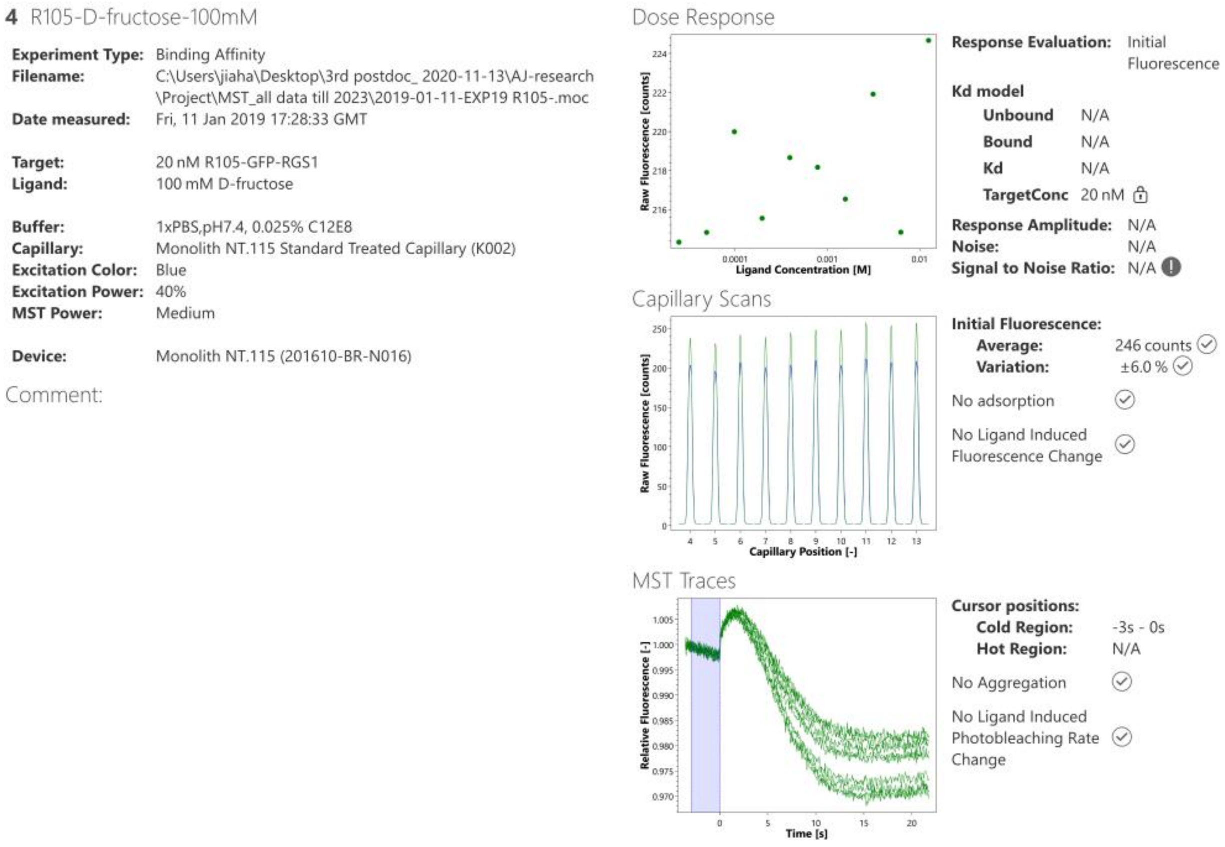


**Figure S4 (continued)**

**Binding profiles of tested ligands to GFP–RGS1 determined by microscale thermophoresis (MST).**

Assay conditions are described in Methods.

**Ligand class: Sugar**

Target: GFP-RGS1

Ligand: D-Fructose

Rep2

Response evaluation: Initial Fluorescence

Signal to Noise: 6.3 (should be ≥ 5 to conclude target-ligand binding)

Conclusion: binding

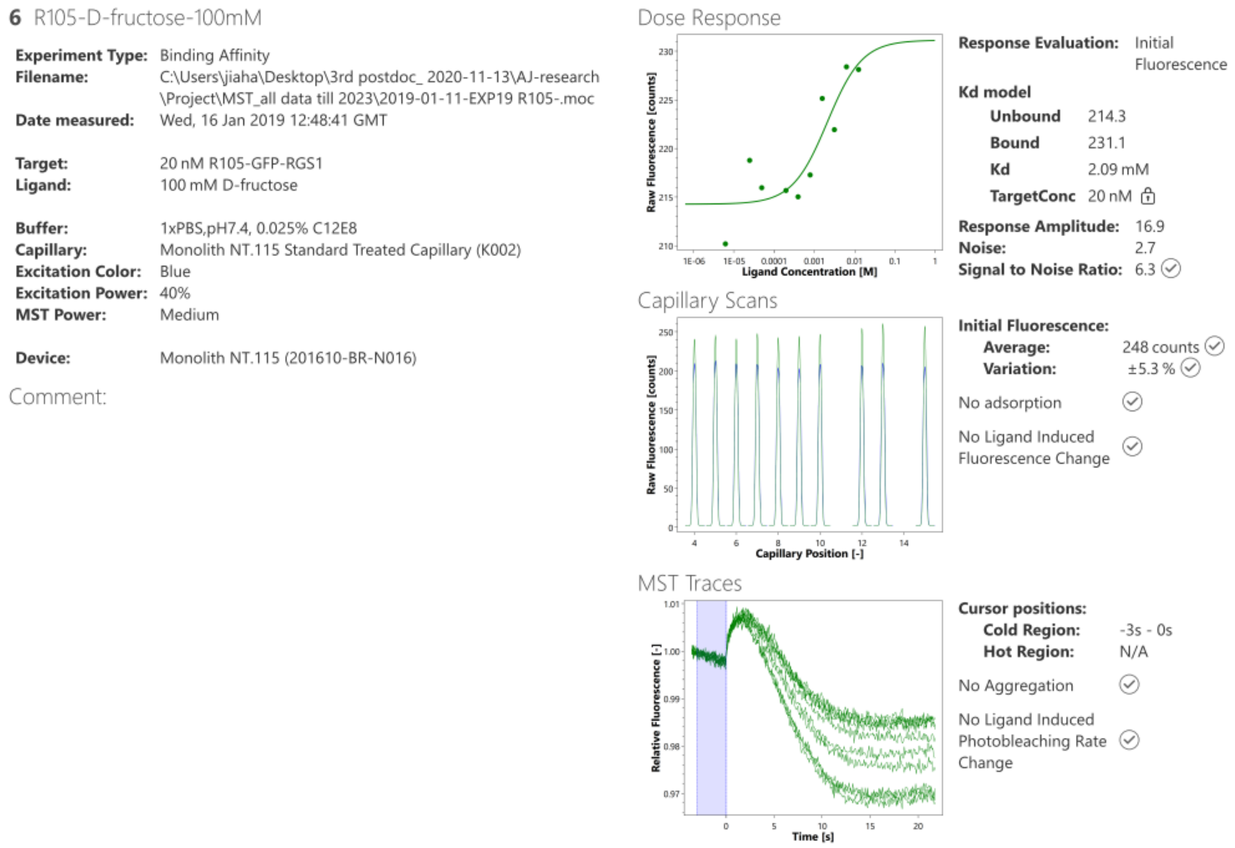


**Figure S4 (continued)**

**Binding profiles of tested ligands to GFP–RGS1 determined by microscale thermophoresis (MST).**

Assay conditions are described in Methods.

**Ligand class: Sugar**

Target: GFP-RGS1

Ligand: Palatinose

Response evaluation: Initial Fluorescence

Signal to Noise: N/A (should be ≥ 5 to conclude target-ligand binding)

Conclusion: No binding

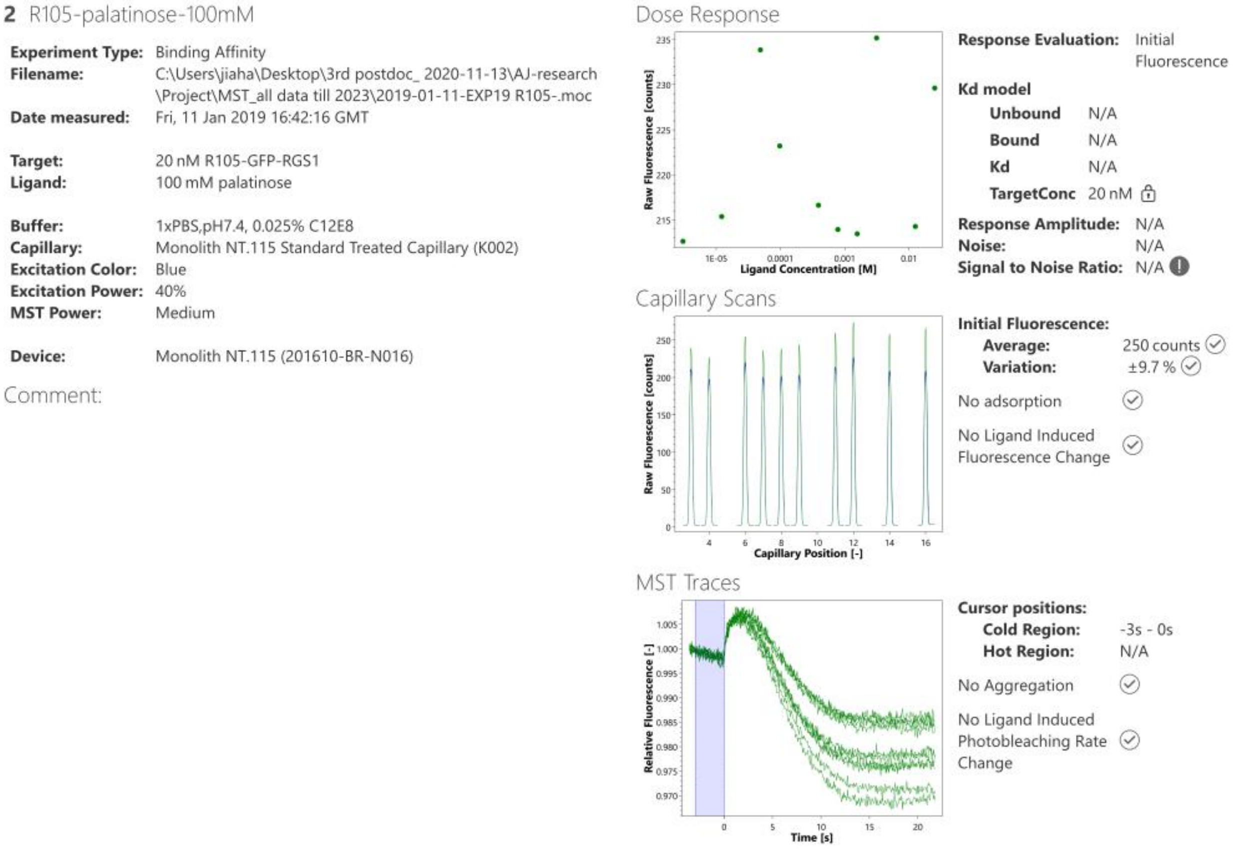


**Figure S4 (continued)**

**Binding profiles of tested ligands to GFP–RGS1 determined by microscale thermophoresis (MST).**

Assay conditions are described in Methods.

**Ligand class: Sugar**

Target: GFP-RGS1

Ligand: Trehalose-6-P

Response evaluation: Initial Fluorescence

Signal to Noise: 14.5 (should be ≥ 5 to conclude target-ligand binding)

Conclusion: binding

Replicate 1: $K_d$ = 3.58mM

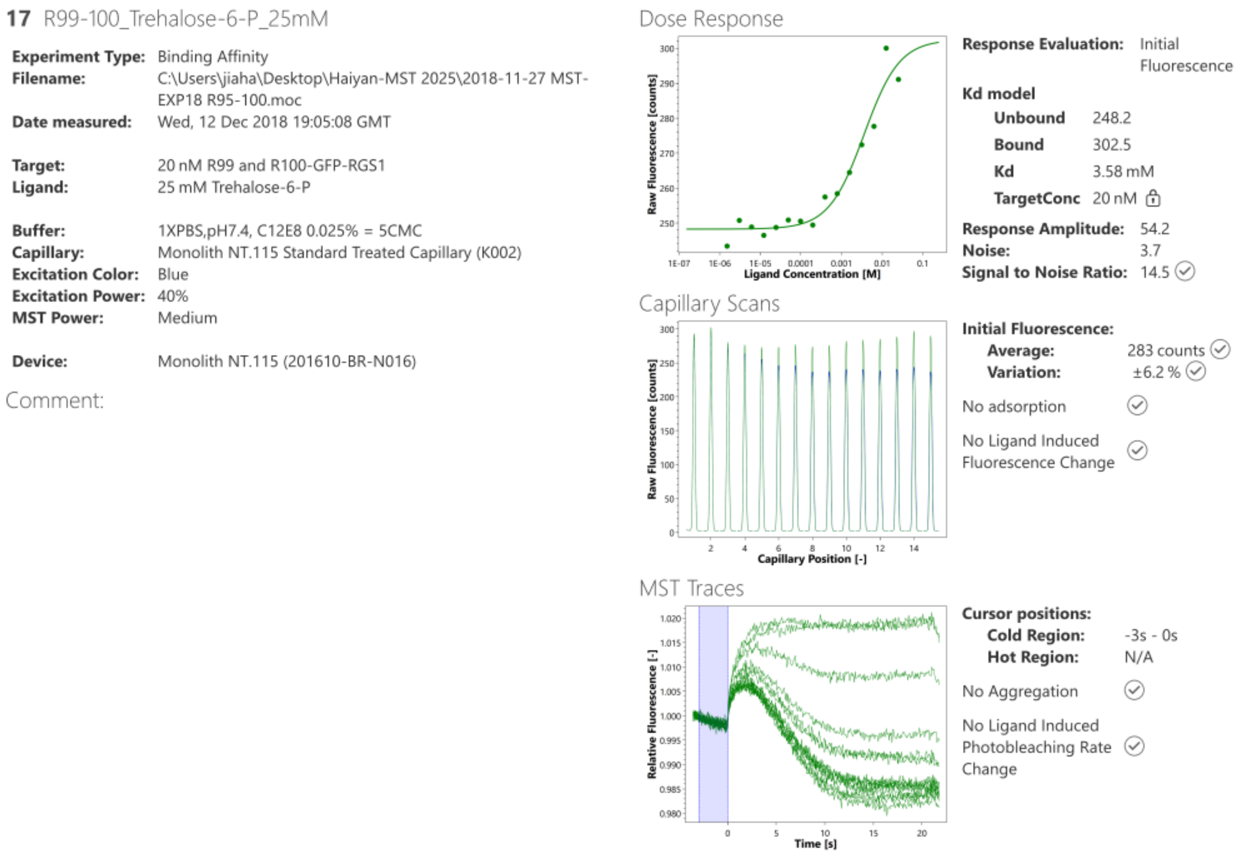


**Figure S4 (continued)**

**Binding profiles of tested ligands to GFP–RGS1 determined by microscale thermophoresis (MST).**

Assay conditions are described in Methods.

**Ligand class: Nucleotide**

Target: GFP-RGS1

Ligand: UDP

Response evaluation: Initial Fluorescence

Signal to Noise: 6.2 (should be ≥ 5 to conclude target-ligand binding)

Conclusion: binding

Replicate 1: $K_d$ = 34.8 μM

**8** R70_UDP 5mM

| | |
|---|---|
| **Experiment Type:** | Binding Affinity |
| **Filename:** | C:\Users\jiaha\Desktop\3rd postdoc_ 2020-11-13\AJ-research \Project\MST data V1\2018-11-04 MST-EXP15_R67-69 and R70-71, and R72-73,R74-R84.moc |
| **Date measured:** | Fri, 12 Oct 2018 14:25:02 GMT |
| **Target:** | 20 nM R70-GFP-RGS1 |
| **Ligand:** | 5 mM UDP |
| **Buffer:** | 1XPBS, pH7.4, 0.1% C12E8 |
| **Capillary:** | Monolith NT.115 Standard Treated Capillary (K002) |
| **Excitation Color:** | Blue |
| **Excitation Power:** | 40% |
| **MST Power:** | Medium |
| **Device:** | Monolith NT.115 (201610-BR-N016) |

Comment:

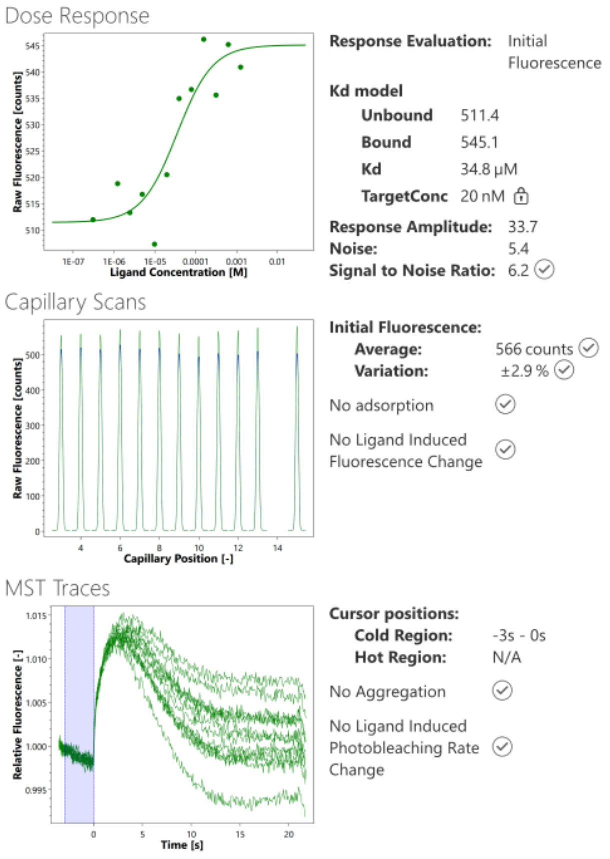


**Figure S4 (continued)**

**Binding profiles of tested ligands to GFP–RGS1 determined by microscale thermophoresis (MST).**

Assay conditions are described in Methods.

**Ligand class: Nucleotide**

Target: GFP-RGS1

Ligand: UDP

Response evaluation: Initial Fluorescence

Signal to Noise: 5.7 (should be ≥ 5 to conclude target-ligand binding)

Conclusion: binding

Replicate 2: $K_d$ = 34.9 μM

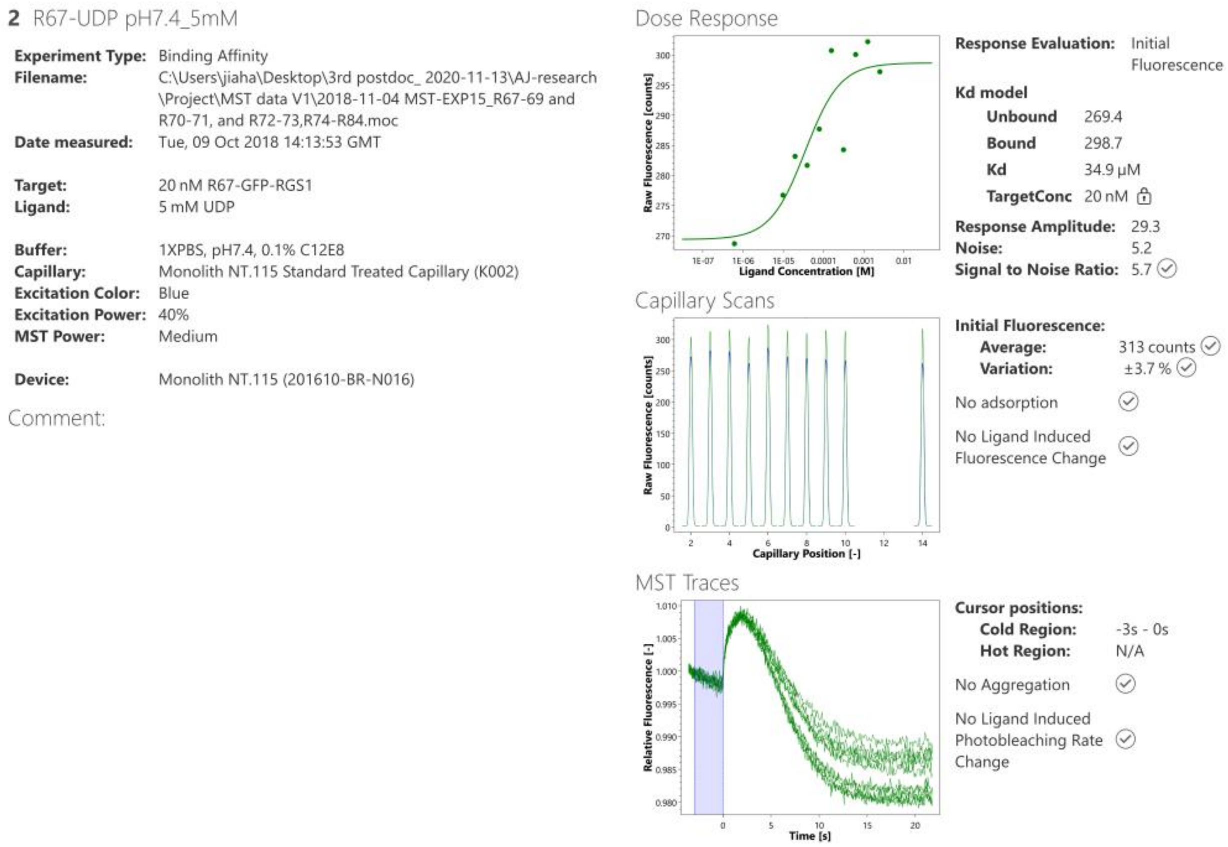


**Figure S4 (continued)**

**Binding profiles of tested ligands to GFP–RGS1 determined by microscale thermophoresis (MST).**

Assay conditions are described in Methods.

**Ligand class: Nucleotide**

Target: GFP-RGS1

Ligand: UDP

Response evaluation: Initial Fluorescence

Signal to Noise: 5.4 (should be ≥ 5 to conclude target-ligand binding)

Conclusion: binding

Replicate 3: $K_d$ = 28.6 μM

**Average $K_d$ = 32.8 ± 3.6 μM**

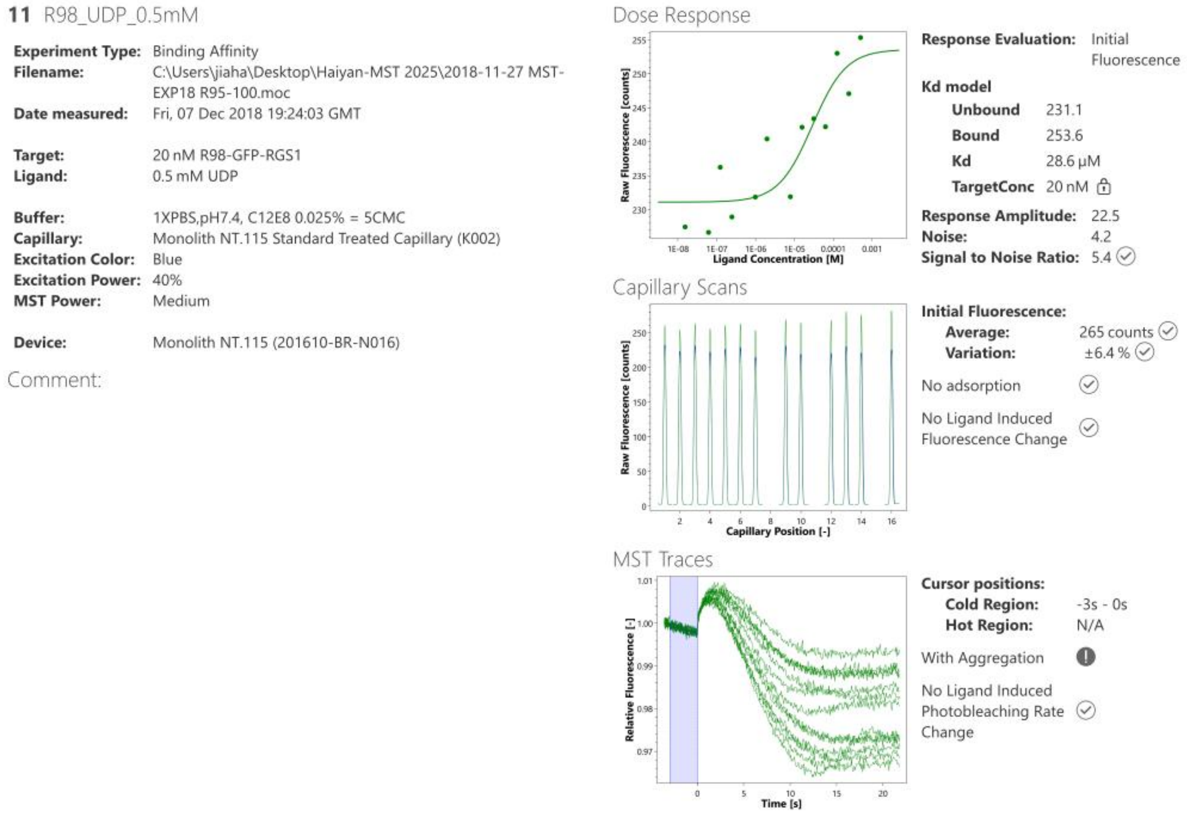


**Figure S4 (continued)**

**Binding profiles of tested ligands to GFP–RGS1 determined by microscale thermophoresis (MST).**

Assay conditions are described in Methods.

**Ligand class: Nucleotide**

Target: GFP-RGS1

Ligand: ADP

Response evaluation: Initial Fluorescence

Signal to Noise: 5.3 (should be ≥ 5 to conclude target-ligand binding)

Conclusion: binding

$K_d$ = 126 μM

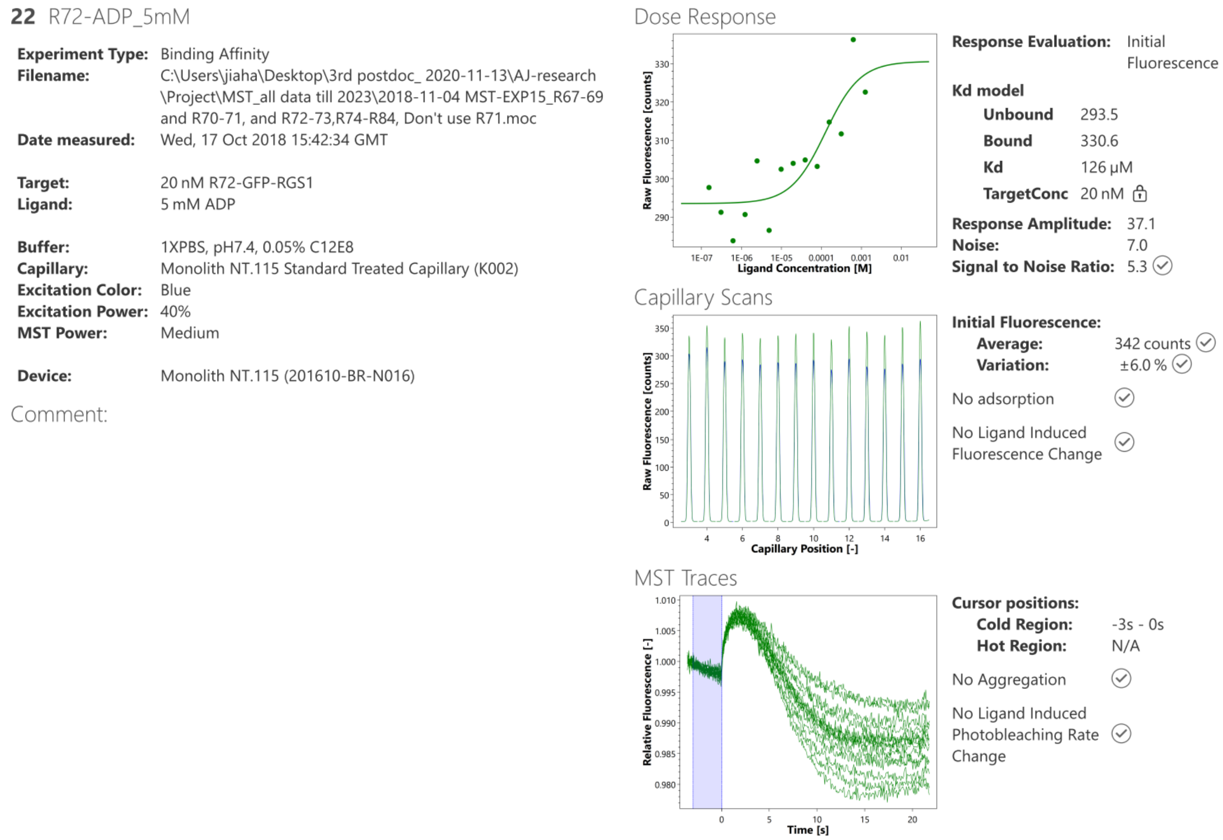


**Figure S4 (continued)**

**Binding profiles of tested ligands to GFP–RGS1 determined by microscale thermophoresis (MST).**

Assay conditions are described in Methods.

**Ligand class: Nucleotide**

Target: GFP-RGS1

Ligand: ATP

Response evaluation: Initial Fluorescence

Signal to Noise: N/A (should be ≥ 5 to conclude target-ligand binding)

Conclusion: No binding

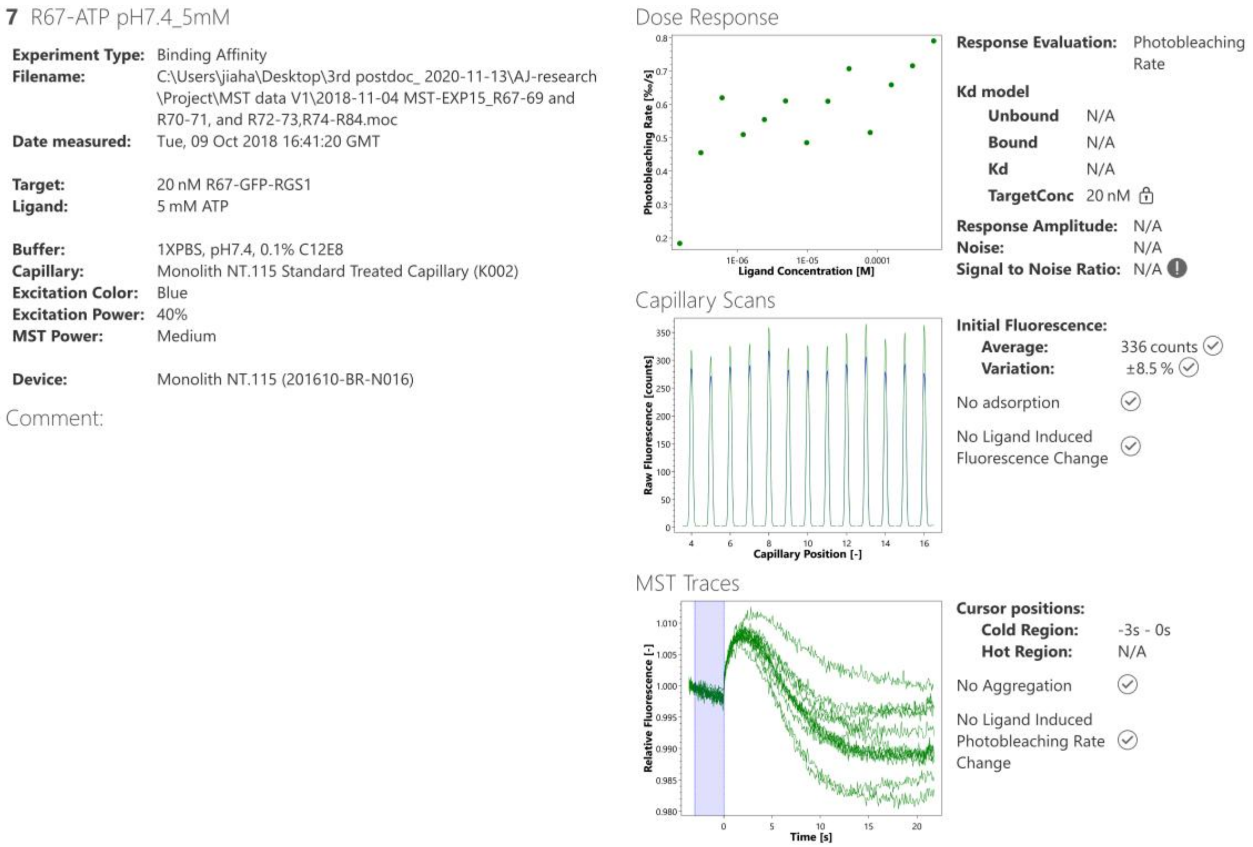


**Figure S4 (continued)**

**Binding profiles of tested ligands to GFP–RGS1 determined by microscale thermophoresis (MST).**

Assay conditions are described in Methods.

**Ligand class: Nucleotide**

Target: GFP-RGS1

Ligand: GTP

Response evaluation: Initial Fluorescence

Signal to Noise: N/A (should be ≥ 5 to conclude target-ligand binding)

Conclusion: No binding

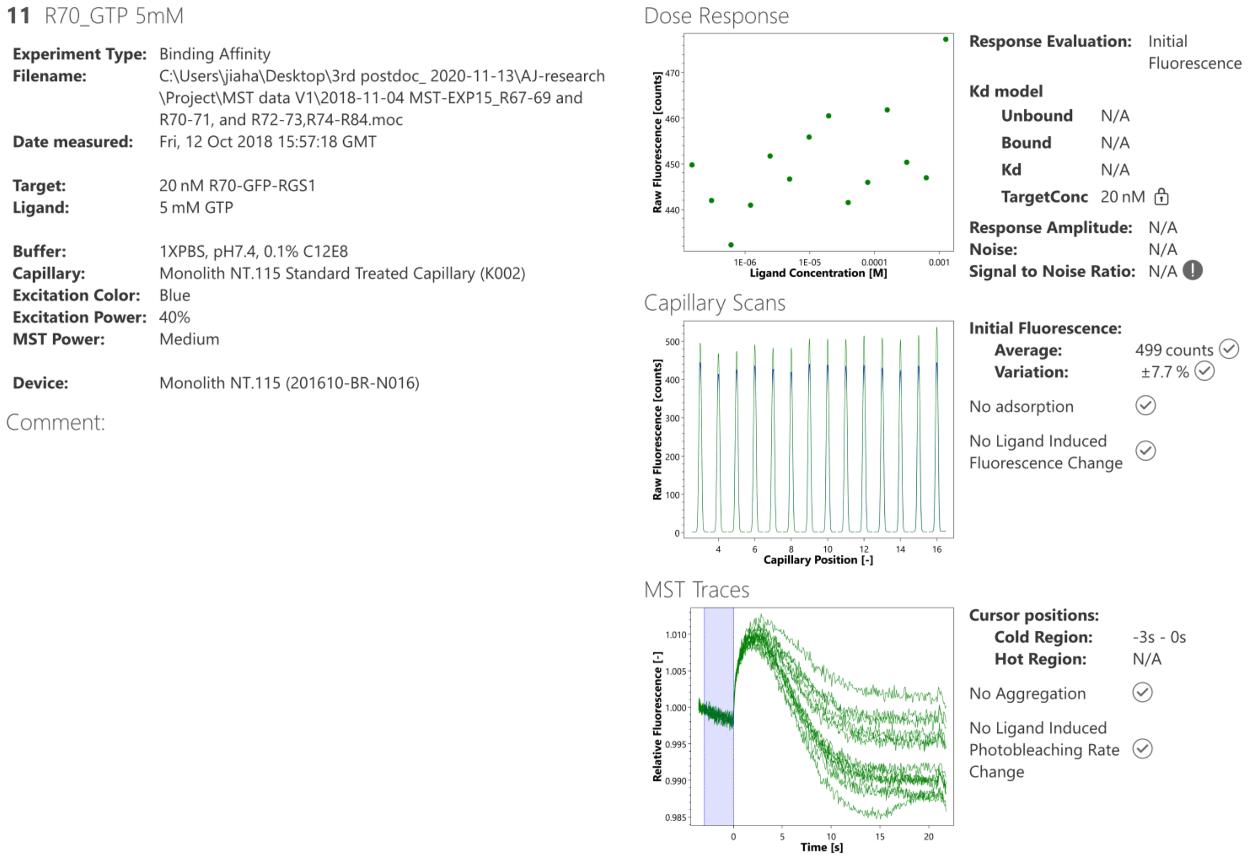


**Figure S4 (continued)**

**Binding profiles of tested ligands to GFP–RGS1 determined by microscale thermophoresis (MST).**

Assay conditions are described in Methods.

**Ligand class: Nucleotide**

Target: GFP-RGS1

Ligand: UTP

Response evaluation: Initial Fluorescence

Signal to Noise: 3.2 (should be ≥ 5 to conclude target-ligand binding)

Conclusion: No binding

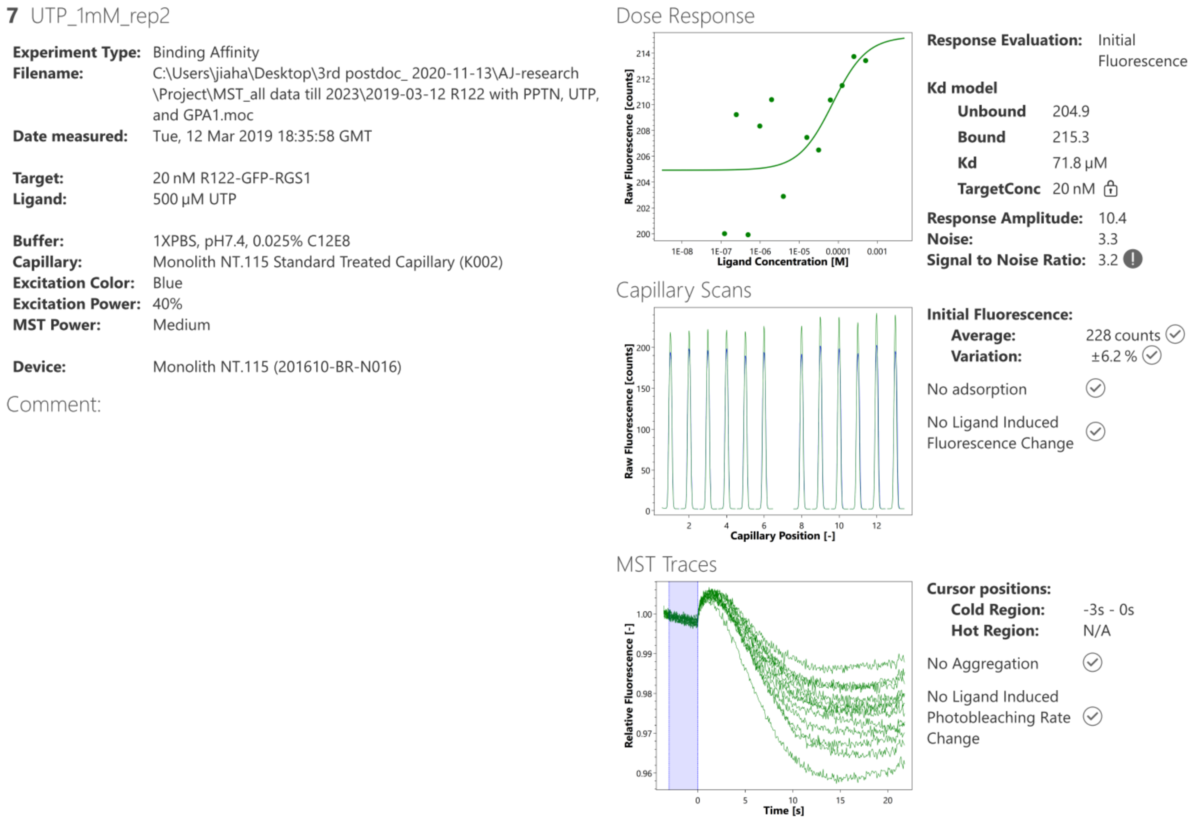


**Figure S4 (continued)**

**Binding profiles of tested ligands to GFP–RGS1 determined by microscale thermophoresis (MST).**

Assay conditions are described in Methods.

**Ligand class: nucleotide sugar**

Target: GFP-RGS1

Ligand: UDPG

Response evaluation: Initial Fluorescence

Rep1

Signal to Noise: 7.1 (should be ≥ 5 to conclude target-ligand binding)

Conclusion: binding

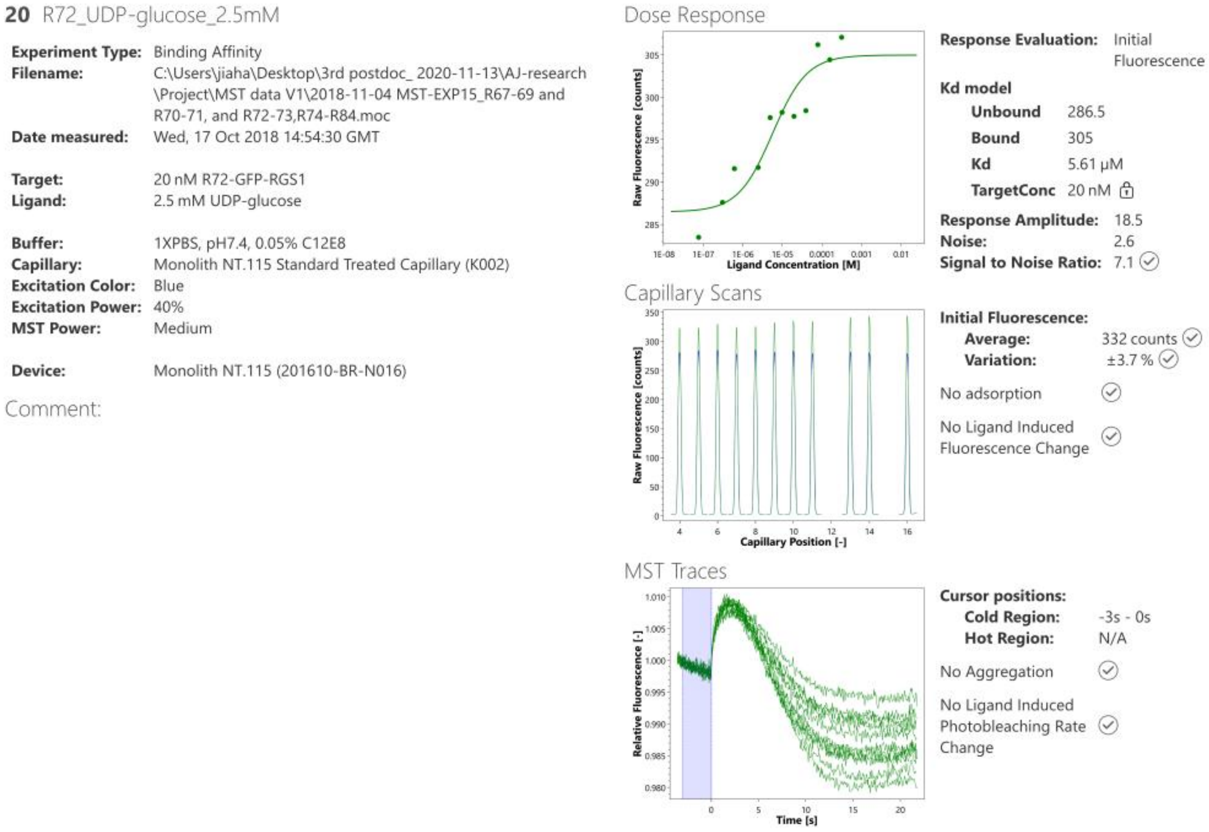


**Figure S4 (continued)**

**Binding profiles of tested ligands to GFP–RGS1 determined by microscale thermophoresis (MST).**

Assay conditions are described in Methods.

**Ligand class: nucleotide sugar**

Target: GFP-RGS1

Ligand: UDPG

Response evaluation: Initial Fluorescence

Rep2

Signal to Noise: 5 (should be ≥ 5 to conclude target-ligand binding)

Conclusion: binding

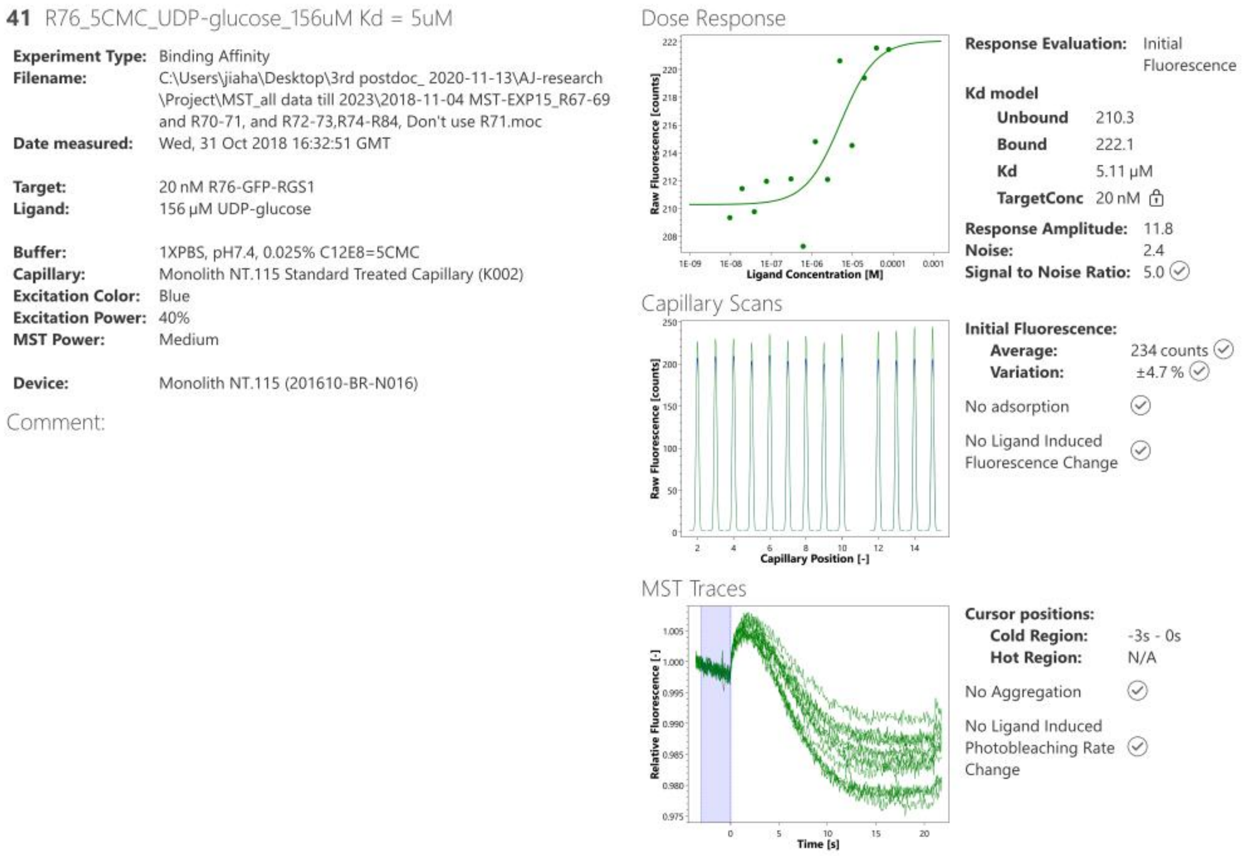


**Figure S4 (continued)**

**Binding profiles of tested ligands to GFP–RGS1 determined by microscale thermophoresis (MST).**

Assay conditions are described in methods.

**Ligand class: nucleotide sugar**

Target: GFP-RGS1

Ligand: UDPG

Response evaluation: Initial Fluorescence

Rep3

Signal to Noise: 5.1 (should be ≥ 5 to conclude target-ligand binding)

Conclusion: binding

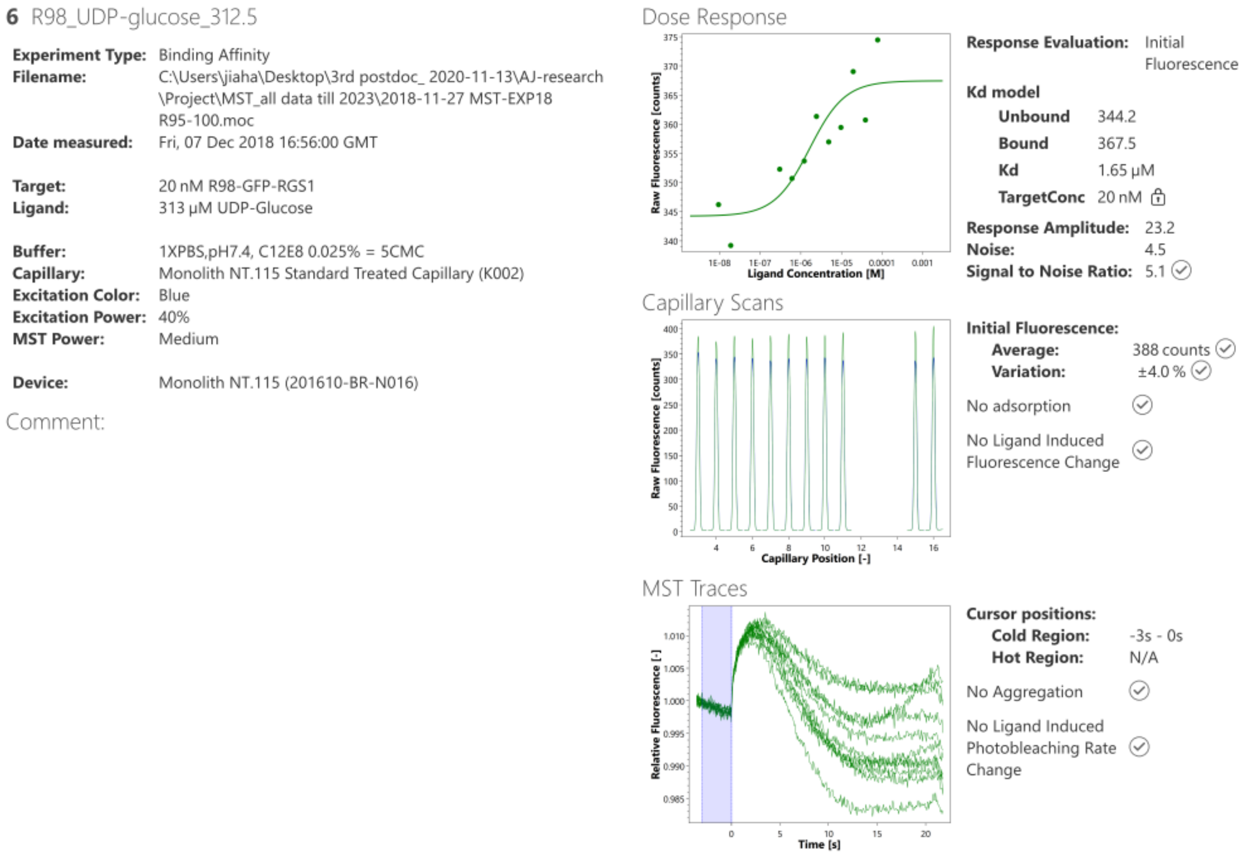


**Figure S4 (continued)**

**Binding profiles of tested ligands to GFP–RGS1 determined by microscale thermophoresis (MST).**

Assay conditions are described in Methods.

**Ligand class: nucleotide sugar**

Target: GFP-RGS1

Ligand: UDPG

Response evaluation: Initial Fluorescence

Rep4

Signal to Noise: 5.8 (should be ≥ 5 to conclude target-ligand binding)

Conclusion: binding

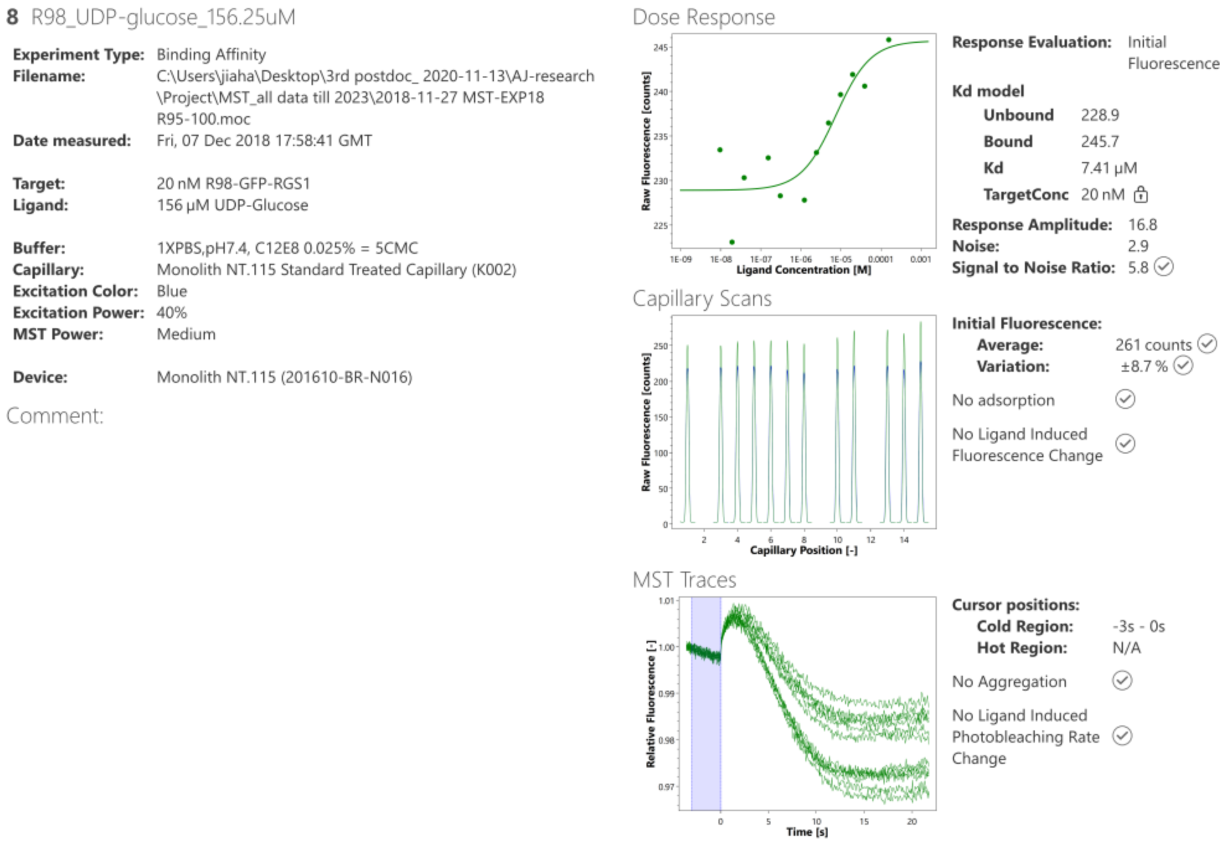


**Figure S4 (continued)**

**Binding profiles of tested ligands to GFP–RGS1 determined by microscale thermophoresis (MST).**

Assay conditions are described in Methods.

**Ligand class: nucleotide sugar**

Target: GFP-RGS1

Ligand: UDPG

Response evaluation: Initial Fluorescence

Rep5

Signal to Noise: 6.5 (should be ≥ 5 to conclude target-ligand binding)

Conclusion: binding

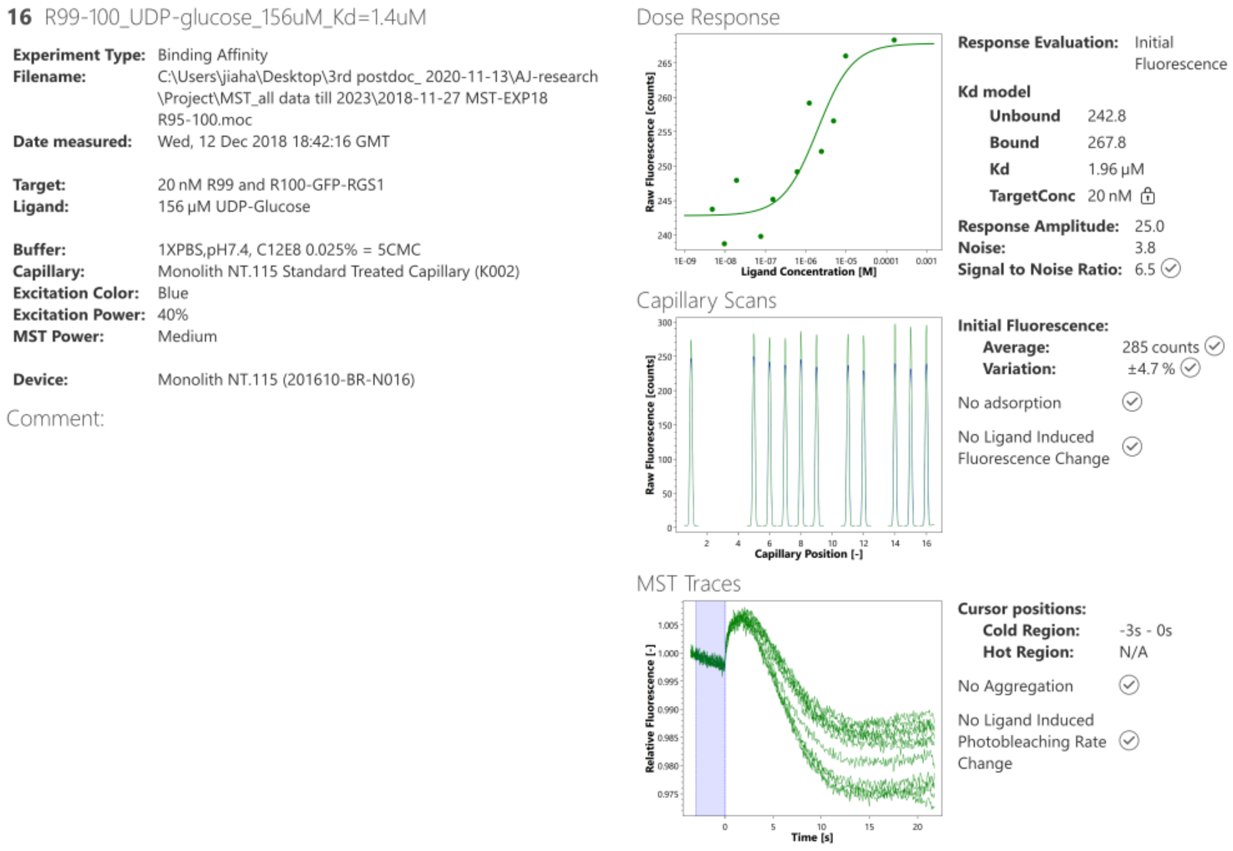


**Figure S4 (continued)**

**Binding profiles of tested ligands to GFP–RGS1 determined by microscale thermophoresis (MST).**

Assay conditions are described in Methods.

**Ligand class: P2Y14 agonist**

Target: GFP-RGS1

Ligand: MRS2905

Response evaluation: Initial Fluorescence

Signal to Noise: 6.7 (should be ≥ 5 to conclude target-ligand binding)

Conclusion: binding

$K_d$ = 1.24 μM

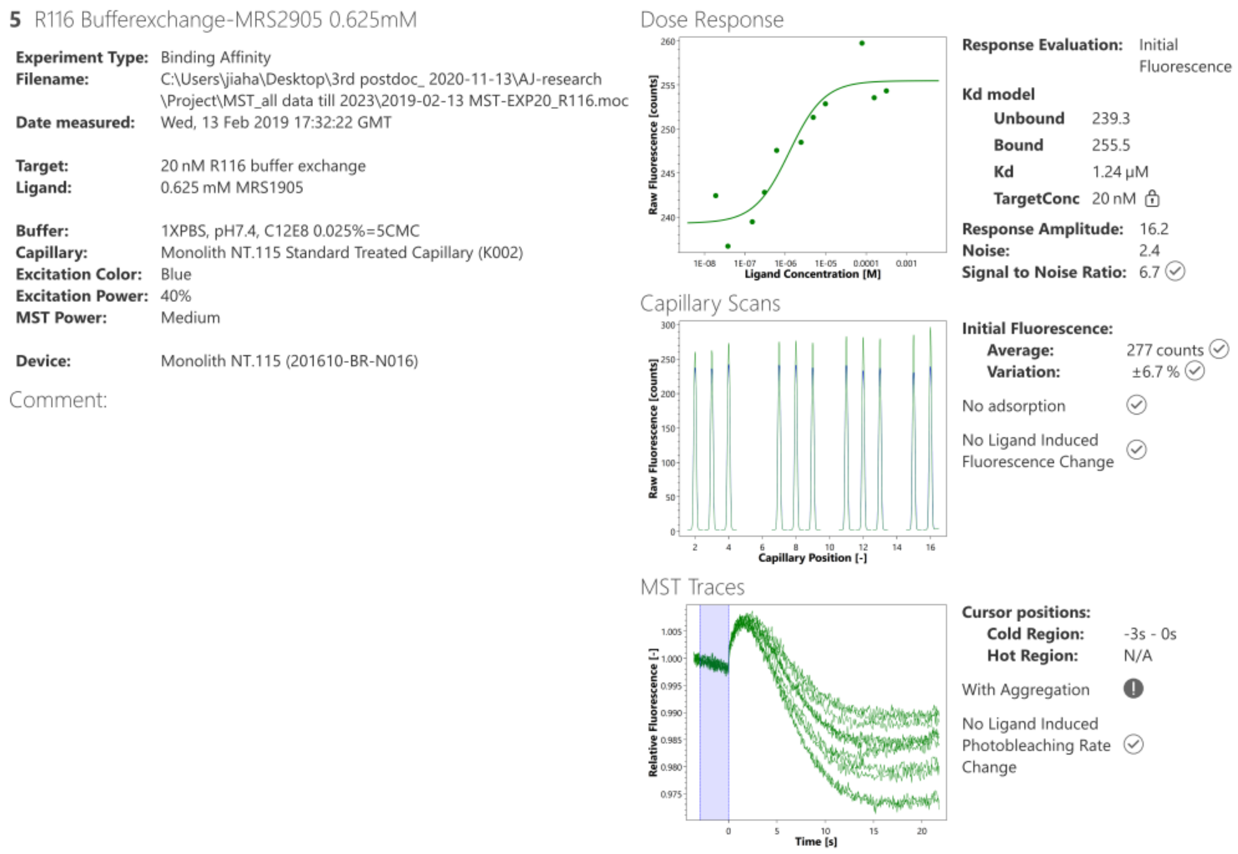


**Figure S4 (continued)**

**Binding profiles of tested ligands to GFP–RGS1 determined by microscale thermophoresis (MST).**

Assay conditions are described in Methods.

**Figure S4 continued.**

**Ligand class: P2Y14 antagonist**

Target: GFP-RGS1

Ligand: PPTN

Response evaluation: Initial Fluorescence

Signal to Noise: 11.8 (should be ≥ 5 to conclude target-ligand binding)

Conclusion: binding

$K_d$ = 14 μM

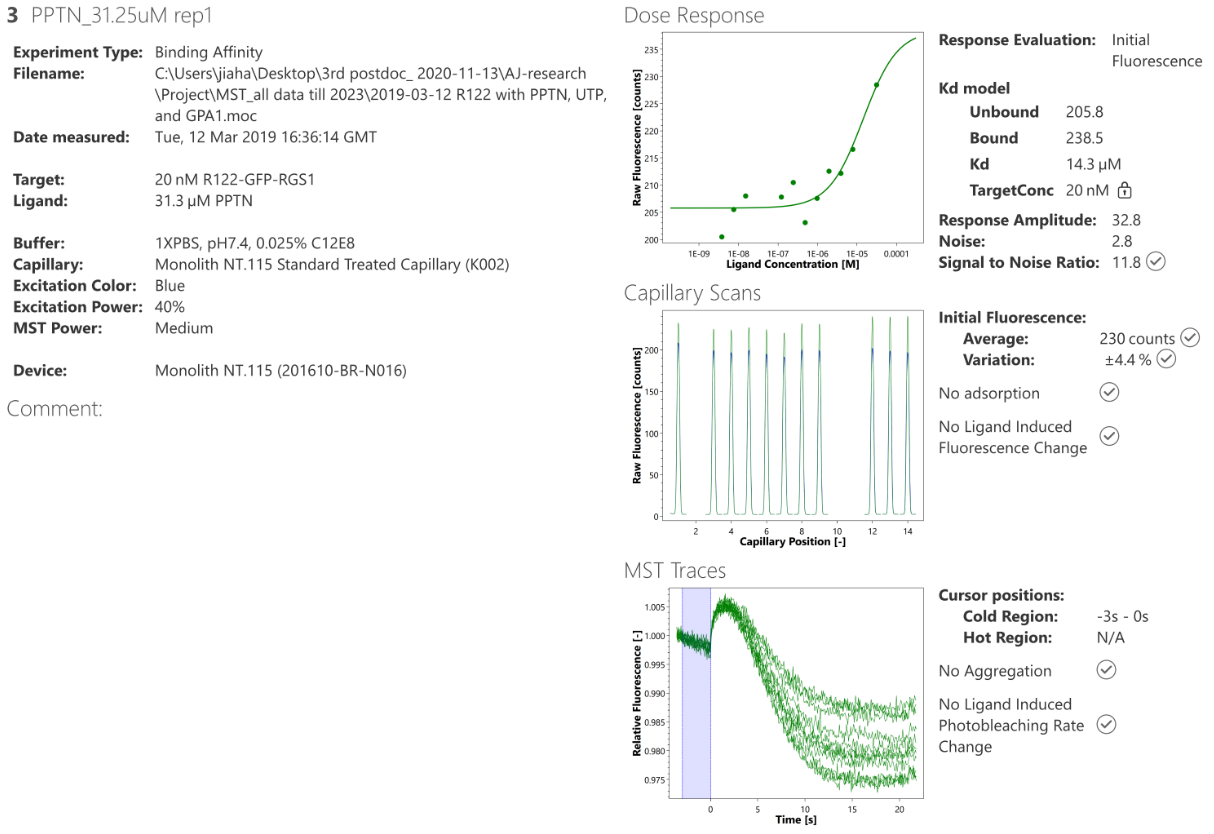


**Figure S4 (continued)**

**Binding profiles of tested ligands to GFP–RGS1 determined by microscale thermophoresis (MST).**

Assay conditions are described in Methods.

**Ligand class: P2Y$_{14}$ antagonist**

Target: GFP-RGS1

Ligand: MRS4574

Response evaluation: Initial Fluorescence

Signal to Noise: N/A (should be ≥ 5 to conclude target-ligand binding)

Conclusion: no binding

**17** 1.MRS4574_250uM_rep2

**Experiment Type:** Binding Affinity
**Filename:** C:\Users\jiaha\Desktop\3rd postdoc_ 2020-11-13\AJ-research \Project\MST_all data till 2023\2019-04-02 MST-R123 MRS4574-4578.moc
**Date measured:** Tue, 02 Apr 2019 22:40:17 GMT

**Target:** 20 nM R123-GFP-RGS1
**Ligand:** 250 μM MRS4574

**Buffer:** 1XPBS,pH7.4, C12E8 0.025%
**Capillary:** Monolith NT.115 Standard Treated Capillary (K002)
**Excitation Color:** Blue
**Excitation Power:** 40%
**MST Power:** Medium

**Device:** Monolith NT.115 (201610-BR-N016)

Comment:

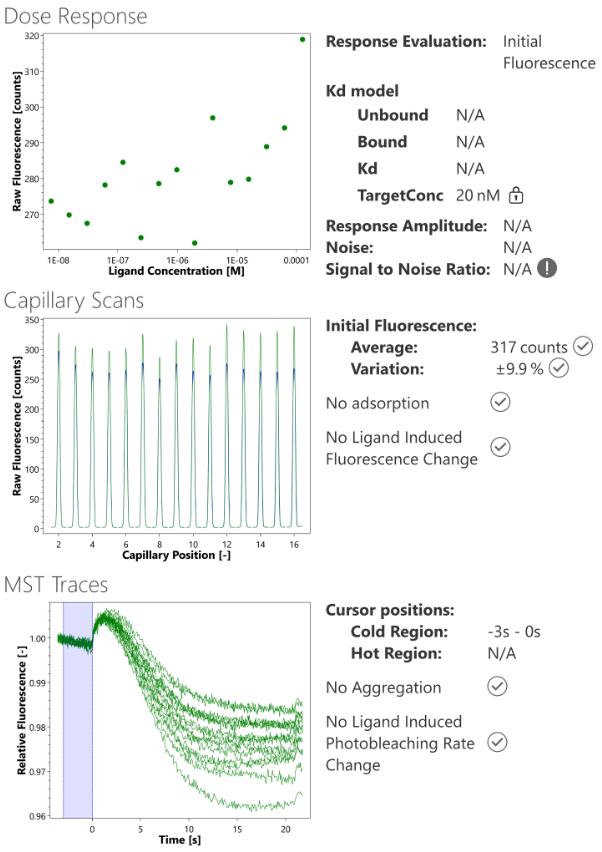


**Figure S4 (continued)**

**Binding profiles of tested ligands to GFP–RGS1 determined by microscale thermophoresis (MST).**

Assay conditions are described in Methods.

**Ligand class: P2Y14 antagonist**

Target: GFP-RGS1

Ligand: MRS4575

Response evaluation: Initial Fluorescence

Signal to Noise: 3.3 (should be ≥ 5 to conclude target-ligand binding)

Conclusion: no binding

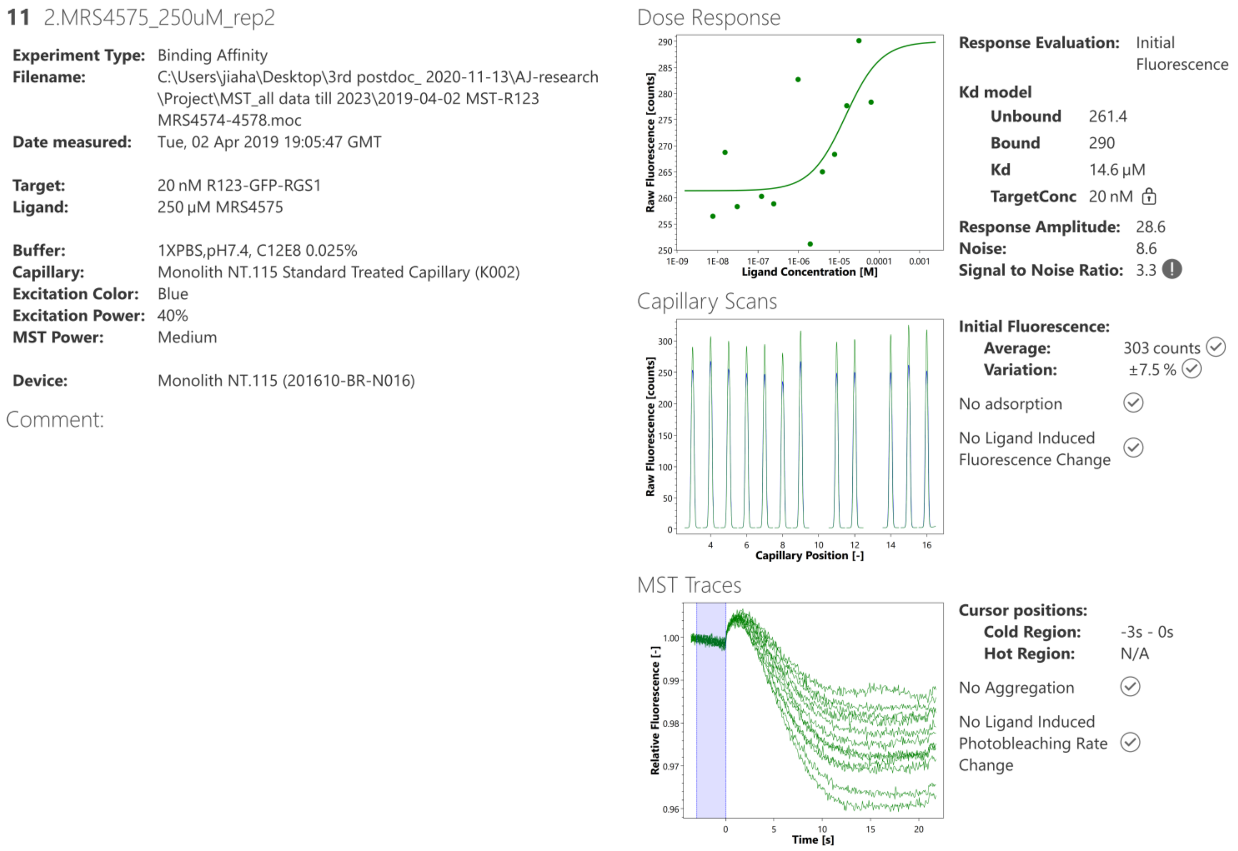


**Figure S4 (continued)**

**Binding profiles of tested ligands to GFP–RGS1 determined by microscale thermophoresis (MST).**

Assay conditions are described in Methods.

**Ligand class: P2Y$_{14}$ antagonist**

Target: GFP-RGS1

Ligand: MRS4576

Response evaluation: Initial Fluorescence

Signal to Noise: N/A (should be ≥ 5 to conclude target-ligand binding)

Conclusion: no binding

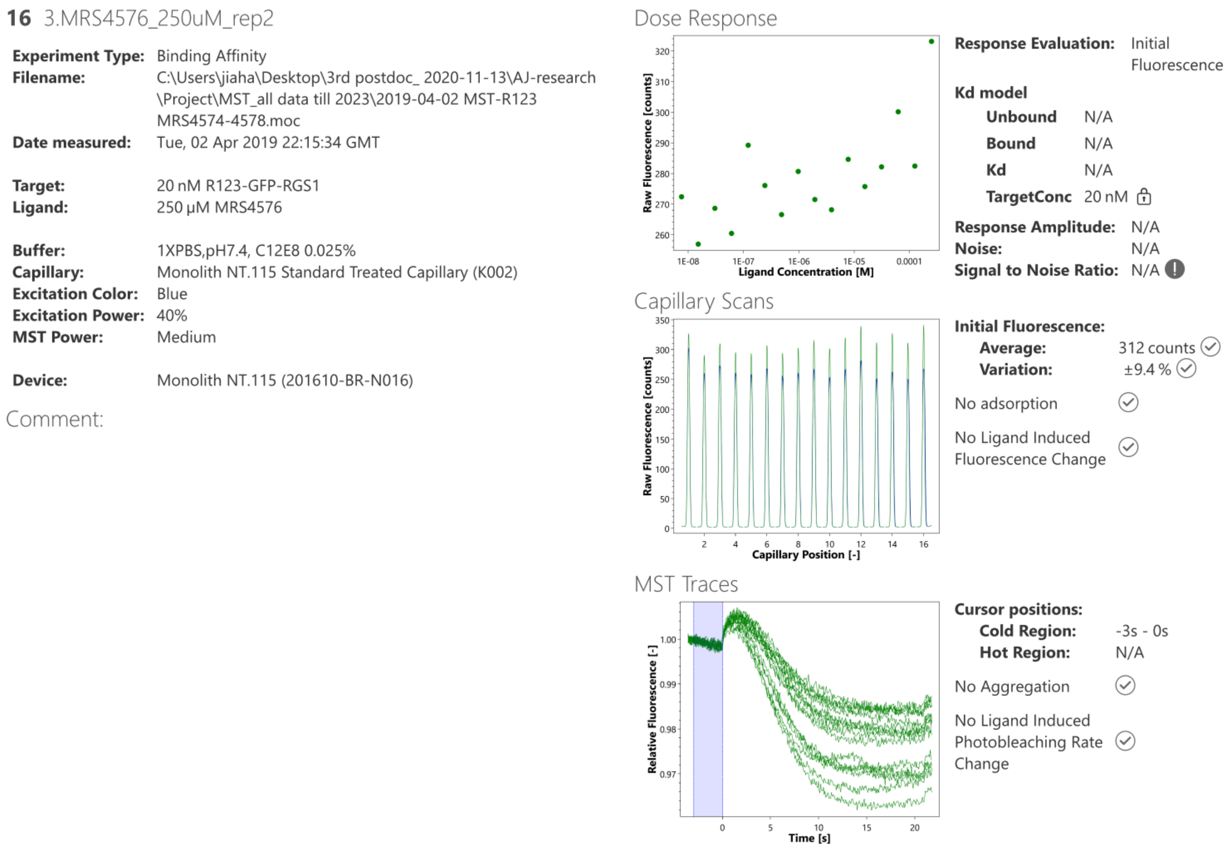


**Figure S4 (continued)**

**Binding profiles of tested ligands to GFP–RGS1 determined by microscale thermophoresis (MST).**

Assay conditions are described in Methods.

**Ligand class: $P2Y_{14}$ antagonist**

Target: GFP-RGS1

Ligand: MRS4577

Response evaluation: Initial Fluorescence

Signal to Noise: 5.7 (should be ≥ 5 to conclude target-ligand binding)

Conclusion: binding

$K_d$ = 5.21 μM

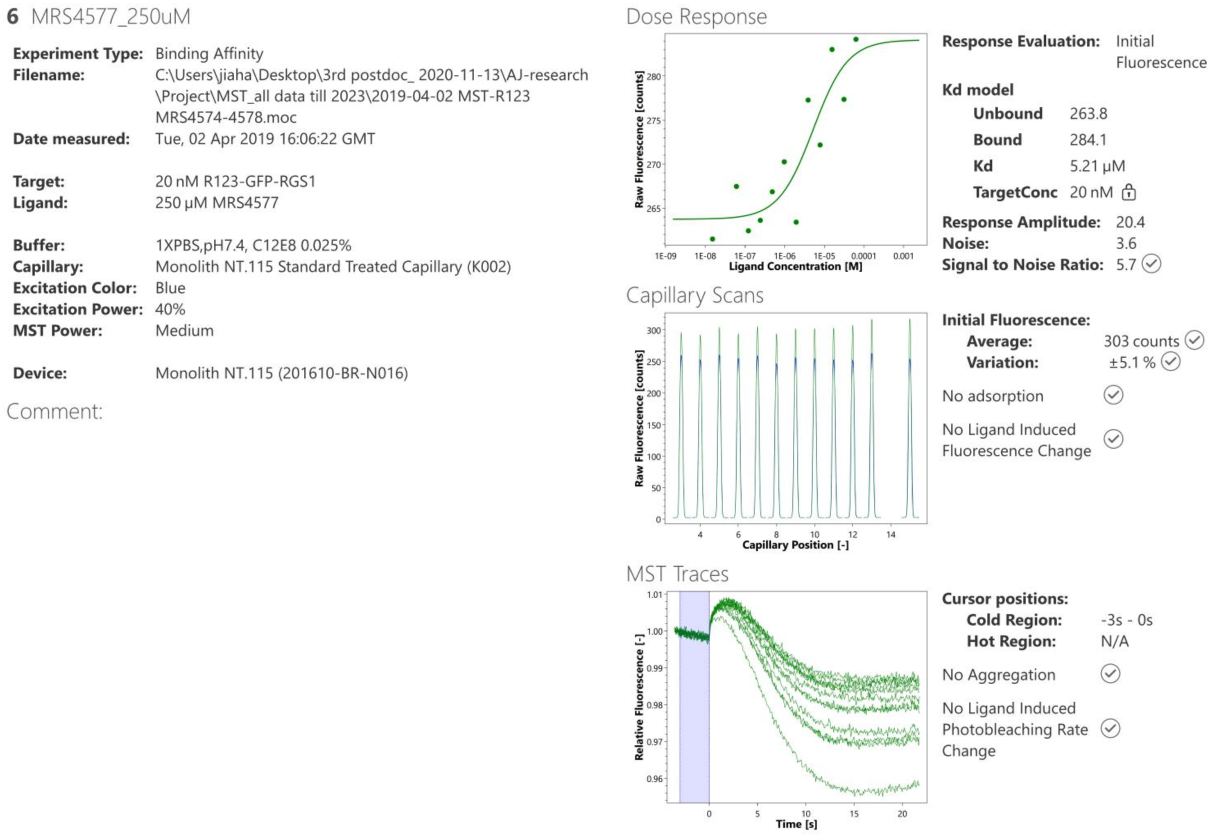


**Figure S4 (continued)**

**Binding profiles of tested ligands to GFP–RGS1 determined by microscale thermophoresis (MST).**

Assay conditions are described in Methods.

**Ligand class: P2Y$_{14}$ antagonist**

Target: GFP-RGS1

Ligand: MRS4578

Response evaluation: Initial Fluorescence

Signal to Noise: 1.3 (should be ≥ 5 to conclude target-ligand binding)

Conclusion: no binding

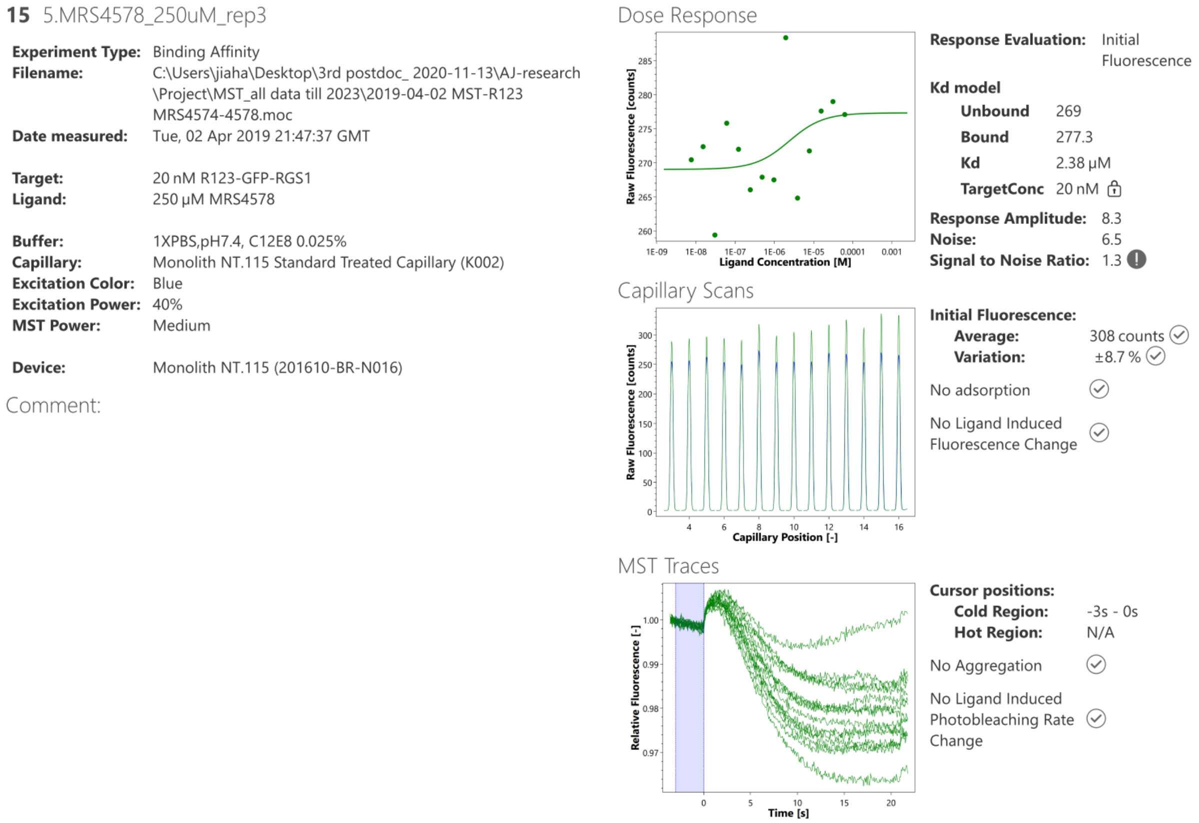


**Figure S4 (continued)**

**Binding profiles of tested ligands to GFP–RGS1 determined by microscale thermophoresis (MST).**

Assay conditions are described in Methods.

Model: RGS1-7TM

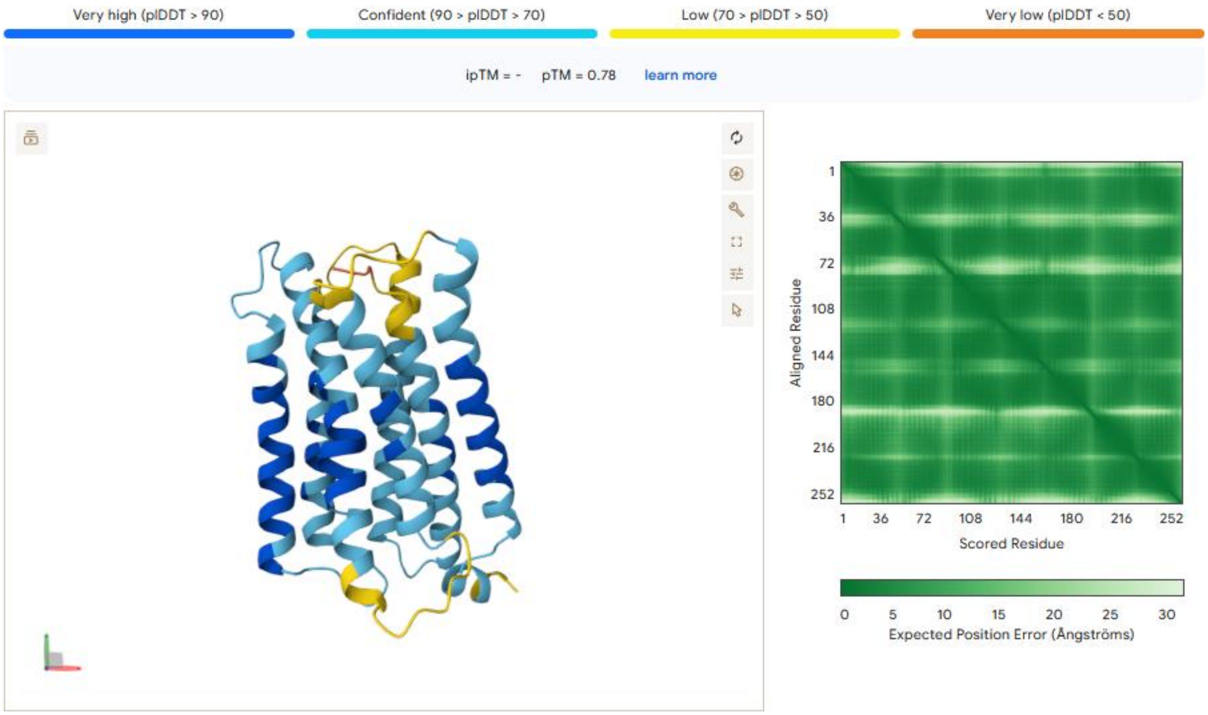


Model: hP2Y14

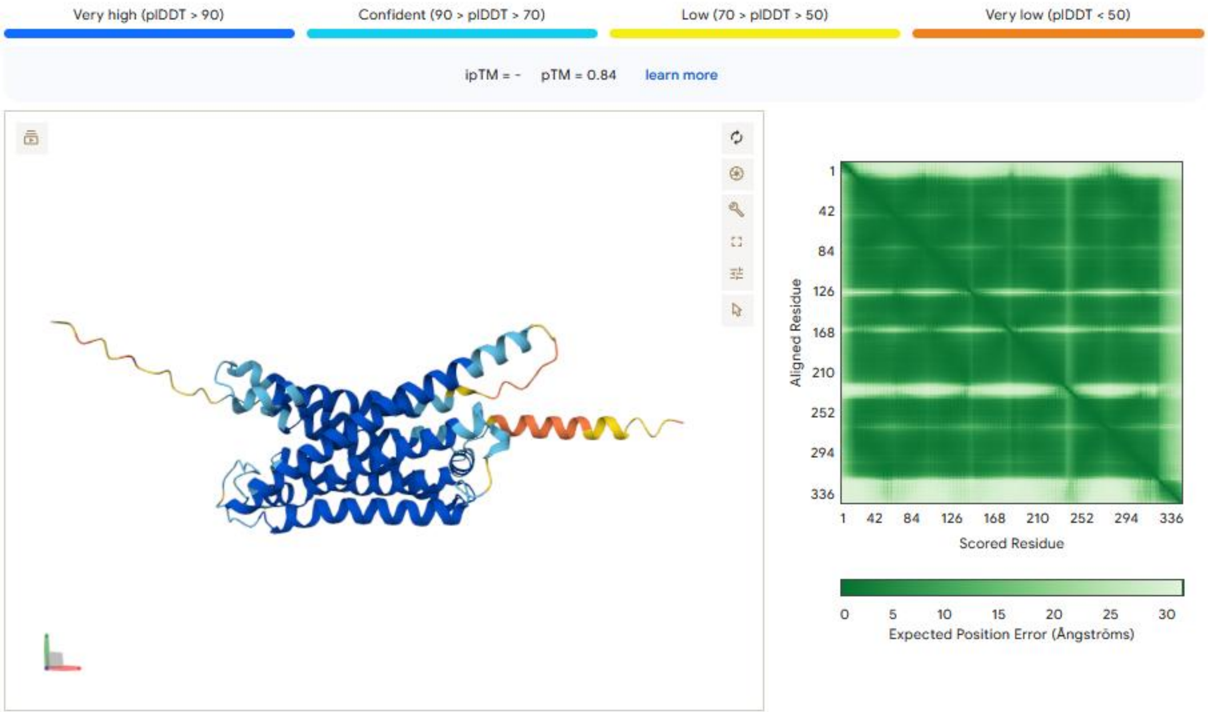

**Figure S5.**

**pLDDT confidence scores of 7TM domain models in Figure 1D.**

Per-residue pLDDT scores for the predicted 7TM structures of RGS1 and hP2Y14, generated by AlphaFold3, are shown. High scores indicate high model confidence for the corresponding residues.

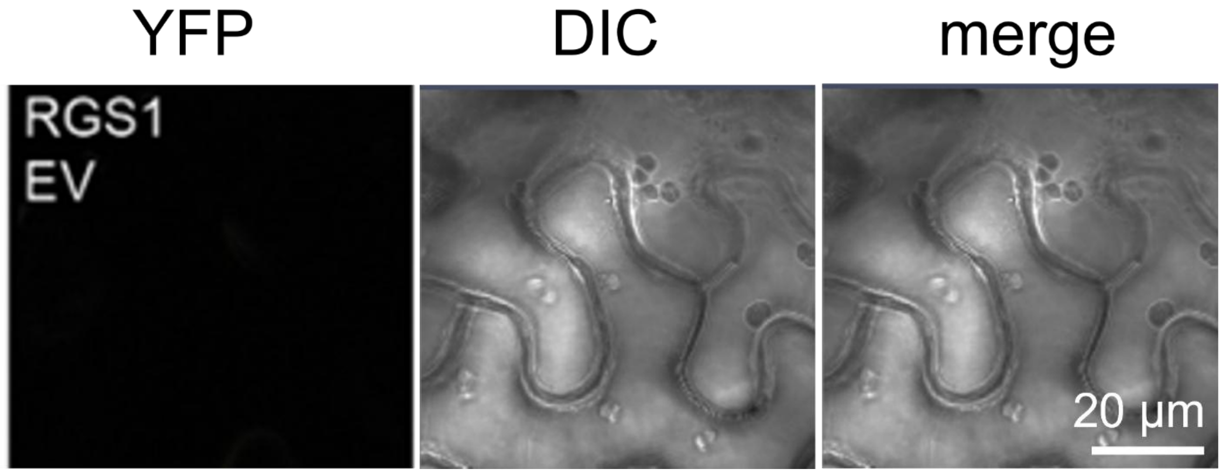


**Figure S6.**

**BiFC image of RGS1 and EV ctrl.**

RGS1-nYFP and cYFP-HA empty vector (EV) were transiently expressed in *N. benthamiana* leaves via *Agrobacterium* infiltration. Leaf tissues were imaged by confocal microscopy two days post-infiltration.

RGS1 and SUS1: Rep1

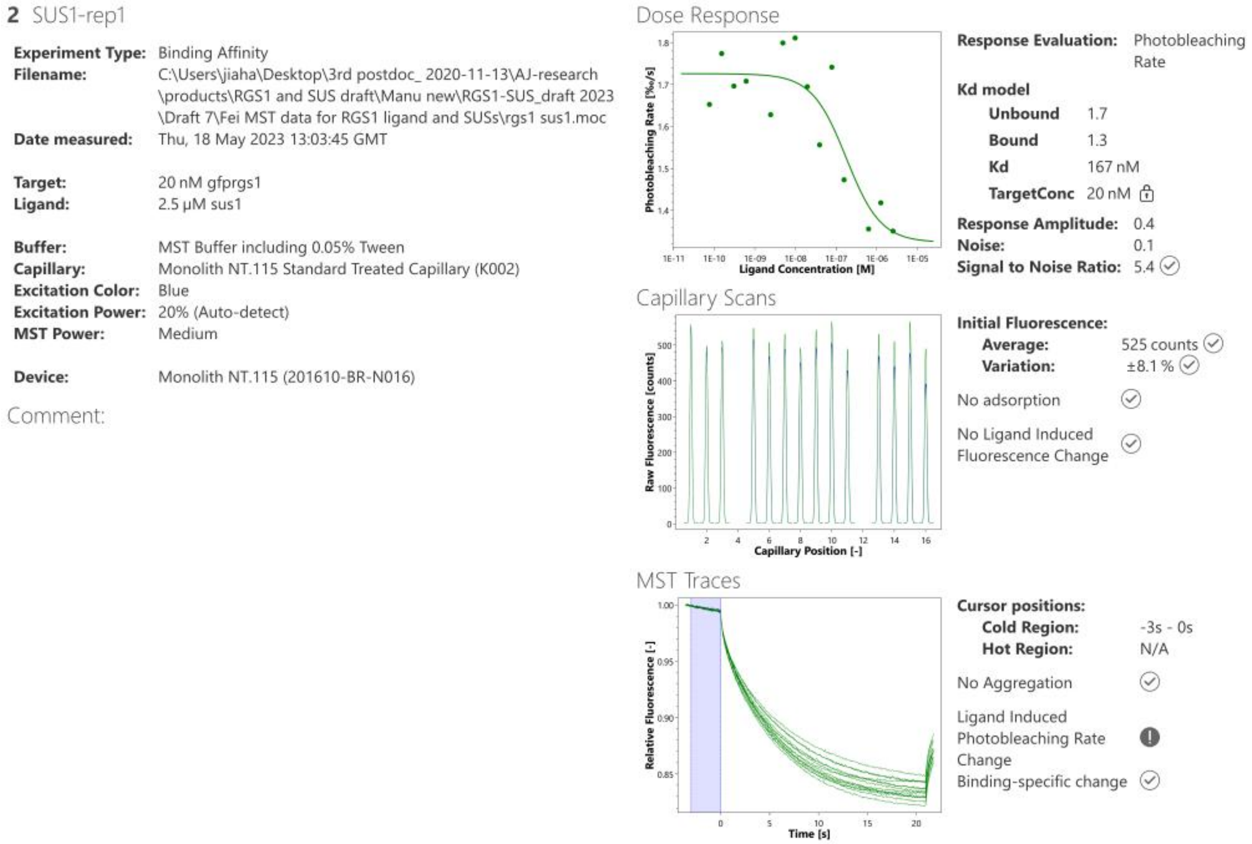


**Figure S7.**

**sfGFP-RGS1 and SUS1 binding profile in MST assay**

Microscale thermophoresis (MST) binding analysis of sfGFP-RGS1 (ΔC-tail) with His-SUS1. Serial dilutions of His-SUS1 were titrated against a fixed concentration of sfGFP-RGS1 (ΔC-tail). Changes in thermophoretic mobility were measured to generate the binding profile, and the dissociation constant ($K_d$) was determined by fitting the data to a standard binding model.

RGS1 and SUS1: Rep2

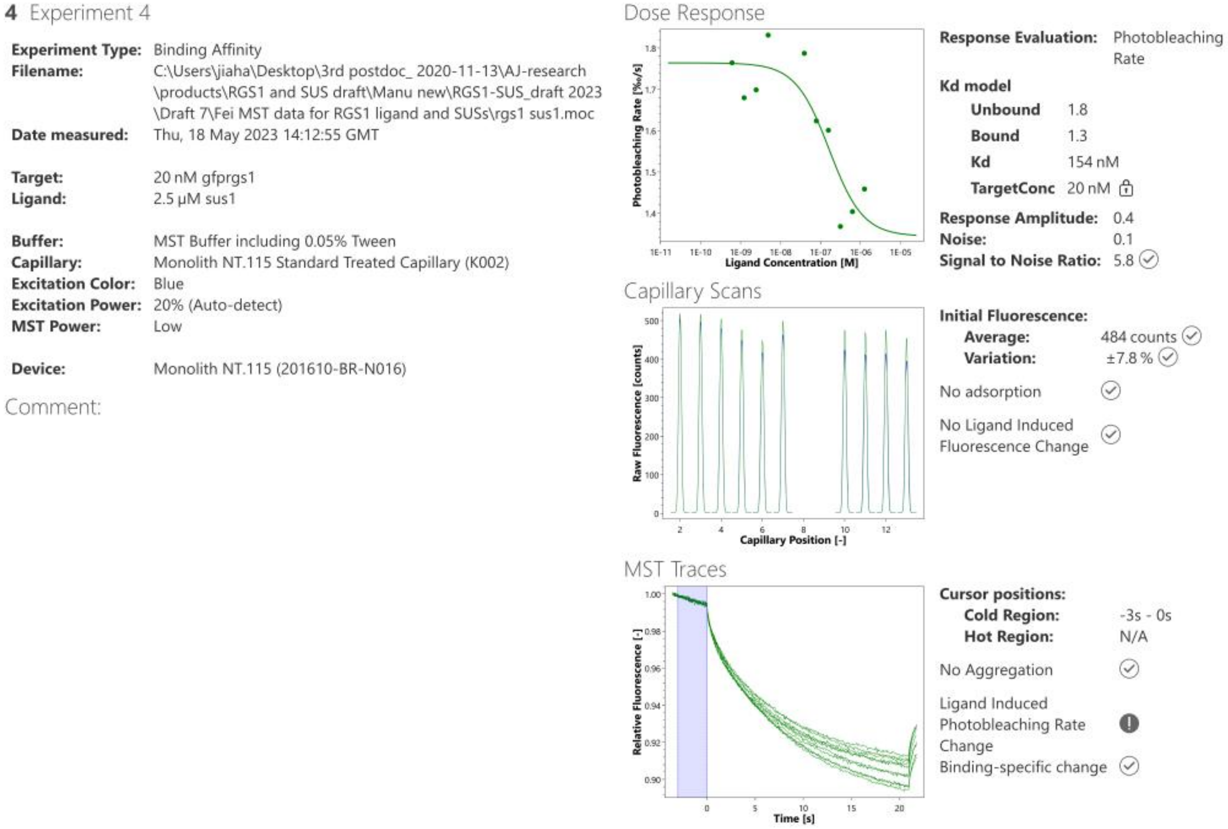


**Figure S7 (continued).**

**sfGFP-RGS1 and SUS1 binding profile in MST assay**

Microscale thermophoresis (MST) binding analysis of sfGFP-RGS1 (ΔC-tail) with His-SUS1. Serial dilutions of His-SUS1 were titrated against a fixed concentration of sfGFP-RGS1 (ΔC-tail). Changes in thermophoretic mobility were measured to generate the binding profile, and the dissociation constant ($K_d$) was determined by fitting the data to a standard binding model.

RGS1 and SUS4: Rep1

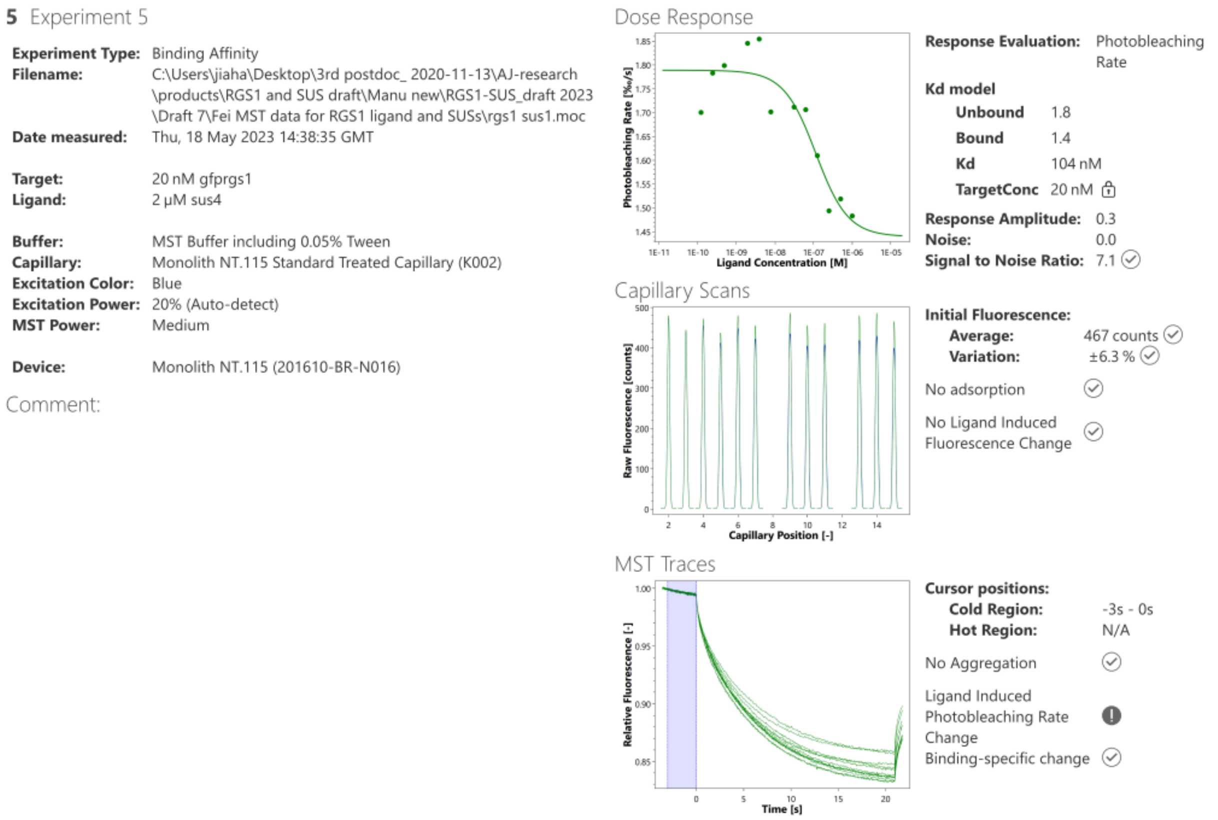


**Figure S8.**

**sfGFP-RGS1 and SUS4 binding profile in MST assay**

Microscale thermophoresis (MST) binding analysis of sfGFP-RGS1 (ΔC-tail) with His-SUS4. Serial dilutions of His-SUS4 were titrated against a fixed concentration of sfGFP-RGS1 (ΔC-tail). Changes in thermophoretic mobility were measured to generate the binding profile, and the dissociation constant ($K_d$) was determined by fitting the data to a standard binding model.

RGS1 and SUS4: Rep2

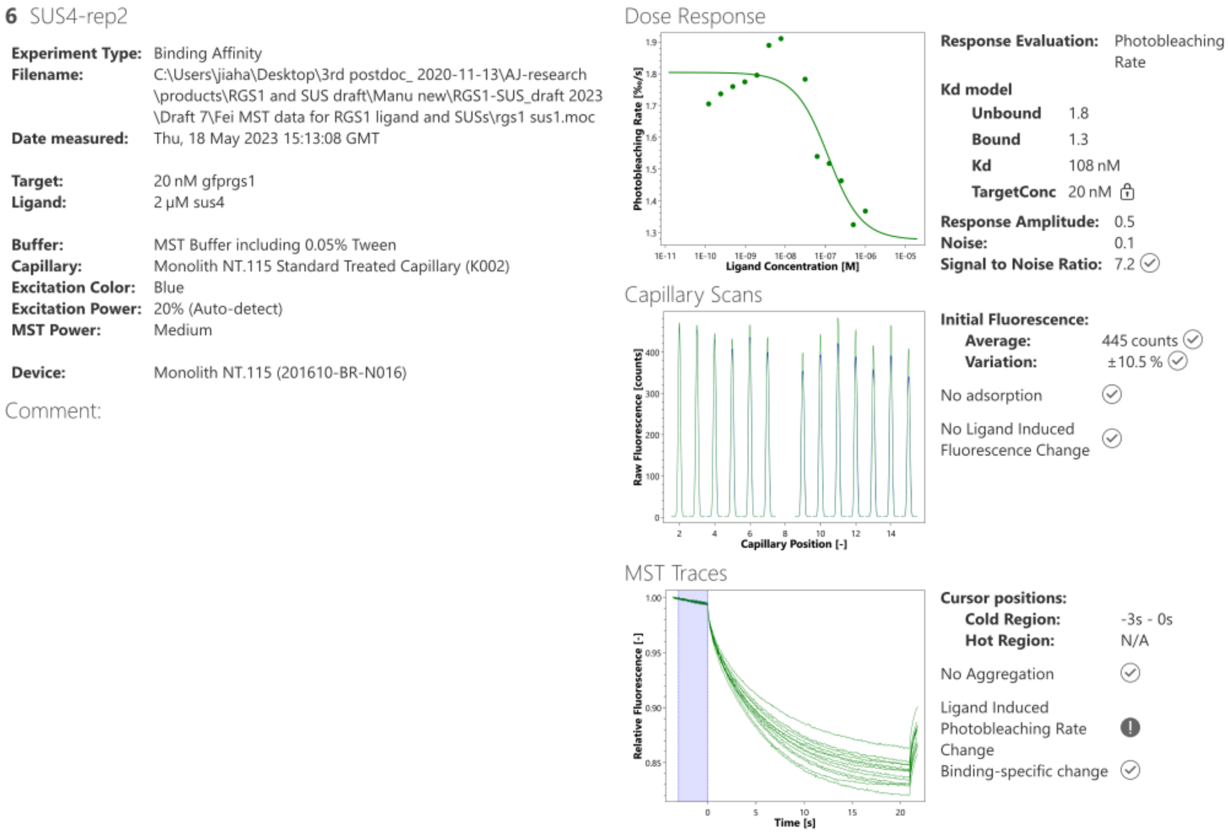


**Figure 8 (continued).**

**sfGFP-RGS1 and SUS4 binding profile in MST assay**

Microscale thermophoresis (MST) binding analysis of sfGFP-RGS1 (ΔC-tail) with His-SUS4. Serial dilutions of His-SUS4 were titrated against a fixed concentration of sfGFP-RGS1 (ΔC-tail). Changes in thermophoretic mobility were measured to generate the binding profile, and the dissociation constant ($K_d$) was determined by fitting the data to a standard binding model.

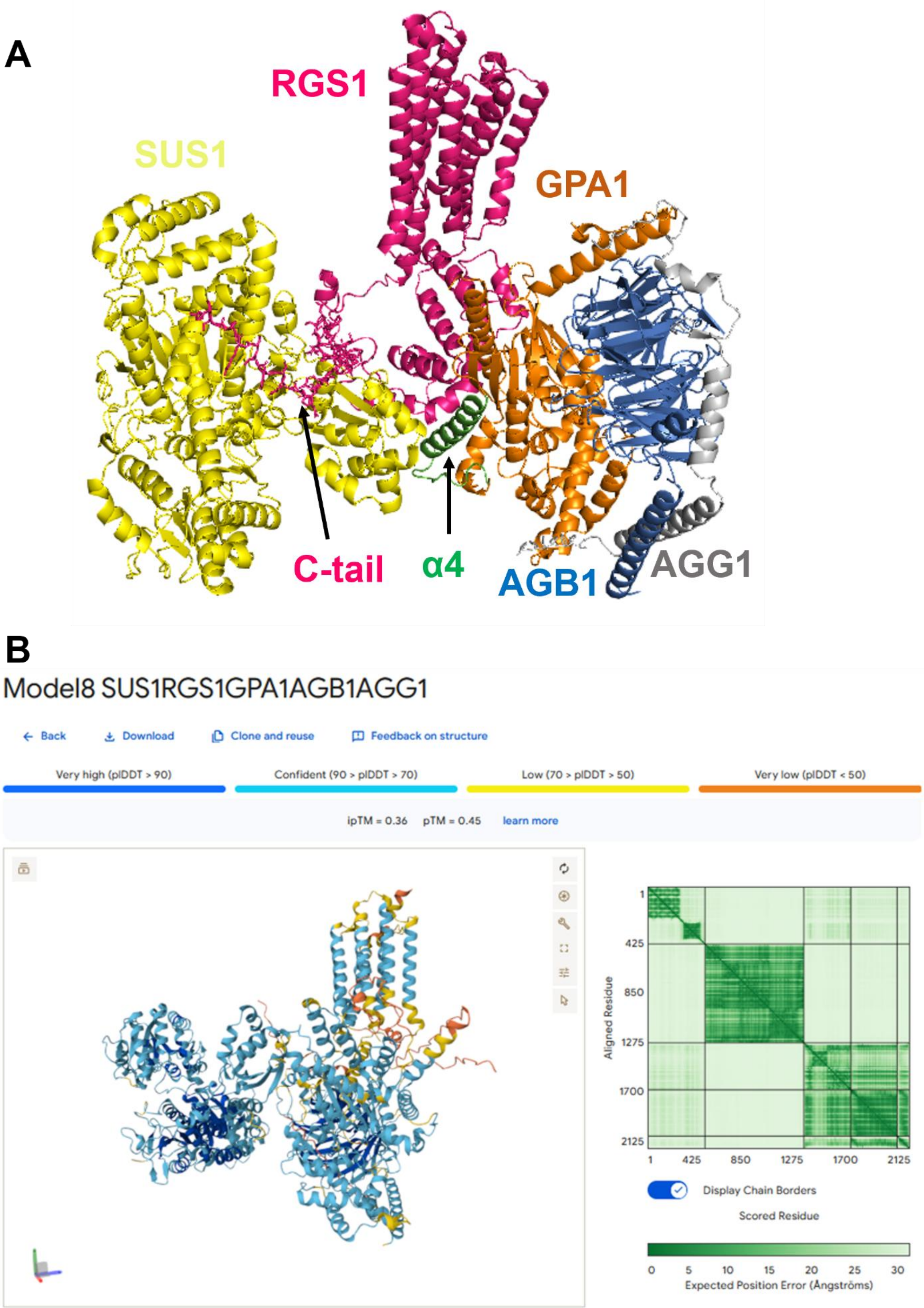
A
RGS1
SUS1
GPA1
C-tail
α4
AGB1
AGG1
B
Model8 SUS1RGS1GPA1AGB1AGG1
Back
Download
Clone and reuse
Feedback on structure
Very high (pIDDT > 90)
Confident (90 > pIDDT > 70)
Low (70 > pIDDT > 50)
Very low (pIDDT < 50)
ipTM = 0.36 pTM = 0.45 learn more
Aligned Residue
1
425
850
1275
1700
2125
Display Chain Borders
Scored Residue
0 5 10 15 20 25 30
Expected Position Error (Ångströms)

**Figure S9.**

**A. AlphaFold3-predicted structure of the SUS1–RGS1–G protein complex.**

The complex was modeled using AlphaFold3 (https://alphafoldserver.com/). RGS1 (pink), GPA1 (orange), AGB1 (blue), AGG1 (gray), and SUS1 (yellow) are shown. The C-terminal tail of RGS1 (S428–G459) forms multiple contact interfaces with SUS1, while the α4 helix (Green) of GPA1 also contributes to SUS1 binding.

**B. pLDDT confidence scores of AlphaFold3-predicted structure of the SUS1–RGS1–G protein complex.**

High scores indicate high model confidence for the corresponding residues.

**Top enriched GO terms among UDPG-upregulated genes in Col-0 (n = 848).**

| Gene Set Name(NO. Genes) | Description | Category | NO. Genes in Overlap (k) | p value | FDR |
|---|---|---|---|---|---|
| RESPONSE_TO_CHEMICAL_STIMULUS(3953) | GO:0042221 response to chemical stimulus, GOslim:biological_process | GO_BP | 287 | 4.89e-48 | 5.24e-44 |
| RESPONSE_TO_CHITIN(421) | GO:0010200 response to chitin, GOslim:biological_process | GO_BP | 81 | 3.79e-37 | 2.03e-33 |
| RESPONSE_TO_ORGANIC_SUBSTANCE(2739) | GO:0010033 response to organic substance, GOslim:biological_process | GO_BP | 204 | 6.56e-34 | 1.76e-30 |
| RESPONSE_TO_STIMULUS(6222) | GO:0050896 response to stimulus, GOslim:biological_process | GO_BP | 343 | 5.36e-34 | 1.76e-30 |
| RESPONSE_TO_WOUNDING(340) | GO:0009611 response to wounding, GOslim:biological_process | GO_BP | 69 | 5.53e-33 | 1.19e-29 |
| RESPONSE_TO_EXTERNAL_STIMULUS(1047) | GO:0009605 response to external stimulus, GOslim:biological_process | GO_BP | 111 | 2.46e-29 | 4.39e-26 |
| RESPONSE_TO_STRESS(4037) | GO:0006950 response to stress, GOslim:biological_process | GO_BP | 247 | 5.49e-29 | 8.41e-26 |
| RESPONSE_TO_ENDOGENOUS_STIMULUS(1604) | GO:0009719 response to endogenous stimulus, GOslim:biological_process | GO_BP | 130 | 3.91e-24 | 5.24e-21 |
| RESPONSE_TO_CARBOHYDRATE_STIMULUS(811) | GO:0009743 response to carbohydrate stimulus, GOslim:biological_process | GO_BP | 87 | 2.24e-23 | 2.67e-20 |
| RESPONSE_TO_JASMONIC_ACID_STIMULUS(471) | GO:0009753 response to jasmonic acid stimulus, GOslim:biological_process | GO_BP | 64 | 3.65e-22 | 3.91e-19 |
| JASMONIC_ACID_METABOLIC_PROCESS(159) | GO:0009694 jasmonic acid metabolic process, GOslim:biological_process | GO_BP | 36 | 4.25e-19 | 4.14e-16 |
| OXYLIPIN_METABOLIC_PROCESS(167) | GO:0031407 oxylipin metabolic process, GOslim:biological_process | GO_BP | 36 | 1.68e-18 | 1.5e-15 |
| RESPONSE_TO_FUNGUS(497) | GO:0009620 response to fungus, GOslim:biological_process | GO_BP | 59 | 5.41e-18 | 4.46e-15 |
| JASMONIC_ACID_BIOSYNTHETIC_PROCESS(135) | GO:0009695 jasmonic acid biosynthetic process, GOslim:biological_process | GO_BP | 31 | 8.82e-17 | 6.75e-14 |
| CELLULAR_RESPONSE_TO_STIMULUS(2302) | GO:0051716 cellular response to stimulus, GOslim:biological_process | GO_BP | 143 | 1.19e-16 | 8.53e-14 |
| RESPONSE_TO_HEAT(303) | GO:0009408 response to heat, GOslim:biological_process | GO_BP | 44 | 1.5e-16 | 1.01e-13 |
| OXYLIPIN_BIOSYNTHETIC_PROCESS(141) | GO:0031408 oxylipin biosynthetic process, GOslim:biological_process | GO_BP | 31 | 2.52e-16 | 1.59e-13 |
| DEFENSE_RESPONSE(1644) | GO:0006952 defense response, GOslim:biological_process | GO_BP | 113 | 5.04e-16 | 3e-13 |
| CELLULAR_RESPONSE_TO_ENDOGENOUS_STIMULUS(804) | GO:0071495 cellular response to endogenous stimulus, GOslim:biological_process | GO_BP | 72 | 1.21e-15 | 6.82e-13 |
| RESPONSE_TO_HORMONE_STIMULUS(1364) | GO:0009725 response to hormone stimulus, GOslim:biological_process | GO_BP | 99 | 1.7e-15 | 9.13e-13 |
| RESPONSE_TO_MECHANICAL_STIMULUS(63) | GO:0009612 response to mechanical stimulus, GOslim:biological_process | GO_BP | 21 | 1.83e-14 | 9.34e-12 |
| CELLULAR_RESPONSE_TO_STRESS(1431) | GO:0033554 cellular response to stress, GOslim:biological_process | GO_BP | 99 | 2.82e-14 | 1.37e-11 |
| RESPONSE_TO_ORGANIC_CYCLIC_COMPOUND(148) | GO:0014070 response to organic cyclic compound, GOslim:biological_process | GO_BP | 29 | 3.09e-14 | 1.38e-11 |
| RESPONSE_TO_CYCLOPENTENONE(148) | GO:0010583 response to cyclopentenone, GOslim:biological_process | GO_BP | 29 | 3.09e-14 | 1.38e-11 |
| RESPONSE_TO_WATER_STIMULUS(421) | GO:0009415 response to water stimulus, GOslim:biological_process | GO_BP | 47 | 1.09e-13 | 4.66e-11 |
| CELLULAR_RESPONSE_TO_JASMONIC_ACID_STIMULUS(282) | GO:0071395 cellular response to jasmonic acid stimulus, GOslim:biological_process | GO_BP | 38 | 1.47e-13 | 5.84e-11 |
| JASMONIC_ACID_MEDIATED_SIGNALING_PATHWAY(282) | GO:0009867 jasmonic acid mediated signaling pathway, GOslim:biological_process | GO_BP | 38 | 1.47e-13 | 5.84e-11 |
| RESPONSE_TO_OXIDATIVE_STRESS(586) | GO:0006979 response to oxidative stress, GOslim:biological_process | GO_BP | 56 | 1.81e-13 | 6.94e-11 |
| INNATE_IMMUNE_RESPONSE(926) | GO:0045087 innate immune response, GOslim:biological_process | GO_BP | 73 | 2.93e-13 | 1.08e-10 |

### Figure S10

Gene ontology analysis of the 848 UDPG upregulated genes in Col-0.

See methods for details in RNA-seq data analysis.

**Top enriched GO terms among UDPG-downregulated genes in Col-0 (n = 362)**

| Gene Set Name(NO. Genes) | Description | Category | NO. Genes in Overlap (k) | p value | FDR |
|---|---|---|---|---|---|
| CYTOKINESIS_BY_CELL_PLATE_FORMATION(204) | GO:0000911 cytokinesis by cell plate formation, GOslim:biological_process | GO_BP | 25 | 1.64e-16 | 9.99e-13 |
| CELL_CYCLE_CYTOKINESIS(208) | GO:0033205 cell cycle cytokinesis, GOslim:biological_process | GO_BP | 25 | 2.47e-16 | 9.99e-13 |
| CYTOKINESIS(235) | GO:0000910 cytokinesis, GOslim:biological_process | GO_BP | 25 | 3.27e-15 | 8.8e-12 |
| REGULATION_OF_CELL_CYCLE(296) | GO:0051726 regulation of cell cycle, GOslim:biological_process | GO_BP | 24 | 2.7e-12 | 5.45e-09 |
| CELL_CYCLE(802) | GO:0007049 cell cycle, GOslim:biological_process | GO_BP | 38 | 8.23e-12 | 7.44e-09 |
| REGULATION_OF_DNA_REPLICATION(135) | GO:0006275 regulation of DNA replication, GOslim:biological_process | GO_BP | 17 | 8.24e-12 | 7.44e-09 |
| SPINDLE_ASSEMBLY(45) | GO:0051225 spindle assembly, GOslim:biological_process | GO_BP | 12 | 4.83e-12 | 7.44e-09 |
| MICROTUBULE-BASED_PROCESS(313) | GO:0007017 microtubule-based process, GOslim:biological_process | GO_BP | 24 | 8.08e-12 | 7.44e-09 |
| DEFENSE_RESPONSE(1644) | GO:0006952 defense response, GOslim:biological_process | GO_BP | 57 | 8.29e-12 | 7.44e-09 |
| SPINDLE_ORGANIZATION(48) | GO:0007051 spindle organization, GOslim:biological_process | GO_BP | 12 | 9.24e-12 | 7.46e-09 |
| CELL_DIVISION(362) | GO:0051301 cell division, GOslim:biological_process | GO_BP | 25 | 2.44e-11 | 1.78e-08 |
| MICROTUBULE_CYTOSKELETON_ORGANIZATION(246) | GO:0000226 microtubule cytoskeleton organization, GOslim:biological_process | GO_BP | 21 | 2.64e-11 | 1.78e-08 |
| ORGANELLE_ASSEMBLY(57) | GO:0070925 organelle assembly, GOslim:biological_process | GO_BP | 12 | 5.27e-11 | 3.27e-08 |
| AROMATIC_COMPOUND_BIOSYNTHETIC_PROCESS(679) | GO:0019438 aromatic compound biosynthetic process, GOslim:biological_process | GO_BP | 31 | 1.78e-09 | 1.03e-06 |
| REGULATION_OF_DNA_METABOLIC_PROCESS(198) | GO:0051052 regulation of DNA metabolic process, GOslim:biological_process | GO_BP | 17 | 1.95e-09 | 1.05e-06 |

### Figure S11

Gene ontology analysis of the 362 UDPG down-regulated genes in Col-0.

See methods for details in RNA-seq data analysis.

## A

**Top enriched GO terms for UDPG-upregulated genes in *rgs1-2* vs. Col-0 (n = 124).**

| Gene Set Name(NO. Genes) | Description | Category | NO. Genes in Overlap (k) | p value | FDR |
|---|---|---|---|---|---|
| RESPONSE_TO_DISACCHARIDE_STIMULUS(212) | GO:0034285 response to disaccharide stimulus, GOslim:biological_process | GO_BP | 9 | 5.04e-07 | 8.24e-4 |
| RESPONSE_TO_FRUCTOSE_STIMULUS(144) | GO:0009750 response to fructose stimulus, GOslim:biological_process | GO_BP | 8 | 3.2e-07 | 8.24e-4 |
| RESPONSE_TO_SUCROSE_STIMULUS(209) | GO:0009744 response to sucrose stimulus, GOslim:biological_process | GO_BP | 9 | 4.49e-07 | 8.24e-4 |
| RESPONSE_TO_HEXOSE_STIMULUS(170) | GO:0009746 response to hexose stimulus, GOslim:biological_process | GO_BP | 8 | 1.06e-06 | 1.04e-3 |
| RESPONSE_TO_MONOSACCHARIDE_STIMULUS (170) | GO:0034284 response to monosaccharide stimulus, GOslim:biological_process | GO_BP | 8 | 1.06e-06 | 1.04e-3 |
| RESPONSE_TO_CHEMICAL_STIMULUS(3953) | GO:0042221 response to chemical stimulus, GOslim:biological_process | GO_BP | 35 | 1.97e-05 | 0.0161 |
| RESPONSE_TO_ORGANIC_SUBSTANCE(2739) | GO:0010033 response to organic substance, GOslim:biological_process | GO_BP | 27 | 3.79e-05 | 0.0266 |

## B

**Top enriched GO terms for UDPG-downregulated genes in *rgs1-2* vs. Col-0 (n = 571)**

| Gene Set Name(NO. Genes) | Description | Category | NO. Genes in Overlap (k) | p value | FDR |
|---|---|---|---|---|---|
| PHOTOSYNTHESIS(436) | GO:0015979 photosynthesis, GOslim:biological_process | GO_BP | 57 | 2.25e-27 | 2.1e-23 |
| PHOTOSYNTHESIS,_LIGHT_REACTION(333) | GO:0019684 photosynthesis, light reaction, GOslim:biological_process | GO_BP | 46 | 3.76e-23 | 1.75e-19 |
| DNA-DEPENDENT_TRANSCRIPTION,_ELONGATION(133) | GO:0006354 DNA-dependent transcription, elongation, GOslim:biological_process | GO_BP | 32 | 9.45e-23 | 2.93e-19 |
| GENERATION_OF_PRECURSOR_METABOLITES_AND_ENERGY(730) | GO:0006091 generation of precursor metabolites and energy, GOslim:biological_process | GO_BP | 60 | 2.34e-19 | 5.45e-16 |
| TRANSCRIPTION,_DNA-DEPENDENT(2616) | GO:0006351 transcription, DNA-dependent, GOslim:biological_process | GO_BP | 98 | 1.5e-09 | 2.17e-06 |
| RNA_BIOSYNTHETIC_PROCESS(2619) | GO:0032774 RNA biosynthetic process, GOslim:biological_process | GO_BP | 98 | 1.59e-09 | 2.17e-06 |
| RESPONSE_TO_CHEMICAL_STIMULUS(3953) | GO:0042221 response to chemical stimulus, GOslim:biological_process | GO_BP | 132 | 1.63e-09 | 2.17e-06 |
| ELECTRON_TRANSPORT_CHAIN(133) | GO:0022900 electron transport chain, GOslim:biological_process | GO_BP | 16 | 3.87e-08 | 3.6e-05 |
| PHOTOSYNTHETIC_ELECTRON_TRANSPORT_CHAIN(80) | GO:0009767 photosynthetic electron transport chain, GOslim:biological_process | GO_BP | 13 | 3.13e-08 | 3.6e-05 |
| SULFUR_COMPOUND_METABOLIC_PROCESS(682) | GO:0006790 sulfur compound metabolic process, GOslim:biological_process | GO_BP | 38 | 3.54e-08 | 3.6e-05 |
| RESPONSE_TO_ORGANIC_SUBSTANCE(2739) | GO:0010033 response to organic substance, GOslim:biological_process | GO_BP | 95 | 1.04e-07 | 8.04e-05 |
| RESPONSE_TO_STIMULUS(6222) | GO:0050896 response to stimulus, GOslim:biological_process | GO_BP | 178 | 9.71e-08 | 8.04e-05 |
| RESPONSE_TO_ENDOGENOUS_STIMULUS(1604) | GO:0009719 response to endogenous stimulus, GOslim:biological_process | GO_BP | 64 | 1.86e-07 | 1.33e-4 |

**Figure S12**

Gene ontology analysis of the up- and down-regulated genes in *rgs1-2* vs. Col-0.

See methods for details in RNA-seq data analysis.

### Top enriched GO terms for genes co-regulated by UDPG and RGS1 (n = 247)

| Gene Set Name(NO. Genes) | Description | Category | NO. Genes in Overlap (k) | p value | FDR |
|---|---|---|---|---|---|
| RESPONSE_TO_CHEMICAL_STIMULUS(3953) | GO:0042221 response to chemical stimulus, GOslim:biological_process | GO_BP | 81 | 2.70e-14 | 4.09e-11 |
| RESPONSE_TO_ORGANIC_SUBSTANCE(2739) | GO:0010033 response to organic substance, GOslim:biological_process | GO_BP | 57 | 3.42e-10 | 1.56e-07 |
| RESPONSE_TO_CHITIN(421) | GO:0010200 response to chitin, GOslim:biological_process | GO_BP | 20 | 1.58e-09 | 6.36e-07 |
| RESPONSE_TO_STIMULUS(6222) | GO:0050896 response to stimulus, GOslim:biological_process | GO_BP | 95 | 1.77e-09 | 6.70e-07 |
| RESPONSE_TO_WOUNDING(340) | GO:0009611 response to wounding, GOslim:biological_process | GO_BP | 18 | 2.14e-09 | 7.89e-07 |
| ETHYLENE_BIOSYNTHETIC_PROCESS(131) | GO:0009693 ethylene biosynthetic process, GOslim:biological_process | GO_BP | 12 | 3.92e-09 | 1.22e-06 |
| ETHYLENE_METABOLIC_PROCESS(131) | GO:0009692 ethylene metabolic process, GOslim:biological_process | GO_BP | 12 | 3.92e-09 | 1.22e-06 |
| ALKENE_BIOSYNTHETIC_PROCESS(133) | GO:0043450 alkene biosynthetic process, GOslim:biological_process | GO_BP | 12 | 4.60e-09 | 1.36e-06 |
| CELLULAR_ALKENE_METABOLIC_PROCESS(133) | GO:0043449 cellular alkene metabolic process, GOslim:biological_process | GO_BP | 12 | 4.60e-09 | 1.36e-06 |
| RESPONSE_TO_ENDOGENOUS_STIMULUS(1604) | GO:0009719 response to endogenous stimulus, GOslim:biological_process | GO_BP | 36 | 1.81e-07 | 3.13e-05 |
| RESPONSE_TO_EXTERNAL_STIMULUS(1047) | GO:0009605 response to external stimulus, GOslim:biological_process | GO_BP | 27 | 5.67e-07 | 8.99e-05 |
| METHIONINE_METABOLIC_PROCESS(236) | GO:0006555 methionine metabolic process, GOslim:biological_process | GO_BP | 12 | 1.62e-06 | 2.41e-04 |
| RESPONSE_TO_CARBOHYDRATE_STIMULUS(811) | GO:0009743 response to carbohydrate stimulus, GOslim:biological_process | GO_BP | 22 | 3.06e-06 | 4.30e-04 |
| CELLULAR_MODIFIED_AMINO_ACID_METABOLIC_PROCESS(708) | GO:0006575 cellular modified amino acid metabolic process, GOslim:biological_process | GO_BP | 20 | 4.90e-06 | 6.69e-04 |
| RESPONSE_TO_JASMONIC_ACID_STIMULUS(471) | GO:0009753 response to jasmonic acid stimulus, GOslim:biological_process | GO_BP | 16 | 4.98e-06 | 6.73e-04 |
| ASPARTATE_FAMILY_AMINO_ACID_METABOLIC_PROCESS(277) | GO:0009066 aspartate family amino acid metabolic process, GOslim:biological_process | GO_BP | 12 | 7.80e-06 | 1.00e-03 |
| RESPONSE_TO_FUNGUS(497) | GO:0009620 response to fungus, GOslim:biological_process | GO_BP | 16 | 9.54e-06 | 1.16e-03 |
| RESPONSE_TO_STRESS(4037) | GO:0006950 response to stress, GOslim:biological_process | GO_BP | 59 | 2.65e-05 | 2.76e-03 |
| JASMONIC_ACID_BIOSYNTHETIC_PROCESS(135) | GO:0009695 jasmonic acid biosynthetic process, GOslim:biological_process | GO_BP | 8 | 3.25e-05 | 3.26e-03 |
| OXYLIPIN_BIOSYNTHETIC_PROCESS(141) | GO:0031408 oxylipin biosynthetic process, GOslim:biological_process | GO_BP | 8 | 4.36e-05 | 4.19e-03 |
| RESPONSE_TO_OXIDATIVE_STRESS(586) | GO:0006979 response to oxidative stress, GOslim:biological_process | GO_BP | 16 | 6.55e-05 | 5.92e-03 |
| CELLULAR_MODIFIED_AMINO_ACID_BIOSYNTHETIC_PROCESS(532) | GO:0042398 cellular modified amino acid biosynthetic process, GOslim:biological_process | GO_BP | 15 | 7.84e-05 | 6.70e-03 |
| RESPONSE_TO_BRASSINOSTEROID_STIMULUS(114) | GO:0009741 response to brassinosteroid stimulus, GOslim:biological_process | GO_BP | 7 | 8.23e-05 | 6.87e-03 |
| RESPONSE_TO_HEAT(303) | GO:0009408 response to heat, GOslim:biological_process | GO_BP | 11 | 8.70e-05 | 7.20e-03 |
| JASMONIC_ACID_METABOLIC_PROCESS(159) | GO:0009694 jasmonic acid metabolic process, GOslim:biological_process | GO_BP | 8 | 9.73e-05 | 7.85e-03 |
| OXYLIPIN_METABOLIC_PROCESS(167) | GO:0031407 oxylipin metabolic process, GOslim:biological_process | GO_BP | 8 | 1.35e-04 | 1.03e-02 |
| RESPONSE_TO_HORMONE_STIMULUS(1364) | GO:0009725 response to hormone stimulus, GOslim:biological_process | GO_BP | 26 | 1.51e-04 | 1.12e-02 |
| RESPONSE_TO_OTHER_ORGANISM(1411) | GO:0051707 response to other organism, GOslim:biological_process | GO_BP | 26 | 2.54e-04 | 1.76e-02 |
| SULFUR_AMINO_ACID_METABOLIC_PROCESS(465) | GO:0000096 sulfur amino acid metabolic process, GOslim:biological_process | GO_BP | 13 | 2.54e-04 | 1.76e-02 |
| DEFENSE_RESPONSE_BY_CALLOSE_DEPOSITION(62) | GO:0052542 defense response by callose deposition, GOslim:biological_process | GO_BP | 5 | 2.69e-04 | 1.84e-02 |
| RESPONSE_TO_MECHANICAL_STIMULUS(63) | GO:0009612 response to mechanical stimulus, GOslim:biological_process | GO_BP | 5 | 2.89e-04 | 1.95e-02 |
| RESPONSE_TO_BIOTIC_STIMULUS(1675) | GO:0009607 response to biotic stimulus, GOslim:biological_process | GO_BP | 28 | 6.72e-04 | 4.02e-02 |
| FATTY_ACID_METABOLIC_PROCESS(587) | GO:0006631 fatty acid metabolic process, GOslim:biological_process | GO_BP | 14 | 6.95e-04 | 4.09e-02 |
| INTRACELLULAR_SIGNAL_TRANSDUCTION(222) | GO:0035556 intracellular signal transduction, GOslim:biological_process | GO_BP | 8 | 8.36e-04 | 4.81e-02 |

### Figure S13

Gene ontology analysis of genes co-regulated by UDPG and RGS1.

See methods for details in RNA-seq data analysis.
